# Identifying a next-generation antimalarial trioxolane in a landscape of artemisinin partial resistance

**DOI:** 10.1101/2025.01.02.631109

**Authors:** Matthew T. Klope, Poulami Talukder, Brian R. Blank, Sevil Chelebieva, Jun Chen, Shaun D. Fontaine, Ryan L. Gonciarz, Priyadarshini Jaishankar, Grace J. Lee, Jennifer Legac, Vineet Mathur, Avani Narayan, Martin Okitwi, Stephen Orena, Nicholas S. Settineri, Juan Tapia, Yoweri Taremwa, Patrick K. Tumwebaze, Aswathy Vinod, Jeremy N. Burrows, Philip J. Rosenthal, Roland A. Cooper, Adam R. Renslo

## Abstract

For over two decades, artemisinin-based combination therapy (ACT) has been the standard of care for the treatment of uncomplicated falciparum malaria. However, artemisinin partial resistance (ART-R) is now prevalent in Southeast Asia and has emerged in eastern Africa, threatening ACT efficacy. Mechanistically, ART-R results from an endocytosis defect that limits concentrations of host-derived free heme in the parasite digestive vacuole, allowing early ring-stage parasites to survive exposure to the artemisinin component of ACT. The artemisinin-inspired 1,2,4-trioxolane artefenomel exhibits an extended pharmacokinetic exposure profile that predicts efficacy against ART-R parasites. Unfortunately, the development of artefenomel was halted recently after almost a decade of clinical trials. Herein, we describe the discovery of RLA-4735 and its single-enantiomer form RLA-5764, next-generation antimalarial trioxolanes that exhibit excellent in vitro potency against *Plasmodium falciparum* and single-exposure efficacy in a murine *P. berghei* model, thus retaining many of the favorable pharmacokinetic and pharmacodynamic properties of artefenomel while markedly improving solubility and development potential. In *P. falciparum* samples collected from patients in Uganda in 2019 and 2023, ex vivo ring-stage survival assays revealed the emergence of the ART-R phenotype over this timeframe, and furthermore demonstrated markedly superior activity of artefenomel and RLA-4735 as compared to dihydroartemisinin (the active metabolite of artemisinin components of ACTs) against ART-R parasites. Overall, our findings suggest a role for next-generation trioxolanes in addressing ART-R, and present a potent new, artefenomel-adjacent chemotype with good potential to deliver new development candidates.

**Summary Sentence:** Klope et. al. described the discovery and in vivo characterization of antimalarial endoperoxides effective against artemisinin-resistant parasites as potential development candidates for uncomplicated, blood-stage malaria.

## Introduction

For two decades artemisinin combination therapy (ACT) has been the standard of care to treat uncomplicated falciparum malaria, likely saving millions of lives (*1*). The unprecedented molecular pharmacology of artemisinins involving initial activation by heme iron (*2–4*), followed by promiscuous labelling of the parasite proteome (*5–7*), results in potent killing of all asexual erythrocytic stages of *Plasmodium falciparum* (*8*). Inspired by these agents and their clinical utility, researchers have long sought to identify synthetic endoperoxides that could be manufactured at low cost, and whose pharmacokinetic properties could be optimized to overcome the rapid clearance profile of dihydroartemisinin (DHA), the active metabolite of all clinically used artemisinins (Fig.1). Among the various endoperoxide chemotypes explored (*9*), only the 1,2,4- trioxolanes arterolane (*10*) and artefenomel (**1**, Fig.1) have so far been studied in clinical trials (*11, 12*). While arterolane is approved in India and some African countries as a combination with piperaquine, its pharmacokinetics are only marginally improved over those of DHA, and the drug has seen limited clinical use. Much higher hopes had been invested in artefenomel (**1**), which combines an extended in vivo exposure profile and pre-clinically produces single-exposure cures, an important (*13, 14*) benchmark in antimalarial discovery, although whether single-exposure cure should also be a clinical benchmark remains controversial (*15*). However, after more than a decade of clinical study (*16–18*), the development of **1** was recently discontinued. The failure of **1**, combined with either piperaquine or ferroquine, to achieve target efficacy in clinical studies following a single dose was disappointing. The clinical challenge of achieving single-exposure cure in these studies was confounded by the high doses of **1** and partner drug required for efficacy (totaling nearly 2 g in some studies) and the profound formulation difficulties encountered with **1**, which required administration as an oral suspension in clinical studies.

**Fig. 1.**
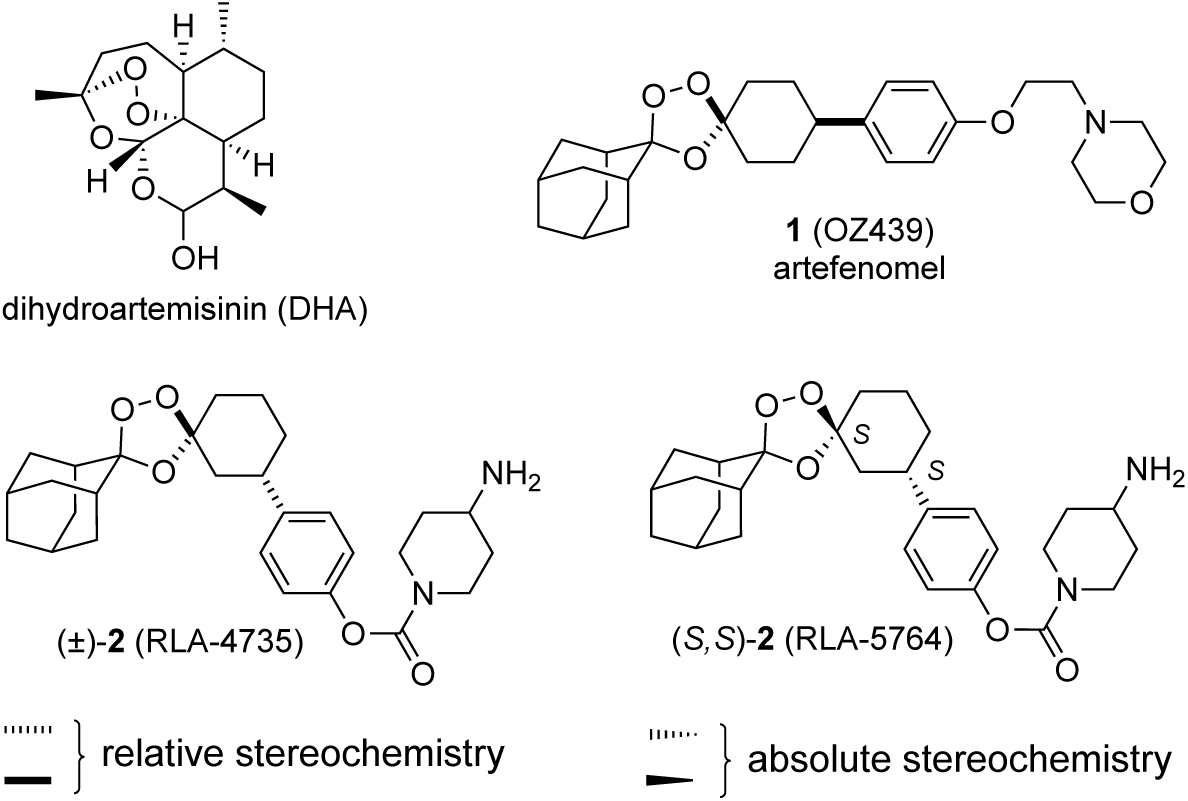
Structures of dihydroartemisinin, the active form of all artemisinins used clinically, the *cis*- 4’ trioxolane artefenomel (**1**), and *trans*-3’ trioxolane analogs (±)-**2** (RLA-4735) and (*S,S*)-**2** (RLA- 5764). Drawing conventions denoting relative or absolute configurations of stereocenters are indicated.

Given the central role of artemisinins in ACT, the emergence of artemisinin partial resistance (ART-R) in Southeast Asia (*19*) and its more recent emergence in eastern Africa (*20, 21*) is of considerable concern (*22*). Clinically, the ART-R phenotype manifests as a delayed parasite clearance that can be accompanied by treatment failures, in particular with coexistent resistance to ACT partner drugs. Befitting the remarkable molecular pharmacology of artemisinins is an equally remarkable mechanism of drug resistance. Thus, ART-R is associated with destabilizing mutations (*19, 23*) in the propeller domain of the *Kelch13* (K13) protein, causing an endocytosis defect that reduces parasite uptake of host hemoglobin, thus limiting the parasite pool of artemisinin-activating free heme iron, particularly in early rings (*24, 25*). This mechanism is supported by recent studies defining the role of the K13 protein in parasite endocytosis (*24, 26*), and by earlier studies in which an ART-R phenotype was observed in parasites deficient in hemoglobin catabolism (*27, 28*). In summary, the relative resistance of K13 mutant rings to DHA, combined with the drug’s rapid in vivo clearance profile, allows K13 mutant *P. falciparum* to survive brief DHA exposure and progress through the erythrocytic cycle.

The unusual, ring-stage specific resistance of K13 mutant parasites presents interesting challenges and opportunities for the discovery of a next-generation agent. Among the challenges are that traditional in vitro growth inhibition assays do not reveal the reduced susceptibility of ART-R parasites to DHA. To reveal the ART-R phenotype, a ring-stage survival assay (RSA) was developed, in which synchronized ring-stage parasite subjected to a short pulse of DHA to mimic its pharmacokinetic profile (*29*). Although explicitly developed to assess DHA susceptibility, similar pulsed-exposure experiments have been used to model the PK exposure of **1**, and to predict that this longer-acting agent should be effective clinically against ART-R parasites (*30, 31*). While this prediction was not wholly borne out in clinical studies of **1** performed in ART-R endemic areas (*18*), the suboptimal dosing regimens used and high rates of emesis suggest that target exposure levels of **1** and partner drug were not achieved, quite possibly leaving this key hypothesis untested.

With an understanding of the molecular mechanism underlying ART-R, it is now possible to consider a rational approach to the design of next-generation endoperoxide agents. On the one hand, the fact that K13 mutant rings possess a limiting pool of activating ferrous iron could argue for an agent with *enhanced* iron reactivity. However, such an approach would run counter to the need for good drug stability in the bloodstream of infected patients (*11*) and is further argued against by the reduced stability of endoperoxides at high levels of parasitemia (*32*). Instead, an agent with *enhanced* stability toward iron combined with a prolonged in vivo and intra-parasite exposure profile should allow the drug to persist through the resistant ring stages of ART-R parasites and then realize potent activity against more susceptible trophozoite and schizont stages. This activity would depend, of course, on the molecule retaining sufficient iron reactivity (*33, 34*) to produce the carbon-centered radical species (Fig. S1) that confer toxic effects in the parasite (*5–7, 35*). To this end, it is well known (*36*) that iron sensitivity of the dispiro-1,2,4-trioxolane pharmacophore (as found in **1**) is modulated by conformational dynamics of the cyclohexane ring, which in turn are determined by the nature and stereochemistry of the 4” side chain. Specifically, 1,3-diaxial interactions of the *cis*-4” substituent favor a ground-state conformer in which the peroxide bond is shielded by four neighboring axial C–H bonds and prevented from inner-sphere coordination with iron (Fig. 2 and Fig. S2). Thus, by employing a bulky 4” aryl side chain in **1**, Vennerstrom and co-colleagues achieved enhanced stability of the endoperoxide function, whilst retaining sufficient iron reactivity to exert potent antiparasitic effects (*11, 37*).

**Fig. 2.**
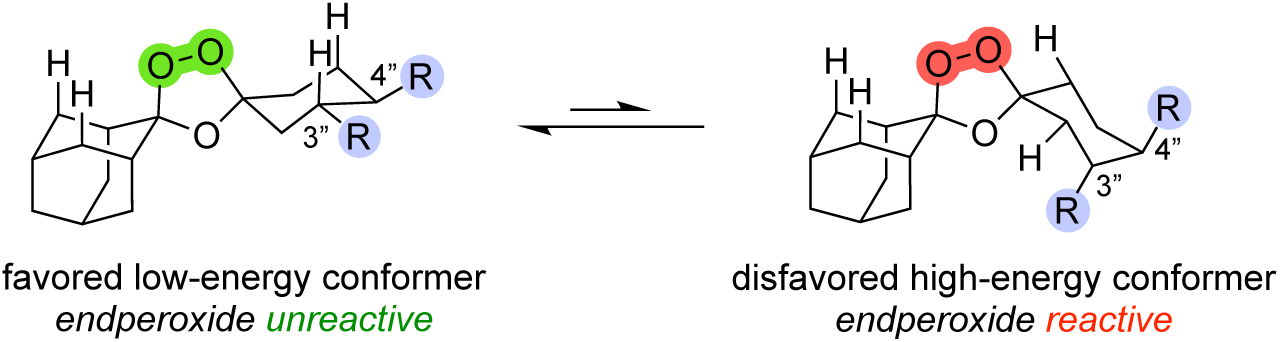
Iron reactivity of trioxolane antimalarials is determined by conformational equilibria of the cyclohexane ring. *Cis*-4” substitution as in **1** or *trans*-3” substitution as in **2** and related analogs discussed herein stabilize the endoperoxide bridge (in green and red highlights) in a pharmacologically optimal range for antimalarial effects.

To identify next-generation trioxolane agents with differentiated properties, our laboratory is exploring alternate means to modulate conformational dynamics in the dispiro-1,2,4-trioxolane pharmacophore. Thus, we have explored (*38–41*) novel analogs bearing *trans*-3” substitution, which conformational analysis (Fig. 2 and Fig. S2) suggests should stabilize the endoperoxide bond to a similar degree as traditional *cis*-4” substitution. Indeed, congeneric *trans*-3” analogs of arterolane and **1** exhibited promising levels of antiplasmodial activity, both in vitro and in vivo, confirming that *trans*-3” substitution modulates iron reactivity in a pharmacologically desirable regime (*39, 41*). Described herein are the results of our ongoing efforts in this vein, culminating in the discovery of **2** (Fig. 1), an advanced lead and potential trioxolane development candidate effective against ART-R parasites.

## Results

Medicines for Malaria Venture has detailed (*14, 42*) drug product profiles for new agents to be used in future antimalarial combination therapies, including Target Candidate Profile 1 (TCP-1), which emphasizes rapid reduction of parasite burden to minimal levels. At the outset of our effort, **1** remained the best hope for a new agent meeting the criteria of TCP-1, and thus **1** was a benchmark compound for in vitro and in vivo studies. Compound evaluation and progression was guided by three primary assays, 1) *in vitro* activity against the chloroquine-resistant W2 strain of *P. falciparum*, 2) *in vitro* human liver microsome (HLM) apparent intrinsic clearance, and 3) aqueous solubility in pH 7.4 PBS. We employed the *P. berghei* mouse model with a single oral dose of test compounds to initially assess the PD profiles of our more promising lead compounds. We predicted that new analogs that combined potent in vitro activity with good in vitro ADME properties and single-exposure efficacy in the *P. berghei* model (indicative of a prolonged exposure profile) should also prove effective against ART-R strains, as was suggested by preclinical time-kill studies of various trioxolanes (*30, 31*), including **1**. Subsequent testing of K13 mutant parasites using the RSA would be performed on the best of the analogs identified.

The move to *trans*-3” substitution of the trioxolane pharmacophore introduces two stereogenic centers, resulting in four possible diastereomers. However, the Griesbaum co-ozonolysis reaction used to prepare the trioxolane pharmacophore is an inherently stereoselective process, favoring *cis* (relative) stereochemistry in the case of 4”-substitution (*43*) and *trans* stereochemistry in the case of 3” substitution (*38*). Thus, we employed a short six-step route to prepare (±)-**2** and related analogs (Fig. 1 and Fig. S3) very similar to the four-step synthesis we recently reported (*41*) for the artefenomel regioisomer (±)-**3** (RLA-3107, Fig. 3). To access enantiopure forms such as (*R,R*)-**2** and (*S,S*)-**2** simply required the use of enantiopure cyclohexanone intermediates in the Griesbaum reaction (Fig. S4 and S5, respectively). Further discussion of the synthesis of *trans*-3”-aryl trioxolane analogs is provided as Supporting Text (Supporting Materials).

**Fig. 3.**
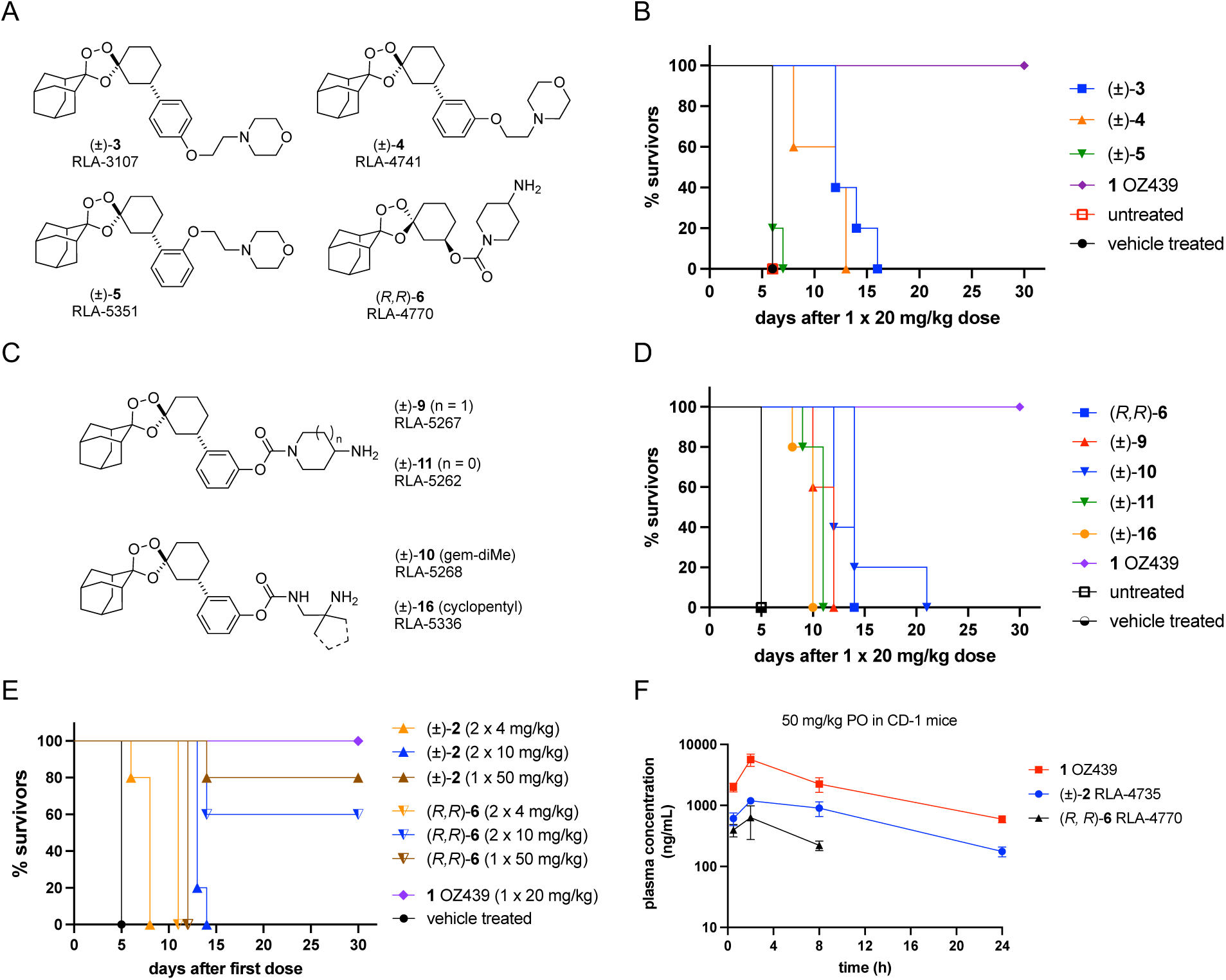
In vivo efficacy and pharmacokinetic profiles of *trans*-3’ trioxolane analogs as compared to the *cis*-4’ comparator artefenomel (**1**, OZ439). (**A**) Structures of regioisomeric *trans*-3’-aryl trioxolane analogs **3**-**5** and the *trans*-3’-carbamate exemplar (*R,R*)-**6**. (B) Kaplan-Meier survival curves comparing the efficacy of **3**-**5** in the *P. berghei* mouse infection model following a single oral dose of 20 mg/kg. (C) Structures of selected *trans*-3’-aryl carbamate analogs **9**-**11** and **16.** (D) Kaplan-Meier survival curves showing efficacy of **6**, **9**-**11**, and **16** in the *P. berghei* model (E) Kaplan-Meier curve for **1**, (±)-**2**, and **6** in the *P. berghei* mouse infection model when administered orally once daily at the indicated frequency and dose. (F) Plasma exposure curve for **1**, (±)-**2**, and (*R*,*R*)-**6** after a single oral dose of 50 mg/kg.

In our recent report (*41*), we found that artefenomel regioisomer **3** had improved HLM stability and aqueous solubility as compared to **1**, while producing single-exposure cures in the murine *P. berghei* model at 80 mg/kg. Following on these findings, we evaluated the closely related analogs **4** and **5**, which bear side chains regioisomeric with **3** (Fig. 3A) and could be prepared in four steps, as detailed in Supporting Information (Fig. S6-S8). We found that meta isomer **4** exhibited in vitro potency similar to that of **3**, and about 4-fold weaker than that of **1**, while ortho isomer **5** had potency about 8-fold weaker than that of **1** (Table 1). The HLM stability of **4** was inferior to that of **3** and roughly comparable to that of **1**, whereas **5** was rapidly metabolized in the HLM preparations (Table 1). The in vivo efficacy of **3**-**5** at a 20 mg/kg dose paralleled their HLM stabilities (suggesting similar murine hepatic extraction ratios), with compound **3** affording a greater survival benefit than **4**, and compound **5** showing no benefit over vehicle (Fig. 3B). By comparison, the artefenomel control was fully curative at a single 20 mg/kg dose, as previously reported by Vennerstrom and co-workers (*11*).

**Table 1.** Selected in vitro antiplasmodial activity and in vitro metabolism and solubility values for **1, 2** in racemic and single enantiomer forms, regioisomeric 3’-aryl substituted congeners **3**-**5** and 3’- carbamate **6**.

| compound | In vitro antiplasmodial or ADME assay data |  |  |  |
| --- | --- | --- | --- | --- |
|  | W2 *P. falc.* IC<sub>50</sub><br>(nM) | HLM CL<sub>int</sub><br>μL/min/mg | HLM t<sub>1/2</sub><br>(min) | solubility PBS pH 7.4<br>(μM) |
| **1** | 11 ± 0.5 | 72.1<sup>a</sup> | n.r.<sup>a</sup> | 0.181<sup>a</sup> |
| (±)-**2** | 6.1 ± 0.3 | 6.0<sup>b</sup> | 229<sup>b</sup> | 3.5–11.5 |
| (±)-**3** | 24 ± 0.6 | 21.6 | 65 | 24.8 |
| (±)-**4** | 29 ± 5.1 | 57.2 | 24 | n.d. |
| (±)-**5** | 53 ± 0.5 | 282 | 4.9 | n.d. |
| (*R,R*)-**2** | 3.8 ± 0.2 | <11.6 | >120 | 137–164 |
| (*S,S*)-**2** | 2.9 ± 0.2 | <11.6 | >120 | 78–80 |
| (*R,R*)-**6** | 1.7 ± 0.1<sup>c</sup> | 123<sup>b</sup> | 11.3<sup>b</sup> | 144.5<sup>b</sup> |

<sup>a</sup>as reported by MMV (46). <sup>b</sup>data generated via MMV. cfrom our previous report (40). IC<sub>50</sub> values are average of three replicates ± SEM. HLM; human liver microsome stability. Ranges reported when multiple measurements were performed. n.r. not reported; n.d. not determined.

Next, we conducted a larger survey of aryl ether and carbamate *trans*-3” side chains to identify analogs with improved properties (see Supplemental Text for a full discussion). We found that *carbamate* linked side chains bearing basic primary amines yielded analogs with in vitro potencies superior to those of ether-linked analogs like **3**-**5** (Fig. 3C and Fig. S10). Metabolic stability trends mirrored those observed for compounds **3**-**5**, with meta and para-substituted analogs more stable than ortho analogs. The best of the aryl carbamates, including compounds such as **2, 9**, **10**, **18**, and **21,** (Fig. 3C and S10) combined single-digit nM IC_50_ values with favorable aqueous solubility in the mid-μM regime, and HLM clearance values of <11 μL/min/mg, as compared to 72.1 μL/min/mg for **1** (Table 1 and Fig. S10). However, despite their favorable in vitro properties, meta carbamate analogs **9**, **10**, **11**, and **16** showed a median survival of between 9-12 days following a single 20 mg/kg dose (Fig. 3D), as compared to compound **3** that previously showed a median survival 11 days at the same dose (*41*). This lead us to focus further efforts on the more promising para carbamates, and in particular compound **2**.

To better understand PK/PD relationships in mice, we compared the in vivo efficacy and PK of compounds **1**, (±)-**2**, and (*R,R*)-**6**, the latter an optimized *trans*-3” heteroaliphatic carbamate described by our group previously (*40*). For the congeneric 4-aminopiperidines **2** and **6**, we compared a single 50 mg/kg dose with two daily doses of 10 mg/kg. We found for **2** that a single 50 mg/kg dose was more efficacious, producing cures in 4/5 mice. By contrast for (*R,R*)-**6**, two doses of 10 mg/kg was partially curative (3/5 mice) while the single larger dose produced no cures (Fig. 3E). This implied PK differences between the two compounds, since their in vitro potencies and ADME properties were similar. In a mouse PK experiment following a single 50 mg/kg oral dose, we found that compound **2** (like **1**) exhibited an extended exposure profile, with total plasma concentrations in excess of in vitro IC_50_ values 24 hrs after the single dose (Fig. 3F and Table S1). By contrast, (*R,R*)-**6** was undetectable at 24 hrs following the single dose, and exhibited an AUC that was only ∼20% that of **2** and just ∼6% that of **1** (Table S1). In summary, by comparing optimized *trans*-3” analogs bearing 4-aminopiperidine terminated side chains, we revealed the PK advantages of 3-aryl carbamates like **2** over congeneric 3-heteroalkyl carbamates like **6**. The enhanced iron stability of the 3-aryl carbamates, as discussed later, likely contribute to their superior PK profile.

Overall, our SAR studies nominated (±)-**2** as a candidate for more extensive ADME and PK profiling. Although the compound could not match the remarkable efficacy of **1** in mice, the overall similar plasma exposure profile combined with superior HLM stability suggested the potential for improved human PK and efficacy. Moreover, its improved aqueous solubility suggested **2** might avoid the formulation challenges that stymied the clinical development of **1** (*44, 45*). We next sought to compare the in vitro and in vivo properties of the enantiomerically pure forms (*R,R*)-**2** and (*S,S*)-**2**. The solution of a small molecule crystal structure of the form (*R,R*)-**2** (RLA-5763, Fig. 4A) as the hydrochloride salt confirmed the absolute stereochemistry of the material and validated our asymmetric synthetic routes to both enantiopure forms (Fig. S4 and S5). Evaluation of the enantiopure forms against laboratory-adapted clinical strains of *P. falciparum* bearing K13 mutations showed similar, low single-digit nM or high pM IC_50_ values (Table 2). While the growth inhibition assay, unlike RSAs, could not show the artemisinin partial resistance phenotype of these parasites, these studies did reveal the excellent potency of enantiopure forms of **2** in laboratory-adapted clinical isolates. With regard to ADME profile, the HLM clearance for (*R,R*)-**2** and (*S,S*)-**2** were both excellent, and interestingly, both enantiopure forms showed aqueous solubility ∼10-fold greater than that of the racemate (Table 1). This fortuitous observation was confirmed in repeated solubility studies of compound **2** as the formate salt in both racemic and enantiopure forms. Largely mirroring the efficacy of the racemate, both (*R,R*)-**2** and (*S,S*)-**2** exhibited fully curative efficacy (5/5 mice) in the *P. berghei* model when administered as a single oral dose of 50 mg/kg (Fig. 4B).

**Fig. 4.**
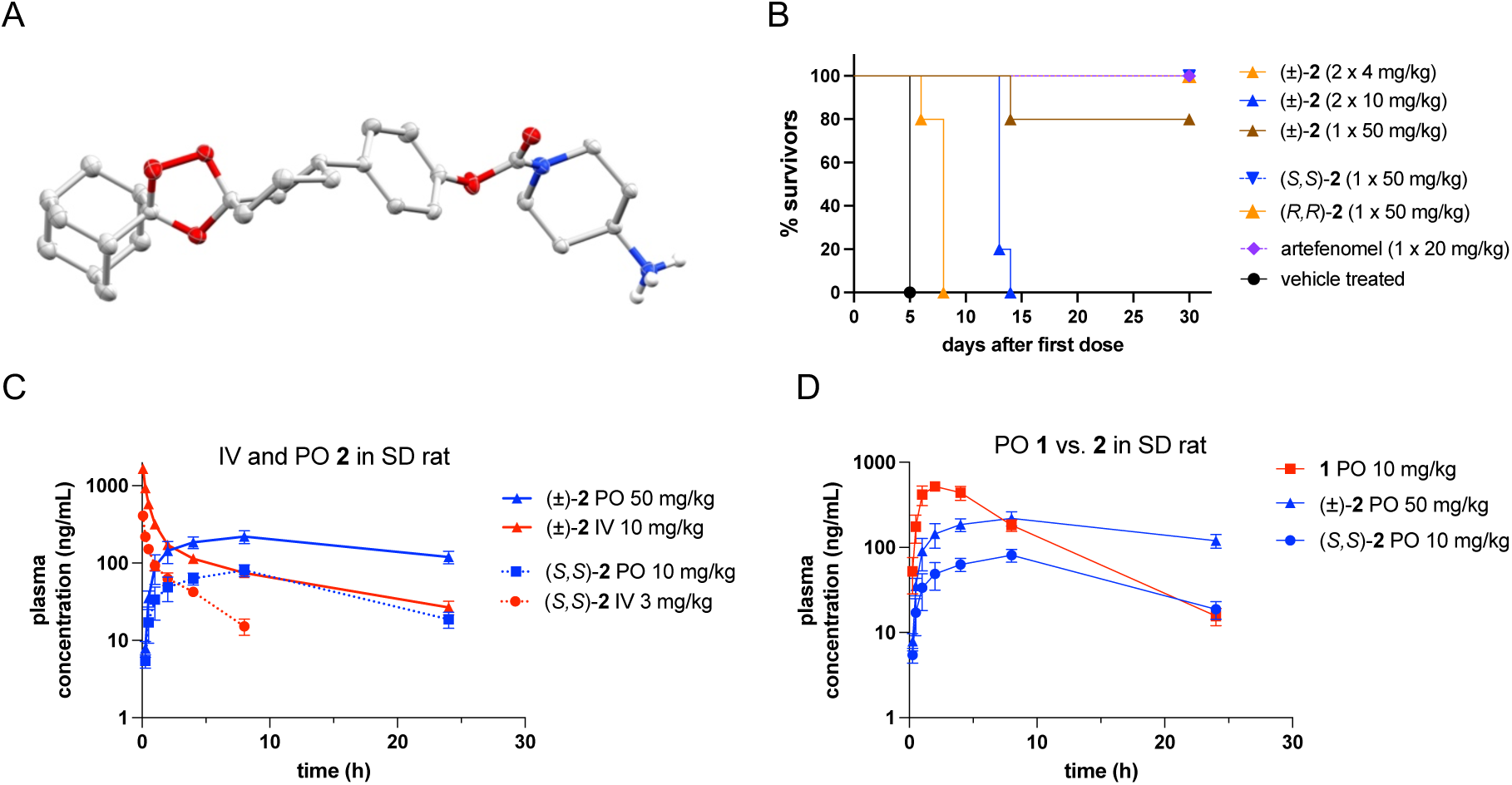
Confirmation of absolute configuration of (*R,R*)-**2** and in vivo PK/PD properties of **2** in racemic and enantiomerically pure forms. (A) Molecular structure of (*R,R*)-**2** (RLA-5763) from the crystal structure of the HCl salt (thermal ellipsoids drawn at the 30% probability level). Additional identical copies in the asymmetric unit, water molecules, chloride counterions, and carbon-bound hydrogens have been omitted for clarity. (B) Kaplan-Meier curves showing in vivo efficacy of (±)- **2**, (*S,S*)-**2**, and (*R,R*)-**2** in the *P. berghei* mouse malaria model at the indicated doses and dosing regimens. (C) Pharmacokinetic profile of (±)-**2** and (*S,S*)-**2** in SD rat, following oral and IV doses of 50 and 10 mg/kg, and 10 and 3 mg/kg, respectively. (D) Comparison of plasma exposure in SD rat, following oral dose of **1**, (±)-**2** or (*S,S*)-**2** at the indicated dose. PK Data for **1** is from (*11*) and was kindly provided by S. Charman (Monash Institutes of Pharmaceutical Sciences).

**Table 2.** *In vitro* antiplasmodial activity of (*R,R*)-**2** and (*S,S*)-**2** against lab-adapted Ugandan *P. falciparum* bearing K13 mutations. IC_50_ values determined in standard growth inhibition assay and not the ring stage assay.

| Compound | PBC-403<br>K13 A578S<br>IC<sub>50</sub> (nM) | BUS-25<br>K13 C469Y<br>IC<sub>50</sub> (nM) | PAT-25<br>K13 C469Y<br>IC<sub>50</sub> (nM) | PAT-26<br>K13 C469Y<br>IC<sub>50</sub> (nM) | BUS-41<br>K13 WT<br>IC<sub>50</sub> (nM) |
| --- | --- | --- | --- | --- | --- |
| (R,R)-**2** | 2.78 ± 1.46 | 1.34 ± 0.55 | 1.90 ± 1.02 | 0.50 ± 0.18 | 0.73 ± 0.37 |
| (S,S)-**2** | 1.56 ± 0.84 | 0.16 ± 0.03 | 0.76 ± 0.23 | 0.50 ± 0.21 | 0.29 ± 0.07 |
| DHA | 1.49 | 3.88 ± 1.04 | 0.67 | 1.84 ± 0.48 | 0.57 |

Having established similar in vitro and in vivo properties of enantiopure forms of **2**, we selected the form (*S,S*)-**2** (RLA-5764) for more extensive preclinical ADME and PK assessment (Table 3 and Table S2). Thus, (*S,S*)-**2** showed excellent stability in human and mouse plasma as well as in microsomes and hepatocytes from rat, dog, and human (t_1/2_ >240 min in all assays). Binding to plasma protein was exceptionally high (>99.9%) across three species, and to human hepatocytes and microsomes, similar to values reported for **1** (*46*). The blood to plasma ratio for (*S,S*)-**2** was 1.61 as compared to a value of 0.78 reported for **1.** Thermodynamic solubility (*S,S*)-**2** in fed-state simulated intestinal fluid was substantially higher (4525 μg/mL) than in the fasted state (2.3 μg/mL), suggesting potential drug-food interaction, a potential concern given the challenge of this issue in the development of **1**. Permeability in Caco-2 monolayers was performed using human plasma as the assay media, as has been recommended by Charman for compounds likely to have high membrane retention and low mass balance using purely aqueous transport media (*47*). Corrected for unbound drug concentration, Caco-2 permeability was exceptionally high, with no evidence of active efflux. In summary, compound (*S,S*)-**2** as a drug substance is characterized by high permeability, variable solubility depending on the matrix, as well as very high protein binding and low clearance by microsomes and hepatocytes across preclinical and clinical species. Overall, the in vitro metabolic profile, aqueous solubility and blood-plasma partitioning of (*S,S*)-**2** were judged favorable for a potential TCP-1 antimalarial agent with a long in vivo half-life.

**Table 3.** Selected in vitro ADME properties determined for artefenomel (**1**), (±)-**2**, and (*S,S*)-**2**. Additional ADME assay data and values for all controls is provided in Table S1 of the supporting information file.

| ADME property<sup>a</sup> | **1** | (±)-**2** | (S,S)-**2** |
| --- | --- | --- | --- |
| HLM CLint, app (μL/min/mg protein) | 63.7<sup>b</sup> | 6.0<sup>b</sup> | <11.6 |
| MLM CLint, app (μL/min/mg protein) | 8.4<sup>b</sup> | 14.1<sup>b</sup> | n.d. |
| Human plasma stability $t_{1/2}$ (min) | n.d. | n.d. | >480 |
| Mouse plasma stability $t_{1/2}$ (min) | n.d. | n.d. | 251.4 |
| pH 7.4 PBS solubility (μM) | 0.181<sup>c</sup> | 11.5 | 79.7 |
| LogD (octanol/PBS pH 7.4) measured | 5.81<sup>b</sup> | 4.54<sup>b</sup> | n.d. |
| FaSSIF pH 6.5 (μg/mL) | 40.6<sup>c</sup> | n.d. | 2.3 |
| FeSSIF pH 5.0 (μg/mL) | 536<sup>c</sup> | n.d. | 4525 |

<sup>a</sup>human liver microsome (HLM) and mouse liver microsome intrinsic clearance, kinetic solubility in pH 7.4 PBS; thermodynamic solubility in fasted-state simulated intestinal fluid (FaSSIF) or fed-state simulated intestinal fluid (FeSSIF). <sup>b</sup>Values generated in head-to-head experiments contracted by MMV (project code CR302FB2). <sup>c</sup>Data from ref 46. Unless otherwise indicated, ADME data was determined by Quintara Biosciences (South San Francisco, CA). n.d. not determined

We next evaluated the forms (±)-**2** and (*S,S*)-**2** in additional mouse and rat PK studies with both IV and PO administration. A PK study in mice administered (±)-**2** by the oral (50 mg/kg) and IV (10 mg/kg) routes revealed a very large volume of distribution (Vss = 22.7 L/kg) and surprisingly high plasma clearance (CL = 147 mL/min/kg) compared to the in vitro microsome and hepatocyte clearance values (Fig. S11 and Table S3). However, despite high clearance, exposure by the oral route was favorable with a t_1/2_ of 7 hrs and a mean residence time of 9.8 hrs, suggesting that half-life is driven by Vss rather than CL (Table S3). We next performed PK studies of (±)-**2** and (*S,S*)- **2** in Sprague-Dawley rats at different dosing levels, and with collection of urine in the study of (*S,S*)- **2** to evaluate renal clearance. When (±)-**2** was administered to male rats at the same high 50 mg/kg PO and 10 mg/kg IV doses used in the mouse study, we again observed a high volume of distribution (Vss = 34.4 L/kg) and high clearance, but again a prolonged exposure profile with t_1/2_ and MRT of ∼10 hrs (Fig. 4C and Table S4).

Next, we studied (S*,S*)-**2** in SD rats at lower doses of 10 mg/kg PO and 3 mg/kg IV to better compare with published data for **1**, which is reported as blood clearance and blood volume of distribution. Converting plasma CL and Vss values using the measured blood/plasma ratio for (S*,S*)-**2** (1.61) yielded CL_blood_ = 55.8 mL/min/kg and Vss_blood_ = 10.9 L/kg for (*S,S*)-**2**, values that lie between those reported for arterolane and **1** (Table 4 and Table S5). Renal clearance of (*S,S*)-**2** was minimal, at 0.82 mL/min/kg with the fraction in urine <1%, thereby ruling out renal extraction as an important route of clearance (Table S5). Despite relatively high blood clearance, the oral bioavailability of (*S,S*)-**2** was 76%, exactly the same value reported for **1** (Table 4), although it bears noting that different formulations were employed in our PK studies versus in the prior studies of **1**. With this caveat noted, comparing the plasma exposure curves for (*S,S*)-**2** with the historical data for **1** indicates significantly higher exposure of **1** at the earlier timepoints, but similar exposure for **1** and (*S,S*)-**2** at 24 hours (Fig. 4D). The more favorable blood:plasma partitioning of (*S,S*)-**2** compared to **1**, combined with a high volume of distribution and very high protein-binding likely contribute to an extended plasma exposure profile despite blood clearance that approaches the rate of liver blood flow. In any case, the PK profile of (*S,S*)-**2** in rodents, and predicted low human metabolism based on in vitro measures, met one of our key objectives for a next-generation endoperoxide – a significantly extended in vivo half-life as compared to artemisinin.

**Table 4.** Comparison of selected pharmacokinetic parameters for arterolane and artefenomel compared to (*S,S*)-**2** in male SD rats (n = 3 per group).

|  | Log D<sub>7.4</sub> | B:P ratio | IV values (3 mg/kg) |  | PO values (10 mg/kg) |  |
| --- | --- | --- | --- | --- | --- | --- |
|  |  |  | CL<sub>blood</sub><br>(mL/min/kg) | Vss<sub>blood</sub><br>(L/kg) | T<sub>1/2</sub> | F (%) |
| arterolane (OZ277) | 2.6<sup>a</sup> | 1.5<sup>a</sup> | 61 | 4.0 | 0.92 | 13 |
| artefenomel (**1**) | 5.81 | 0.8<sup>a</sup> | 33<sup>†</sup> | 15<sup>†</sup> | 24 | 76 |
| (S,S)-**2** | 4.54 | 1.6 | 55.8 | 10.9 | 9 | 76 |

CL<sub>blood</sub> and Vss<sub>blood</sub> values are calculated from the plasma PK values and blood:plasma (B:P) ratios. Data for arterolane and artefenomel as reported (11). Arterolane measured LogD value as reported (46). <sup>†</sup>IV dose for artefenomel was 2 mg/kg. For reference, liver blood flow (rat) = 68 mL/min/kg.

An second important objective, and one that also influenced the development of **1** (*37*), was improved ferrous iron stability compared to artemisinin and the first-generation trioxolane arterolane, which had shown sub-optimal exposure and serum half-life in infected patients (*48, 49*). In the current landscape of ART-R, enhanced iron stability (along with half-life extension) has been advanced as a criterion for addressing ART-R in next generation endoperoxides (*4, 30*). We therefore compared the cell-free iron reactivity of **1** and (±)-**2** with that of the alkyl carbamate analog (*R,R*)-**6** which is structurally more similar to arterolane. As predicted based on our conformational analyses, aryl carbamates **1** and (±)-**2** were dramatically more stable toward ferrous iron than **6**, while (±)-**2** showed the slowest rate of reaction overall, with >50% of (±)-**2** still intact 24 hours after exposure to excess ferrous ammonium sulfate in an anoxic chamber to prevent iron oxidation by atmospheric oxygen (Table 5 and Fig. S12).

**Table 5.** Relative reactivity of artefenomel (**1**), (±)-**2**, and (*S,S*)-**6** in 50 mM ferrous ammonium sulfate in citrate buffer under anaerobic conditions.

| compound | % parent compound remaining at indicated timepoint |  |  |  |
| --- | --- | --- | --- | --- |
|  | 0 hr | 2 hr | 8 hr | 24 hr |
| **1** | 100% | 83% | 54% | 0% |
| (±)-**2** | 100% | 83% | 79% | 67% |
| (S,S)-**6** | 100% | 0%<sup>a</sup> | 0% | 0% |

<sup>a</sup>(S,S)-**6** was fully consumed by 15 min.

Next, we performed in vitro RSA experiments using both W2 (K13 wild type) and a K13 C580Y mutant strain of *P. falciparum.* Briefly, tightly synchronized 0-3 hr post-invasion ring stage parasites were exposed to 700 nM compound for 6 hours, followed by removal of drug pressure, and quantification of parasitemia after 72 hours. The standard protocol was modified to include additional washing steps and transfer to fresh plates, given the reported difficulty of removing highly protein-bound drug substances from plastic assay plates (*30*). In the W2 parasites, RSA survival rates were similar for DHA (∼10%) and **1** (∼7%), and somewhat lower for (±)-**2**. By contrast, RSA survival rates in the K13 C580Y strain were much higher for DHA (∼50%) than **1** (∼7%) or compound (±)-**2**, which exhibited an RSA survival rate of 0% (Fig. S13). While encouraging, we note again that the RSA assay as typically implemented is designed to reveal parasite susceptibility to artemisinin agents specifically, and not the activity of novel agents in ART-R parasites (*30*). Ideally, future RSA assays of novel agents such as (±)-**2** would employ drug concentrations based on predicted or measured C_max_ values in human subjects.

With the above caveat noted, susceptibility testing (IC_50_) and ex vivo RSAs were next performed for DHA, **1**, and (±)-**2** on clinical isolates from subjects with symptomatic *P. falciparum* infection, including samples collected in 2019 in eastern Uganda and in 2023-24 in northern and eastern Uganda. In the isolates tested in 2019, the median IC_50_ values were 1.5 nM for DHA, 0.5 nM for **1**, and 2.6 nM for compound (±)-**2**, while RSA median survival was 0% for all three test compounds, consistent with a near absence of ART-R in this region in 2019 (Fig. 5A-B). In the larger collection of isolates from 2023-24, median IC_50_ values were 3.8 nM for DHA, 3.0 nM for **1**, and 3.6 nM for (±)-**2**, while RSA median survival was 5.3% (range 0.0%-39.1%) for DHA, 0.0% for **1**, and 0.0% for (±)-**2** (Fig. 5C-D and Table 6). The higher median RSA values for DHA in 2023-24 as compared to 2019 are consistent with emergent ART-R over this timeframe (*20, 21*). Overall, the improved physiochemical properties, predicted human metabolism, enhanced iron stability, and superior ex vivo RSAs of compound **2** revealed the *trans*-3” aryl chemotype to be a promising new sub-type of endoperoxide agents with single-exposure efficacy in mice and activity against clinically relevant ART-R parasites.

**Fig. 5.**
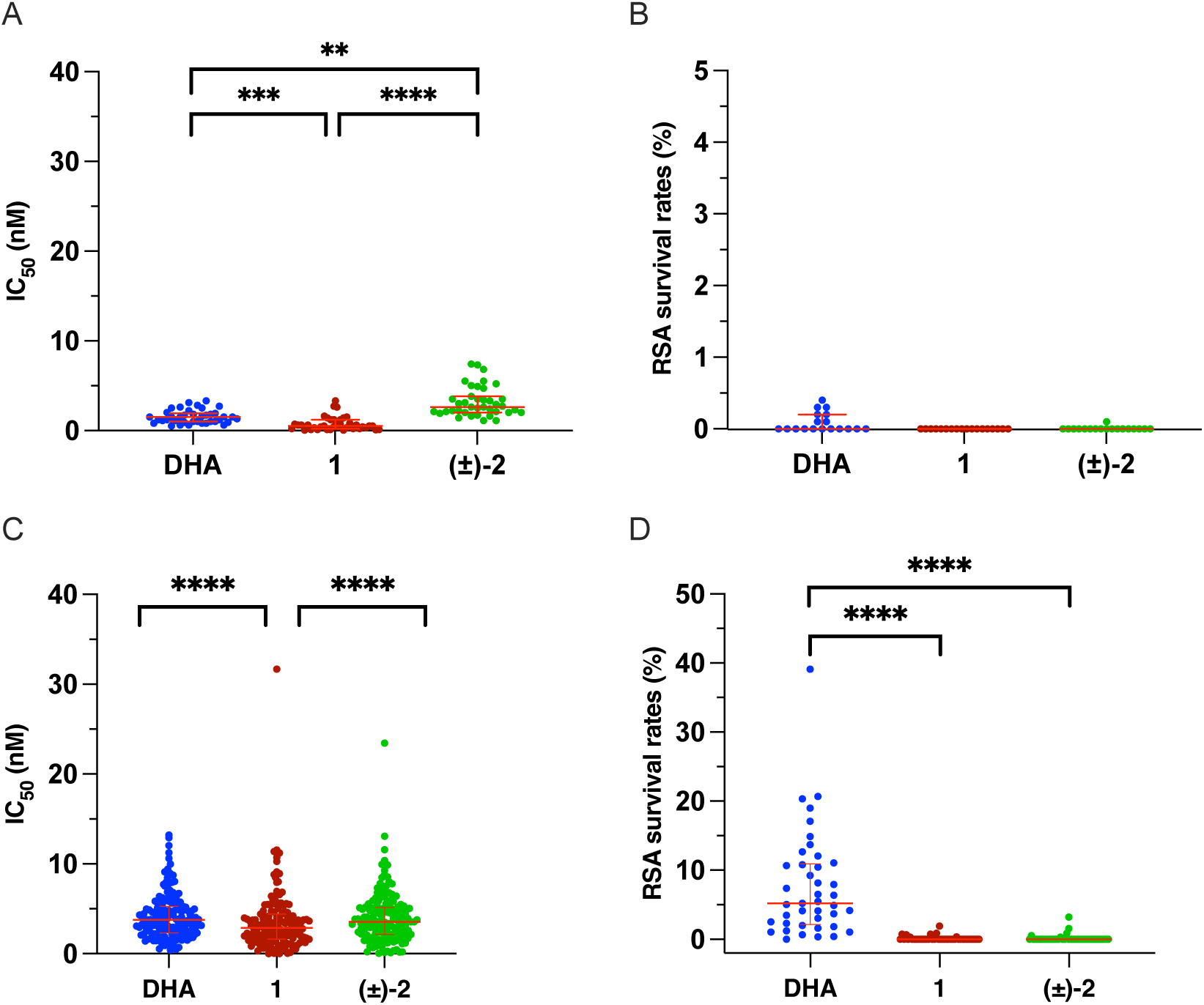
Drug susceptibility of *P. falciparum* field isolates from Uganda determined in 2019 (A and B) and 2023-24 (C and D) against dihydroartemisinin (DHA), artefenomel (**1**), and (±)-**2** measured as standard growth inhibition assay IC_50_ values (A and C) and ring-stage survival assay (RSA) survival rates (B and D; note different scale on Y-axie). Multiple comparisons were made using the Friedman test for paired measures with Dunn’s post-hoc multiple comparison test (*, P < 0.05; **, P < 0.01; ***, P < 0.001; ****, P < 0.0001); ns, not significant.

**Table 6.** Drug susceptibility of Ugandan *P. falciparum* isolates tested in 2023-24 to **DHA**, **1**, and (±)-**2** by region.

|  | Northern Uganda (Kalongo) |  |  | Eastern Uganda (Tororo) |  |  |
| --- | --- | --- | --- | --- | --- | --- |
| Compound | DHA | **1** | **(±)-2** | DHA | **1** | **(±)-2** |
| IC<sub>50</sub> assays performed, n | 115 | 115 | 115 | 51 | 51 | 51 |
| Successful IC<sub>50</sub> assays, n | 106 | 106 | 106 | 47 | 47 | 45 |
| IC<sub>50</sub> values (nM) | **3.7** | **2.3** | **3.5** | **4.1** | **3.8** | **3.9** |
| median (IQR) | (2.2-5.1) | (1.2-3.9) | (1.7-5.0) | (2.8-6.8) | (2.5-6.0) | (2.8-6.1) |
| RSAs with two reads, n | 32 | 32 | 32 | 28 | 28 | 28 |
| RSAs with valid reads (n) | 22 | 22 | 22 | 20 | 20 | 20 |
| RSA survival rate (%)<sup>a</sup> | **3.6** | **0.0** | **0.0** | **7.8** | **0.0** | **0.0** |
| median (IQR) | (1.2-8.8) | (0.0-0.0) | (0.0-0.0) | (4.1-12.5) | (0.0-0.0) | (0.0-0.0) |

## Discussion

The current study was motivated by the worrisome emergence of ART-R in malaria endemic regions and by the recent decision to halt further clinical evaluation of **1**. The remarkable efficacy of **1** in mouse malaria models and a keen desire for agents that achieve single-exposure cure led to its nomination as a development candidate. However, achieving single-dose efficacy clinically proved challenging, especially considering the high doses of **1** and partner drug required, and the significant formulation problems encountered with **1**. We hypothesized that the transverse molecular symmetry and amphiphilic nature of **1** likely contribute to its poor solubility and the formation (*45*) of micelles and lamellar phases in aqueous solution. By breaking symmetry with *trans*-3” substitution as in **2**, we sought to both improve aqueous solubility while maintaining pharmacologically desirable rates of iron reactivity. Although we did observe significant improvements in aqueous solubility (in kinetic assays), further studies will be required to fully assess the effects of de-symmetrization on more complex solution-phase behavior.

The unusual iron-reactive and pleiotropic pharmacology of endoperoxide-class agents arguably requires design criteria distinct from agents that act by a more typical, reversible, drug-like mechanism. Thus, the “free drug hypothesis” posits that only unbound, free drug can exert a pharmacodynamic effect at its target, making the free drug fraction and unbound clearance important parameters for predicting efficacy and dose. The particular case of endoperoxide antimalarials is likely more akin to that of covalent drugs, in that covalent cross-linking reactions to parasite protein are responsible for the pharmacodynamic effect. The interplay of endoperoxide binding affinity for parasite vs. host proteins, and the kinetics of iron activation and cross linking is undoubtedly complex, and how these factors impact efficacy is largely unknown. We hypothesize however that to address ART-R, a high level of binding to *parasite* protein is almost certainly a favorable property, insofar as it encourages retention of stable, active endoperoxide species through the transition from resistant rings to more susceptible trophozoite and schizont parasite stages, where activating free iron is again abundant.

With these considerations in mind and guided by initial assessment for in vitro potency, HLM stability, and aqueous solubility, we identified the favorable properties of *trans*-3” trioxolane analogs bearing para-aryl carbamate side chains. In addition to **2**, other promising exemplars of this chemotype include **18** and **21** (Fig. S10), both of which combine single-digit nM in vitro potencies with robust HLM stability and good to excellent aqueous solubility. The most extensive in vitro ADME and in vivo PK/PD profiling was performed on the enantiopure form (*S,S*)-**2,** which exhibited excellent stability toward proteolytic and drug-metabolizing enzymes from major preclinical and clinical species. In a murine *P. berghei* model, (*S,S*)-**2** afforded cures of all animals following a single 50 mg/kg oral dose. In both the mouse and rat, **2** exhibited a very high volume of distribution that likely contributed to an extended in vivo half-life, despite higher than predicted hepatic clearance. Good plasma stability and low renal clearance ruled out these alternate pathways of in vivo clearance. The very high levels of plasma protein binding observed for **2** confounded the measurement of free drug concentrations and clearance, but despite this, the notably higher blood:plasma ratio for (*S,S*)-**2** compared to **1** suggests a favorable partitioning into erythrocytes, the site of antiplasmodial action.

In studies with cultured *P. falciparum*, (*S,S*)-**2** demonstrated potent activity, with IC_50_ values similar to those of DHA and **1** against artemisinin-sensitive parasites and, as for **1**, improved RSA activity compared to DHA against ART-R laboratory and clinical strains of *P. falciparum*. These results offer promise for (*S,S*)-**2** or related, further optimized analogs to treat drug sensitive and ART-R falciparum malaria. Ideally, a next-generation trioxolane will produce cures in patients infected by artemisinin-sensitive or ART-R falciparum malaria, with reduced dosing frequencies, and will be readily co-formulated with suitable partner drug. Considering this, the artefenomel-adjacent chemotype described herein, as exemplified by (*S,S*)-**2**, offers an important advance on the road to next-generation antimalarial combinations that provides reliable treatment of malaria in the setting of ART-R.

## Materials and Methods

Additional *Methods* are found in *SI Appendix*.

### Materials

All chemical reagents were obtained commercially and used without further purification unless otherwise stated. Anhydrous solvents were purchased from Sigma-Aldrich and were used without further purification. Solvents used for flash column chromatography and workup procedures were purchased form either Sigma-Aldrich or Fisher Scientific. Column chromatography was performed on Silicycle Sili-prep cartridges using a Biotage Isolera Four automated flash chromatography system. Compounds (±)-**3** (*41*) and (*R,R*)-**6** (*40*) were prepared as described previously by our group.

### Ferrous Iron Stability Studies of 1, (±)-2, and (*R,R*)-6

A 1 mM solution of test compound (prepared from 10 mM DMSO stock solution) was added to citrate buffer containing 50 mM Fe(NH_4_)_2_(SO_4_)_2_•6H_2_O in an Ar(g)-purged glove bag to minimize oxidation of the ferrous iron salt. The solution was transferred to a heating block and warmed to 37°C. Aliquots (40 µL) of the reaction mixture were taken at four timepoints (T = 15 min, 2 h, 8 h, 19 h) and a 5 µL portion analyzed by UPLC with UV detection at 190 nm to detect the ketone function shared by reaction products from all three test compounds. Quantification of reaction products was performed by integration of peaks for ketone intermediate in the UV chromatogram. Chromatograms were stacked and plotted in GraphPad Prism software.

### *P. berghei* Mouse Malaria Model

Female Swiss Webster mice (∼20 g body weight) were infected intraperitoneally with 10^6^ *P. berghei*-infected erythrocytes collected from a previously infected mouse. Beginning 1 hr after inoculation the mice were treated once daily by oral gavage for 1 or 2 days as indicated with 100 μL of solution of test compound formulated in 10% DMSO, 40% of a 20% 2-hydroxypropyl-β-cyclodextrin solution in water, and 50% PEG400. There were five mice in each test arm. Infections were monitored by daily microscopic evaluation of Giemsa-stained blood smears starting on day seven. Parasitemia was determined by counting the number of infected erythrocytes per 1000 erythrocytes. Body weight was measured over the course of the treatment. Mice were euthanized when parasitemia exceeded 50% or when weight loss of more than 15% occurred. Parasitemia, animal survival, and morbidity were closely monitored for 30 days post-infection, when experiments were terminated.

### Pharmacokinetic Studies in Mice

The mouse pharmacokinetic study of **1**, (±)-**2,** and (*R,R*)-**6** with PO dosing at 50 mg/kg (**Fig. 2F**) were performed in 8 week-old male CD-1 mice (25-30 g, n = 6 per group) using a formulation of 10% DMSO, 40% of a 20% 2-hydroxypropyl-β-cyclodextrin solution in water, and 50% PEG400. Sampling was at 0.5, 2, 6, and 24 hrs. Two blood samples (300 μL each) were collected from each mouse; three mice were assigned for each time point. The second/final blood collection was terminal. The plasma drug level data will be analyzed using Phoenix WinNonlin (version 6.3) software to perform noncompartmental modeling.

The mouse pharmacokinetic study of (±)-**2** with IV (10 mg/kg) and PO (50 mg/kg) dosing (**Fig. 3C**) was performed in male ICR mice (18-22 g, n = 3 per group) with a formulation of 2% DMSO: 98% of a 10.5% 2-hydroxypropyl-β-cyclodextrin solution in water. Microsampling (40 μL) via facial vein was performed at 0, 0.083, 0.25, 0.5, 1, 2, 4, 8, and 24 hrs into K_2_EDTA tubes. The blood samples were collected and centrifuged to obtain plasma (8000 rpm, 5 min) within 15 minutes post sampling. Nine blood samples were collected from each mouse; three samples were collected for each time point. Data was processed by Phoenix WinNonlin (version 8.3); samples below limit of quantitation were excluded in the PK parameters and mean concentration calculation.

### Pharmacokinetic Studies in Rat

The rat pharmacokinetic study of (±)-**2** with IV (10 mg/kg) and PO (50 mg/kg) dosing (**Fig. 3D**) was performed male SD rat (180-250 g, n = 3 per group) using a formulation of 2% DMSO: 98% of a 10.5% 2-hydroxypropyl-β-cyclodextrin solution in water. Blood samples (100 μL) were collected via facial vein at 0, 0.083, 0.25, 0.5, 1, 2, 4, 8, and 24 hrs into K_2_EDTA tubes. The blood samples were collected and centrifuged to obtain plasma (8000 rpm, 5 min) within 15 minutes post sampling. Nine blood samples were collected from each mouse; three samples were collected for each time point. Data was processed by Phoenix WinNonlin (version 8.3); samples below limit of quantitation were excluded in the PK parameters and mean concentration calculation.

The rat pharmacokinetic study of (*S,S*)-**2** with IV (3 mg/kg) and PO (10 mg/kg) dosing (**Fig. 3D**) was performed male SD rat (220-250 g, n = 3 per group) using a formulation of 10% DMSO, 40% of a 20% 2-hydroxypropyl-β-cyclodextrin solution in water, and 50% PEG400. Blood samples (150 μL) were collected via jugular vein at 0, 0.083, 0.25, 0.5, 1, 2, 4, 8, and 24 hrs into K_2_EDTA tubes. The blood samples were collected and centrifuged to obtain plasma (2000 g, 5 min) within 15 minutes post sampling. Nine blood samples were collected from each mouse; three samples were collected for each time point. Urine was collected at 0-4 hrs, 4-8 hrs, and 8-24 hrs. Data was processed by Phoenix WinNonlin (version 8.2); samples below limit of quantitation were excluded in the PK parameters and mean concentration calculation.

## List of Supplementary Materials

Dataset S1: X-ray Crystal Structure Report for (*R,R*)-**2** (RLA-5763)

Supporting Materials Text

Synthetic Methods

Additional Experimental Methods

Fig. S1. Canonical mechanism of 1,2,4-trioxolane activation

Fig. S2. Conformational dynamics underlying 1,2,4-trioxolane activation and antiplasmodial action

Fig. S3. Synthesis of racemic (±)-**2** (RLA-4735)

Fig. S4. Synthesis of (*R,R*)-**2** (RLA-5763) Fig. S5. Synthesis of (*S,S*)-**2** (RLA-5764)

Fig, S6. Synthesis of (±)-**3** (RLA-3107) Fig. S7. Synthesis of (±)-**4** (RLA-4741)

Fig. S8. Synthesis of (±)-**5** (RLA-5351) Fig. S9. General synthetic schemes

Fig. S10. Summary of Structure-Activity Relationships (SAR) for ortho, meta, and para- substituted analogs with aryl carbamate substitution

Fig. S11. Pharmacokinetic profile of (±)-**2** following IV or PO administration in ICR mice

Fig. S12. Iron reactivity studies with **1**, (±)-**2**, and (*R,R*)-**6**

Fig. S13. In vitro RSA survival rates for laboratory strains.

Table S1. Pharmacokinetic parameters from PK study of **1**, (±)-**2**, and (*R,R*)-**6** male CD-1 mice

Table S2. In vitro ADME Properties of (*S,S*)-**2** and relevant controls

Table S3. Pharmacokinetic parameters for (±)-**2** following IV (10 mg/kg) and PO (50 mg/kg) doses in male ICR mice

Table S4. Pharmacokinetic parameters for (±)-**2** following IV (10 mg/kg) and PO (50 mg/kg) doses in male SD rats

Table S5. Pharmacokinetic parameters for (*S,S*)-**2** following IV (3 mg/kg) and PO (10 mg/kg) doses in male SD rats

Table S6. Sample characteristics and drug susceptibility of Ugandan *P. falciparum* isolates tested in 2023 to DHA, **1**, and (±)-**2**

Supporting Materials References

## Acknowledgments

We thank Drs. Susan Charman and Darren Creek (Monash Institutes of Pharmaceutical Sciences) for helpful discussions and Dr. Charman for reviewing the manuscript. We thank Rebecca Christofferson (Louisiana State University) for help with statistical analyses. Selected in vitro ADME assay data (Table 2) was generated by CROs under contract to Medicines for Malaria Venture (Project Code: CR203FB2). All remaining ADME assay data was performed by Quintara Biosciences (South San Francisco, CA). Pharmacokinetic study in CD-1 mice was performed by SRI Biosciences under contract with NIAID Preclinical Services (DMID contract No. HHSN272201100022I). Pharmacokinetic studies of (±)-**2** in ICR mice and SD rat were performed by Tatara Therapeutics (San Francisco, CA). Pharmacokinetic studies of (*S,S*)-**2** in SD rat were performed by Chempartner Corporation (South San Francisco, CA).

## Author contributions

M.T.K., P.T., B.R.B., J.T., J.N.B., P.J.R., R.A.C., and A.R.R. conceived of study and designed research.

M.T.K., P.T., B.R.B., S.C., J.C., R.L.G., P.J., J.L., V.M., A.N., M.O., S.O., N.S.S., J.T., Y.T.,

P.K.T. and A.V. performed experiments and analyzed data.

S.D.F. and G.J.L. oversaw pharmacokinetic experiments and analyzed data.

M.T.K. and A.R.R. wrote the manuscript.

All authors reviewed and edited the manuscript.

## Data materials availability

All data are available in the main text or supplemental materials. A crystallographic dataset is available as a separate file associated with this manuscript online.

## References and Notes

1. World malaria report 2023 (Geneva: World Health Organization; 2023.Licence: CC BY-NC-SA 3.0 IGO, 2023).

2. S. R. Meshnick, A. Thomas, A. Ranz, C. M. Xu, H. Z. Pan, Artemisinin (qinghaosu): the role of intracellular hemin in its mechanism of antimalarial action., Mol. Biochem. Parasitol. 49, 181–189 (1991).

3. G. H. Posner, C. H. Oh, Regiospecifically oxygen-18 labeled 1,2,4-trioxane: a simple chemical model system to probe the mechanism(s) for the antimalarial activity of artemisinin (qinghaosu)., J. Am. Chem. Soc. 114, 8328–8329 (1992).

4. N. Klonis, D. J. Creek, L. Tilley, Iron and heme metabolism in Plasmodium falciparum and the mechanism of action of artemisinins., Curr. Opin. Microbiol. 16, 722–727 (2013).

5. J. Wang, C.-J. Zhang, W. N. Chia, C. C. Y. Loh, Z. Li, Y. M. Lee, Y. He, L.-X. Yuan, T. K. Lim, M. Liu, C. X. Liew, Y. Q. Lee, J. Zhang, N. Lu, C. T. Lim, Z.-C. Hua, B. Liu, H.-M. Shen, K. S. W. Tan, Q. Lin, Haem-activated promiscuous targeting of artemisinin in Plasmodium falciparum., Nat. Commun. 6, 10111 (2015).

6. H. M. Ismail, V. Barton, M. Phanchana, S. Charoensutthivarakul, M. H. L. Wong, J. Hemingway, G. A. Biagini, P. M. O’Neill, S. A. Ward, Artemisinin activity-based probes identify multiple molecular targets within the asexual stage of the malaria parasites Plasmodium falciparum 3D7., Proc Natl Acad Sci USA 113, 2080–2085 (2016).

7. G. Siddiqui, C. Giannangelo, A. De Paoli, A. K. Schuh, K. C. Heimsch, D. Anderson, T. G. Brown, C. A. MacRaild, J. Wu, X. Wang, Y. Dong, J. L. Vennerstrom, K. Becker, D. J. Creek, Peroxide Antimalarial Drugs Target Redox Homeostasis in Plasmodium falciparum Infected Red Blood Cells., ACS Infect. Dis. 8, 210–226 (2022).

8. N. J. White, Assessment of the pharmacodynamic properties of antimalarial drugs in vivo., Antimicrob. Agents Chemother. 41, 1413–1422 (1997).

9. C. W. Jefford, Synthetic peroxides as potent antimalarials. News and views., Curr. Top. Med. Chem. 12, 373–399 (2012).

10. J. L. Vennerstrom, S. Arbe-Barnes, R. Brun, S. A. Charman, F. C. K. Chiu, J. Chollet, Y. Dong, A. Dorn, D. Hunziker, H. Matile, K. McIntosh, M. Padmanilayam, J. Santo Tomas, C. Scheurer, B. Scorneaux, Y. Tang, H. Urwyler, S. Wittlin, W. N. Charman, Identification of an antimalarial synthetic trioxolane drug development candidate., Nature 430, 900–904 (2004).

11. S. A. Charman, S. Arbe-Barnes, I. C. Bathurst, R. Brun, M. Campbell, W. N. Charman, F. C. K. Chiu, J. Chollet, J. C. Craft, D. J. Creek, Y. Dong, H. Matile, M. Maurer, J. Morizzi, T. Nguyen, P. Papastogiannidis, C. Scheurer, D. M. Shackleford, K. Sriraghavan, L. Stingelin, Y. Tang, H. Urwyler, X. Wang, K. L. White, S. Wittlin, L. Zhou, J. L. Vennerstrom, Synthetic ozonide drug candidate OZ439 offers new hope for a single-dose cure of uncomplicated malaria., Proc Natl Acad Sci USA 108, 4400–4405 (2011).

12. H. S. Kim, J. T. Hammill, R. K. Guy, Seeking the Elusive Long-Acting Ozonide: Discovery of Artefenomel (OZ439)., J. Med. Chem. 60, 2651–2653 (2017).

13. 13. malERA Consultative Group on Drugs, A research agenda for malaria eradication: drugs., PLoS Med. 8, e1000402 (2011).

14. J. N. Burrows, R. H. van Huijsduijnen, J. J. Möhrle, C. Oeuvray, T. N. C. Wells, Designing the next generation of medicines for malaria control and eradication., Malar. J. 12, 187 (2013).

15. N. J. White, F. H. Nosten, SERCAP: is the perfect the enemy of the good?, Malar. J. 20, 281 (2021).

16. A. P. Phyo, P. Jittamala, F. H. Nosten, S. Pukrittayakamee, M. Imwong, N. J. White, S. Duparc, F. Macintyre, M. Baker, J. J. Möhrle, Antimalarial activity of artefenomel (OZ439), a novel synthetic antimalarial endoperoxide, in patients with Plasmodium falciparum and Plasmodium vivax malaria: an open-label phase 2 trial., Lancet Infect. Dis. 16, 61–69 (2016).

17. F. Macintyre, Y. Adoke, A. B. Tiono, T. T. Duong, G. Mombo-Ngoma, M. Bouyou-Akotet, H. Tinto, Q. Bassat, S. Issifou, M. Adamy, H. Demarest, S. Duparc, D. Leroy, B. E. Laurijssens, S. Biguenet, A. Kibuuka, A. K. Tshefu, M. Smith, C. Foster, I. Leipoldt, P. G. Kremsner, B. Q. Phuc, A. Ouedraogo, M. Ramharter, OZ-Piperaquine Study Group, A randomised, double-blind clinical phase II trial of the efficacy, safety, tolerability and pharmacokinetics of a single dose combination treatment with artefenomel and piperaquine in adults and children with uncomplicated Plasmodium falciparum malaria., BMC Med. 15, 181 (2017).

18. Y. Adoke, R. Zoleko-Manego, S. Ouoba, A. B. Tiono, G. Kaguthi, J. E. Bonzela, T. T. Duong, A. Nahum, M. Bouyou-Akotet, B. Ogutu, A. Ouedraogo, F. Macintyre, A. Jessel, B. Laurijssens, M. H. Cherkaoui-Rbati, C. Cantalloube, A. C. Marrast, R. Bejuit, D. White, T. N. C. Wells, F. Wartha, D. Leroy, A. Kibuuka, G. Mombo-Ngoma, D. Ouattara, I. Mugenya, B. Q. Phuc, F. Bohissou, D. P. Mawili-Mboumba, F. Olewe, I. Soulama, H. Tinto, FALCI Study Group, A randomized, double-blind, phase 2b study to investigate the efficacy, safety, tolerability and pharmacokinetics of a single-dose regimen of ferroquine with artefenomel in adults and children with uncomplicated Plasmodium falciparum malaria., Malar. J. 20, 222 (2021).

19. Y. He, S. Campino, E. Diez Benavente, D. C. Warhurst, K. B. Beshir, I. Lubis, A. R. Gomes, J. Feng, W. Jiazhi, X. Sun, F. Huang, L.-H. Tang, C. J. Sutherland, T. G. Clark, Artemisinin resistance-associated markers in Plasmodium falciparum parasites from the China-Myanmar border: predicted structural stability of K13 propeller variants detected in a low-prevalence area., PLoS ONE 14, e0213686 (2019).

20. M. D. Conrad, V. Asua, S. Garg, D. Giesbrecht, K. Niaré, S. Smith, J. F. Namuganga, T. Katairo, J. Legac, R. M. Crudale, P. K. Tumwebaze, S. L. Nsobya, R. A. Cooper, M. R. Kamya, G. Dorsey, J. A. Bailey, P. J. Rosenthal, Evolution of partial resistance to artemisinins in malaria parasites in uganda., N. Engl. J. Med. 389, 722–732 (2023).

21. P. J. Rosenthal, V. Asua, M. D. Conrad, Emergence, transmission dynamics and mechanisms of artemisinin partial resistance in malaria parasites in Africa., Nat. Rev. Microbiol. 22, 373–384 (2024).

22. M. D. Conrad, P. J. Rosenthal, Antimalarial drug resistance in Africa: the calm before the storm?, Lancet Infect. Dis. 19, e338–e351 (2019).

23. G. Siddiqui, A. Srivastava, A. S. Russell, D. J. Creek, Multi-omics Based Identification of Specific Biochemical Changes Associated With PfKelch13-Mutant Artemisinin-Resistant Plasmodium falciparum., J. Infect. Dis. 215, 1435–1444 (2017).

24. J. Birnbaum, S. Scharf, S. Schmidt, E. Jonscher, W. A. M. Hoeijmakers, S. Flemming, C. G. Toenhake, M. Schmitt, R. Sabitzki, B. Bergmann, U. Fröhlke, P. Mesén-Ramírez, A. Blancke Soares, H. Herrmann, R. Bártfai, T. Spielmann, A Kelch13-defined endocytosis pathway mediates artemisinin resistance in malaria parasites., Science 367, 51–59 (2020).

25. H. M. Behrens, S. Schmidt, T. Spielmann, The newly discovered role of endocytosis in artemisinin resistance., Med. Res. Rev. 41, 2998–3022 (2021).

26. N. F. Gnädig, B. H. Stokes, R. L. Edwards, G. F. Kalantarov, K. C. Heimsch, M. Kuderjavy, A. Crane, M. C. S. Lee, J. Straimer, K. Becker, I. N. Trakht, A. R. Odom John, S. Mok, D. A. Fidock, Insights into the intracellular localization, protein associations and artemisinin resistance properties of Plasmodium falciparum K13., PLoS Pathog. 16, e1008482 (2020).

27. N. Klonis, M. P. Crespo-Ortiz, I. Bottova, N. Abu-Bakar, S. Kenny, P. J. Rosenthal, L. Tilley, Artemisinin activity against Plasmodium falciparum requires hemoglobin uptake and digestion., Proc Natl Acad Sci USA 108, 11405–11410 (2011).

28. S. C. Xie, C. Dogovski, E. Hanssen, F. Chiu, T. Yang, M. P. Crespo, C. Stafford, S. Batinovic, S. Teguh, S. Charman, N. Klonis, L. Tilley, Haemoglobin degradation underpins the sensitivity of early ring stage Plasmodium falciparum to artemisinins., J. Cell Sci. 129, 406–416 (2016).

29. B. Witkowski, D. Menard, C. Amaratunga, R. M. Fairhurst, Ring-stage Survival Assays (RSA) to evaluate the in-vitro and ex-vivo susceptibility of Plasmodium falciparum to artemisinins., Natl. Inst. Health Proced. RSAv1, 1–16 (2013).

30. T. Yang, S. C. Xie, P. Cao, C. Giannangelo, J. McCaw, D. J. Creek, S. A. Charman, N. Klonis, L. Tilley, Comparison of the Exposure Time Dependence of the Activities of Synthetic Ozonide Antimalarials and Dihydroartemisinin against K13 Wild-Type and Mutant Plasmodium falciparum Strains., Antimicrob. Agents Chemother. 60, 4501–4510 (2016).

31. J. Straimer, N. F. Gnädig, B. H. Stokes, M. Ehrenberger, A. A. Crane, D. A. Fidock, Plasmodium falciparum K13 Mutations Differentially Impact Ozonide Susceptibility and Parasite Fitness In Vitro., MBio 8 (2017), doi:10.1128/mBio.00172-17.

32. C. Giannangelo, L. Stingelin, T. Yang, L. Tilley, S. A. Charman, D. J. Creek, Parasite- Mediated Degradation of Synthetic Ozonide Antimalarials Impacts In Vitro Antimalarial Activity., Antimicrob. Agents Chemother. 62 (2018), doi:10.1128/AAC.01566-17.

33. P. M. O’Neill, G. H. Posner, A medicinal chemistry perspective on artemisinin and related endoperoxides., J. Med. Chem. 47, 2945–2964 (2004).

34. Y. Dong, J. Chollet, H. Matile, S. A. Charman, F. C. K. Chiu, W. N. Charman, B. Scorneaux, H. Urwyler, J. Santo Tomas, C. Scheurer, C. Snyder, A. Dorn, X. Wang, J. M. Karle, Y. Tang, S. Wittlin, R. Brun, J. L. Vennerstrom, Spiro and dispiro-1,2,4-trioxolanes as antimalarial peroxides: charting a workable structure-activity relationship using simple prototypes., J. Med. Chem. 48, 4953–4961 (2005).

35. H. M. Ismail, V. E. Barton, M. Panchana, S. Charoensutthivarakul, G. A. Biagini, S. A. Ward, P. M. O’Neill, A Click Chemistry-Based Proteomic Approach Reveals that 1,2,4-Trioxolane and Artemisinin Antimalarials Share a Common Protein Alkylation Profile., Angew. Chem. Int. Ed 55, 6401–6405 (2016).

36. D. J. Creek, W. N. Charman, F. C. K. Chiu, R. J. Prankerd, K. J. McCullough, Y. Dong, J. L. Vennerstrom, S. A. Charman, Iron-mediated degradation kinetics of substituted dispiro-1,2,4- trioxolane antimalarials., J. Pharm. Sci. 96, 2945–2956 (2007).

37. Y. Dong, X. Wang, S. Kamaraj, V. J. Bulbule, F. C. K. Chiu, J. Chollet, M. Dhanasekaran, C. D. Hein, P. Papastogiannidis, J. Morizzi, D. M. Shackleford, H. Barker, E. Ryan, C. Scheurer, Y. Tang, Q. Zhao, L. Zhou, K. L. White, H. Urwyler, W. N. Charman, H. Matile, S. Wittlin, S. A. Charman, J. L. Vennerstrom, Structure-Activity Relationship of the Antimalarial Ozonide Artefenomel (OZ439)., J. Med. Chem. 60, 2654–2668 (2017).

38. S. D. Fontaine, A. G. DiPasquale, A. R. Renslo, Efficient and stereocontrolled synthesis of 1,2,4-trioxolanes useful for ferrous iron-dependent drug delivery., Org. Lett. 16, 5776–5779 (2014).

39. B. R. Blank, J. Gut, P. J. Rosenthal, A. R. Renslo, Enantioselective synthesis and in vivo evaluation of regioisomeric analogues of the antimalarial arterolane., J. Med. Chem. 60, 6400– 6407 (2017).

40. B. R. Blank, R. L. Gonciarz, P. Talukder, J. Gut, J. Legac, P. J. Rosenthal, A. R. Renslo, Antimalarial Trioxolanes with Superior Drug-Like Properties and In Vivo Efficacy., ACS Infect. Dis. 6, 1827–1835 (2020).

41. B. R. Blank, J. Gut, P. J. Rosenthal, A. R. Renslo, Artefenomel Regioisomer RLA-3107 Is a Promising Lead for the Discovery of Next-Generation Endoperoxide Antimalarials., ACS Med. Chem. Lett. 14, 493–498 (2023).

42. J. N. Burrows, S. Duparc, W. E. Gutteridge, R. Hooft van Huijsduijnen, W. Kaszubska, F. Macintyre, S. Mazzuri, J. J. Möhrle, T. N. C. Wells, New developments in anti-malarial target candidate and product profiles., Malar. J. 16, 26 (2017).

43. Y. Tang, Y. Dong, J. M. Karle, C. A. DiTusa, J. L. Vennerstrom, Synthesis of tetrasubstituted ozonides by the Griesbaum coozonolysis reaction: diastereoselectivity and functional group transformations by post-ozonolysis reactions., J. Org. Chem. 69, 6470–6473 (2004).

44. M. Salim, J. Khan, G. Ramirez, A. J. Clulow, A. Hawley, H. Ramachandruni, B. J. Boyd, Interactions of Artefenomel (OZ439) with Milk during Digestion: Insights into Digestion-Driven Solubilization and Polymorphic Transformations., Mol. Pharm. 15, 3535–3544 (2018).

45. A. J. Clulow, M. Salim, A. Hawley, E. P. Gilbert, B. J. Boyd, The Curious Case of the OZ439 Mesylate Salt: An Amphiphilic Antimalarial Drug with Diverse Solution and Solid State Structures., Mol. Pharm. 15, 2027–2035 (2018).

46. S. A. Charman, A. Andreu, H. Barker, S. Blundell, A. Campbell, M. Campbell, G. Chen, F. C. K. Chiu, E. Crighton, K. Katneni, J. Morizzi, R. Patil, T. Pham, E. Ryan, J. Saunders, D. M. Shackleford, K. L. White, L. Almond, M. Dickins, D. A. Smith, J. J. Moehrle, J. N. Burrows, N. Abla, An in vitro toolbox to accelerate anti-malarial drug discovery and development., Malar. J. 19, 1 (2020).

47. K. Katneni, T. Pham, J. Saunders, G. Chen, R. Patil, K. L. White, N. Abla, F. C. K. Chiu, D. M. Shackleford, S. A. Charman, Using Human Plasma as an Assay Medium in Caco-2 Studies Improves Mass Balance for Lipophilic Compounds., Pharm. Res. 35, 210 (2018).

48. N. Valecha, S. Looareesuwan, A. Martensson, S. M. Abdulla, S. Krudsood, N. Tangpukdee, S. Mohanty, S. K. Mishra, P. K. Tyagi, S. K. Sharma, J. Moehrle, A. Gautam, A. Roy, J. K. Paliwal, M. Kothari, N. Saha, A. P. Dash, A. Björkman, Arterolane, a new synthetic trioxolane for treatment of uncomplicated Plasmodium falciparum malaria: a phase II, multicenter, randomized, dose-finding clinical trial., Clin. Infect. Dis. 51, 684–691 (2010).

49. N. Valecha, D. Savargaonkar, B. Srivastava, B. H. K. Rao, S. K. Tripathi, N. Gogtay, S. K. Kochar, N. B. V. Kumar, G. C. Rajadhyaksha, J. D. Lakhani, B. B. Solanki, R. K. Jalali, S. Arora, A. Roy, N. Saha, S. S. Iyer, P. Sharma, A. R. Anvikar, Comparison of the safety and efficacy of fixed-dose combination of arterolane maleate and piperaquine phosphate with chloroquine in acute, uncomplicated Plasmodium vivax malaria: a phase III, multicentric, open-label study., Malar. J. 15, 42 (2016).

