## Supplemental Files for "Identifying a next-generation antimalarial trioxolane in a landscape of artemisinin partial resistance"

**This PDF file includes:**

Supporting text
Figures S1 to S13
Tables S1 to S6
Legends for Datasets S1
SI References

**Other supporting materials for this manuscript include the following:**

Dataset S1: X-ray Crystal Structure Report for (*R,R*)-**2** (RLA-5763)

### Supporting Materials Text

**Description of synthetic approaches to trans-3' aryl trioxolane analogs.** We have previously reported *(1)* an efficient four-step synthesis of the artefenomel regioisomer (±)-**3** in racemic form as a pure *trans* diastereomer. This route proceeded via stereocontrolled Griesbaum co-ozonolysis of the 3"-subsituted cyclohexanone **S1** to afford **S2** as predominantly the desired *trans* diastereomer (Fig**.** S3). After deprotection of the acetate to afford free phenol **S3**, alkylation of the phenolic function with 4-(2-chloroethyl)morpholine hydrochloride afforded the final product (±)-**3** as a single diastereomer within the limits of $^{1}$H NMR analysis (Fig**.** S6). A directly analogous approach was used to prepare the *meta* and *ortho* substituted congeners (±)-**4** and (±)-**5** described in the current communication. Thus, *in situ* generation of a palladium (**II**) catalytic complex obtained by mixing Pd(acac)$_2$, dppben, and Cu(BF$_4$)$_2$·H$_2$O successfully promoted the 1,4-addition of substituted phenyl boronic acids to 2-cyclohexen-1-one to afford the resultant ketones **S9** and **S13** in 73-91% yield (Fig**.** S7 and S8). The phenolic function in meta intermediate **S9** was next protected by reaction with acetic anhydride and pyridine in $CH_2Cl_2$ to yield ketone **S10** in 78% yield (Fig**. S7**). Griesbaum co-ozonolysis of **S10** in the presence of adamantan-2-one and ozone at 0 °C afforded the respective 3'-aryl substituted 1,2,4-trioxolane intermediate **S11** in ~80% yield, in a diastereomeric ratio of ≥ 7:1. Removal of the acetate in **S11** was achieved in 85-93% yield via addition of aqueous potassium hydroxide in THF/MeOH at 50 °C to afford phenol **S12**, which was then alkylated with 4-(2-chloroethyl)morpholine hydrochloride in the presence of powdered NaOH and (Bu)$_4$NHSO$_4$ at 55 °C to yield (±)-**4** (Fig. S7). An analogous process was used for the preparation of ortho isomer (±)-**5**, proceeding through intermediates **S13**-**S16** as illustrated below (Fig**.** S8).

The same phenolic trioxolane intermediates were employed to prepare phenyl carabamate analogs, including the representative carbamates **7**–**23** detailed herein (Fig**.** S9 and S10). Thus, phenolic trioxolanes **S3**, **S12**, and **S16** were activated as a *p*-nitrophenylcarbonate by reaction with 2-(4-nitrophenyl)acetyl chloride, DIPEA, and DMAP in dichloromethane to afford **S4**, **S18**, and **S17**, respectively (Fig**.** S9). To these intermediates was added a primary or secondary amine in triethylamine and dichloromethane to yield the respective para (**17-23**), meta (**9-16**), and ortho (**7-8**) phenolic carbamates in modest to high yield depending on the amine employed (Fig**.** S9). For some diamine analogues, a final Boc deprotection step was required to afford the final, basic amine side chain analogs. This late stage functionalization of the phenolic intermediates with carbamate side chains mirrors our previous studies *(2)* of 3'-*trans* alkylcarbamates prepared from 3'-hydroxycyclohexane trioxolane intermediates, and is amenable to multigram synthesis and a wide scope of amine coupling partners.

To prepare **2** in its enantiopure forms, we employed non-racemic intermediates (*R*)-**S6** and (*S*)-**S6**, prepared as described previously *(3)*. This asymmetric synthesis employs a Rh(acac)(C$_2$H$_4$)$_2$-mediated conjugate addition of 4-(benzyloxy)phenyl boronic acid to cyclohexanone in the presence of enantiopure BINAP ligand with aqueous potassium hydroxide in dioxane at 100 °C to yield the (*R*)-**S6** and (*S*)-**S6** ketones as benzyl ethers (**Fig.** S4 and S5). The benzyl protecting group was then removed via hydrogenolysis over Pd/C in ethyl acetate at 50 °C, followed by acetylation with acetic anhydride and triethylamine in dichloromethane to yield (*R*)-**S2** and (*S*)-**S2**. These ketones were then carried through the same synthetic route as was used previously for (±)-**2** (Fig**. S3**) to yield the final products (*R,R*)-**2** and (*S,S*)-**2** (Fig**.** S4 and S5). The solution of an X-ray crystal structure of (*R,R*)-**2** (Figure **3A**; Dataset **S1** and **S2**) confirmed both the absolute sense of asymmetric induction in the initial Rh–BINAP mediated coupling to afford enantiopure forms of intermediates **S6** and also the *trans* stereochemistry with respect to the peroxo bridge, which arises from the inherent stereoselectivity of the Griesbaum process. Finally, comparison of retention times for (*R,R*)-**2** and (*S,S*)-**2** with material obtained from the separation of (±)-**2** by chiral HPLC, allowed assignment of the two enantiomers obtained by separation of (±)-**2**.

**Structure–Activity Relationships of 3' Aryl Carbamate Analogs.** We evaluated the aryl carbamate analogs for killing of cultured *P. falciparum* parasites, in vitro kinetic solubility in pH 7.4 aqueous PBS, and for their stability in the presence of cultured human liver microsomes (Fig**.** S10). Comparing a congeneric series of regioisomers bearing 4-amino piperidine (**7**, **9**, **2**) or 2-methylpropane-1,2-diamino (**8**, **10**, **17**) carbamates revealed some interesting trends. Hence, all regioisomeric forms within this comparison retained potent, low- or single-digit nM antiparasitic activity that was superior to morpholino ethers **3**-**5**. All six analogs showed aqueous solubility significantly improved over **1**, in the range of 11.5 μM (**2**) to 103 μM (**10**) as compared to a value of 0.181 μM reported for **1**. Human liver microsome stability was excellent ($t_{1/2}$ > 100 min) for meta isomers **9** and **10**, and for para isomers **2** and **17**, while ortho isomers **7** and **8** were measurably less stable ($t_{1/2}$ = 33.8 and 47.4 min, respectively). This relatively more rapid metabolism led us to focus further efforts on the meta and para carbamates with side chains structurally similar to 4-amino piperidine and 2-methylpropane-1,2-diamino. With regard to antiplasmodial activity, the favorable effect of a primary ionizable amine was notable, with many such analogs exhibiting low (**13, 14**, **17**) or single-digit nanomolar (**2**, **9-11**, **16**, **18**, **21**) $IC_{50}$ values against the W2 strain of *P. falciparum*. Less potent were the 3-amino azetidine **20**, both 1,2-diaminoethane carbamates **12** and **19**, and morpholine carbamates **15** and **22**, and thiomorpholine sulfoxide **23**, which exhibited $IC_{50}$ values between ~50-250 nM, substantially less potent than either **1** or the best of the novel analogs, such as **2**, **9**, **11**, and **21**. Metabolic stability as judged with the HLM assay was generally good to excellent across the meta and para scaffolds, with the most stable analogs ($t_{1/2}$ >120 min) being the meta analogs **9**, **10**, and **12**, and the para analogs **2**, **18**, and **21** (Fig. S10). Although compound **2** became the focus of our subsequent PK/PD studies, it is important to note that other para analogs such as **18** and **21** have been so far understudied and represent promising back-up candidates meriting further PK/PD assessment.

### Methods

#### Synthetic Methods

**Materials** All chemical reagents were obtained commercially and used without further purification unless otherwise stated. Anhydrous solvents were purchased from Sigma-Aldrich and were used without further purification. Solvents used for flash column chromatography and workup procedures were purchased form either Sigma-Aldrich or Fisher Scientific. Column chromatography was performed on Silicycle Sili-prep cartridges using a Biotage Isolera Four automated flash chromatography system.

**Instrumentation** NMR spectra were recorded on a Bruker Avance III HD 400 MHz spectrometer (with 5 mm BBFO Z-gradient Smart Probe), calibrated to CH(D)Cl3 as an internal reference (7.26 and 77.00 ppm for $^{1}$H and $^{13}$C NMR spectra, respectively). Data for $^{1}$H NMR spectra are reported in terms of chemical shift (δ, ppm), multiplicity, coupling constant (Hz), and integration. Data for $^{13}$C NMR spectra are reported in terms of chemical shift (δ, ppm), with multiplicity and coupling constants in the case of C-F coupling. The following abbreviations are used to denote these multiplicities: s = singlet, d = doublet, t = triplet, q = quartet, m = multiplet, br = broad, app = apparent, or combinations of these. LC–MS and compound purity were determined using a Waters Micromass ZQ 4000, equipped with a Waters 2795 Separation Module, a Waters 2996 Photodiode Array Detector, and a Waters 2424 ELSD detector. Separations were carried out with an XBridge BEH C18, 3.5 μm, 4.6 mm × 20 mm column, at ambient temperature (unregulated) using a mobile phase of water–methanol containing a constant 0.10% formic acid.

(±)-**S1: 4-(3-oxocyclohexyl)phenyl acetate** To a solution of 4-acetoxyphenylboronic acid (2.79 g, 15.19 mmol) in anhydrous DME (60 mL) was added palladium(II) acetylacetonate (156 mg, 0.51 mmol), 1,2-bis(diphenylphosphino)benzene (231 mg, 0.51 mmol), $Cu(BF_4)_2$ hydrate (484 mg, 2.02 mmol) and 2-cyclohexen-1-one (1 mL, 10.12 mmol). The reaction mixture was stirred at room temperature for 16 h under argon atmosphere. The solvent was removed in *vacuo*. The residue was purified by column chromatography (petroleum ether / ethyl acetate = 1/10) to afford **(±)-S1**

(2.1 g, 91% yield) as a white solid. $^1$H NMR (400 MHz, $CDCl_3$) δ 7.22 (d, *J* = 8.5 Hz, 2H), 7.04 (d, *J* = 8.5 Hz, 2H), 3.08–2.93 (m, 1H), 2.64–2.55 (m, 1H), 2.55–2.42 (m, 2H), 2.42–2.32 (m, 1H), 2.29 (s, 3H), 2.19–2.03 (m, 2H), 1.90–1.70 (m, 2H); $^{13}$C NMR (100 MHz, $CDCl_3$) δ 210.8, 169.6, 149.2, 141.8, 127.5, 121.7, 48.9, 44.1, 41.1, 32.7, 25.4, 21.1. LCMS: Calculated Exact Mass = 232.1, Found $[M+H]^+$ (ESI+) = 233.1

(±)-**S2: *trans*-4-(dispiro[adamantane-2,3'-[1,2,4]trioxolane-5',1''-cyclohexan]-3''-yl)phenyl acetate.**

To a solution of adamantan-2-one *O*-methyl oxime (3.23 g, 18.0 mmol) in $CCl_4$ (100 mL) was added (±)-**S1** (2.1 g, 9.0 mmol) at room temperature. The solution was then cooled to 0 ˚C and bubbled with ozone for 4 h. After the reaction was completed, the mixture was purged with nitrogen for 10 minutes to remove any dissolved ozone. The solvent was removed in *vacuo*. The residue was purified by column chromatography (petroleum ether / ethyl acetate = 20/1) to afford (±)-**S2** (3.1 g, 86% yield) as a white solid. The diastereoselectivity of reaction was determined by $^1$H NMR to be 8.1:1 in favor of the *trans* diastereomer. $^1$H NMR (400 MHz, $CDCl_3$) δ 7.24–7.18 (m, 2H), 7.05–6.98 (m, 2H), 2.95 (tt, *J* = 12.7, 3.1 Hz, 1H, minor diastereomer), 2.81 (tt, *J* = 12.8, 3.3 Hz, 1H), 2.29 (s, 3H), 2.18–2.09 (m, 1H), 2.04–1.55 (m, 20H), 1.44–1.30 (m, 1H); $^{13}$C NMR (100 MHz, $CDCl_3$) δ 169.6, 148.9, 143.3 (minor diastereomer), 143.1, 127.7, 121.4, 111.8 (minor diastereomer), 111.3, 108.9, 46.9 (minor diastereomer), 42.1, 41.6 (minor diastereomer), 41.3, 41.0 (minor diastereomer), 39.2 (minor diastereomer), 36.7, 36.4, 36.3 (minor diastereomer), 35.0 (minor diastereomer), 34.9 (minor diastereomer), 34.8, 34.7, 34.1, 34.0 (minor diastereomer), 33.5 (minor diastereomer), 32.7, 27.4 (minor diastereomer), 26.9 (minor diastereomer), 26.8, 26.4, 23.5, 21.1. LCMS: Calculated Exact Mass = 398.2, Found $[M+NH_4]^+$ (ESI+) = 416.2

(±)-**S3: *trans*-4-(dispiro[adamantane-2,3'-[1,2,4]trioxolane-5',1''-cyclohexan]-3''-yl)phenol.**

To a solution of (±)-**S2** (3.1 g, 7.78 mmol) in THF (35 mL) and MeOH (70 mL) was added a solution of KOH (1.74 g,31.2 mmol) in water (15 mL). The reaction mixture was stirred for 2 h at 50 ˚C. The solvent was removed in *vacuo*. The crude was diluted with water and extracted with EA (50 mL x 3). The combined organic layers were washed with brine, dried over $Na_2SO_4$, filtered and concentrated under reduced pressure. The residue was purified by flash chromatography (PE/EA = 7/1) to afford (±)-**S3** (2.6 g, 93% yield) as an oil that was determined to be a ~15:1 mixture in favor of the *trans* diastereomer. $^1$H NMR (400 MHz, $CDCl_3$) δ 7.12–7.02 (m, 2H), 6.81–6.72 (m, 2H), 4.95 (br s, 1H), 2.89 (tt, *J* = 12.6, 3.3 Hz, 1H, minor diastereomer), 2.74 (tt, *J* = 12.8, 3.3 Hz, 1H), 2.15–2.08 (m, 1H), 2.05–1.64 (m, 19H), 1.64–1.55 (m, 1H), 1.40–1.28 (m, 1H); $^{13}$C NMR (100 MHz, $CDCl_3$) δ 153.9, 138.0, 127.8, 115.2, 111.4, 109.1; 42.3, 41.8 (minor diastereomer), 41.0, 40.7 (minor diastereomer), 36.8, 36.4, 35.0 (minor diastereomer), 34.9 (minor diastereomer), 34.8, 34.7, 34.2, 33.0, 26.8, 26.5, 23.5, 21.1 (minor diastereomer). LCMS: Calculated Exact Mass = 356.2, Found $[M-H]^-$(ESI+) = 355.2

(±)-**S4: *trans*-4-(dispiro[adamantane-2,3'-[1,2,4]trioxolane-5',1''-cyclohexan]-3''-yl)phenyl (4-nitrophenyl) carbonate.**

To an oven-dried round bottom flask containing a magnetic stir bar under an Ar(g) atmosphere was added (±)-**S3** (0.150 mg, 0.42 mmol, 1.0 equiv) dissolved in dichloromethane (10 mL), *N,N*-diisopropylethylamine (0.22 mL, 1.26 mmol, 3.00 equiv), and 4-dimethylaminopyridine (0.062 g, 0.51 mmol, 1.2 equiv). The mixture was cooled to 0 °C and 4-nitrophenyl chloroformate (0.254 g, 1.26 mmol, 3.00 equiv) was added as a solid in one portion. The solution was allowed to stir at room temperature for 3 hours. The reaction was judged complete and the mixture then diluted with $H_2O$ (100 mL) and subsequently extracted with EtOAc (100 mL). The organic layer was washed repeatedly by potassium carbonate solution until the aqueous layer was colorless and no longer yellow (indicating that *p*-nitrophenol had been successfully removed from the organic layer). The organic layer was dried over anhydrous $Na_2SO_4$, filtered, and concentrated under reduced pressure to yield thick yellow oil. The residue was then purified through flash column chromatography (80 g silica gel cartridge, 0–25% EtOAc/Hexanes, product eluted during 10% EtOAc/Hex) to yield the desired intermediate (±)-**S4** (170 mg, 77%) as a colorless solid. $^1$H NMR (400 MHz, $CDCl_3$-d) δ 8.29 - 8.34 (m, 2H), 7.49 (d, *J* = 9.01 Hz, 2H), 7.20 - 7.30 (m, 4H), 2.85 (tt, *J* = 3.01, 12.69 Hz, 1H),

2.16 (br d, $J$ = 13.15 Hz, 1H), 1.91 - 2.05 (m, 7H), 1.65 - 1.88 (m, 13H), 1.37 - 1.45 (m, 1H); $^{13}C$ NMR (100 MHz, $CDCl_3$) δ 155.4, 151.2, 149.0, 145.6, 144.3, 128.1, 125.4, 121.8, 120.7, 111.5, 108.9, 42.1, 41.4, 36.8, 36.5, 34.9, 34.8, 34.2, 32.8, 26.9, 26.5, 23.5. MS (ESI) calculated for $C_{29}H_{31}NNaO_8$ $[M + Na]^+$ *m/z* 544.19, found 544.23.

(±)-**S5: *trans*-4-(dispiro[adamantane-2,3'-[1,2,4]trioxolane-5',1''-cyclohexan]-3''-yl)phenyl 4-((tert-butoxycarbonyl)amino)piperidine-1-carboxylate**

One-pot procedure from (±)-**S3** via (±)-**S4**. To a solution of (±)-**S3** (2.6 g, 7.30 mmol) in dichloromethane (50 mL) was added DIEA (2.83 g, 21.9 mmol) and bis(4-nitrophenyl)carbonate (2.26 g, 7.45 mmol) at room temperature, and the mixture was stirred at room temperature for 2 h until the formation of (±)-**S4** was judged complete by LC/MS analysis. Next, *tert*-butyl piperidin-4-ylcarbamate (2.20 g, 10.95 mmol) was added and the reaction mixture was stirred for 2 h at room temperature. The mixture was poured into $H_2O$ (50 mL) and extracted with dichloromethane (50 mL x 3). The combined organic layers were washed with brine (100 mL), dried over $Na_2SO_4$, filtered, and concentrated under reduced pressure. The residue was purified by column chromatography (PE/EA = 5/1) to afford (±)-**S5** (1.2 g, 29% yield) as a white solid. $^1H$ NMR (400 MHz, $CDCl_3$) δ 7.16 - 7.20 (m, 2H), 6.99 - 7.03 (m, 2H), 4.61 (br d, $J$ = 6.82 Hz, 1H), 4.13 - 4.25 (m, 2H), 3.66 (br s, 1H), 2.93 - 3.16 (m, 2H), 2.75 - 2.83 (m, 1H), 2.12 (br d, $J$ = 13.39 Hz, 1H), 1.89 - 2.05 (m, 9H), 1.59 - 1.86 (m, 15H), 1.44 - 1.51 (m, 9H), 1.33 - 1.44 (m, 3H); $^{13}C$ NMR (100 MHz, $CDCl_3$) δ 155.2, 153.8, 149.7, 142.6, 127.6, 121.6, 111.4, 109.0, 79.6, 47.7, 43.3, 43.2, 42.2, 41.3, 36.8, 36.5, 34.9, 34.8, 34.2, 32.9, 32.6, 28.5, 26.9, 26.5, 23.5. LCMS: Calculated Exact Mass = 582.3, Found $[M+NH_4]^+$ (ESI+) = 600.5

(±)-**2 (RLA-4735): *trans*-4-(dispiro[adamantane-2,3'-[1,2,4]trioxolane-5',1''-cyclohexan]-3''-yl)phenyl 4-aminopiperidine-1-carboxylate.**

To a solution of (±)-**S5** (1.2 g, 2.06 mmol) in MeOH (50 mL) was added dropwise acetyl chloride (3.21 g, 41.2 mmol) at 0 °C, and the reaction mixture was stirred at room temperature for 4 h. After completion, the mixture was concentrated. The residue was poured into water and adjusted to pH = 10 with sat.aq. $NaHCO_3$. The mixture was extracted with $CH_2Cl_2$ (50 mL x 3). The combined organic layers were washed with brine, dried over $Na_2SO_4$, filtered and concentrated under reduced pressure. The residue was purified by column chromatography (MeOH containing 0.7N ammonia | dichloromethane = 1:10) to afford (±)-**2** (650 mg, 65% yield) as a white solid. $^1H$ NMR (400 MHz, $CDCl_3$) δ ppm 7.20 (d, $J$ = 8.52 Hz, 2 H), 7.04 (d, $J$ = 8.77 Hz, 2 H), 4.18 - 4.32 (m, 2 H), 2.99 - 3.14 (m, 3 H), 2.96 (br s, 2 H), 2.81 (tt, $J$ = 12.69, 3.26 Hz, 1 H), 2.14 (br d, $J$ = 13.15 Hz, 1 H), 1.90 - 2.07 (m, 9 H), 1.60 - 1.88 (m, 13 H), 1.44 - 1.56 (m, 2 H), 1.33 - 1.43 (m, 1 H) $^{13}C$ NMR (100 MHz, $CDCl_3$) δ 153.8, 149.7, 142.6, 127.6, 121.6, 111.4, 109.0, 48.7, 42.1, 41.3, 36.8, 36.4, 34.8, 34.8, 34.2, 32.9, 26.9, 26.5, 23.5. MS (ESI) calculated for $C_{27}H_{37}N_2O_5$ $[M + H]^+$ *m/z* 483.29, found 483.24.

(*R*)-**S6: (3*R*)-3-[4-(benzyloxy)phenyl]cyclohexan-1-one.**

To a round bottom flask under intert atmosphere was added acetylacetonatobis(ethylene)-rhodium(I) (160 mg, 0.625 mmol, 0.03 equiv.) and dioxane (100mL). The flask was flushed with argon, then R-BINAP (1.30 g, 2.08 mmol, 0.1 equiv.), Potassium hydroxide (1M solution, 10 mL), and [4-(benzyloxy)phenyl]boronic acid (11.9 g, 52.0 mmol, 2.5 equiv.) were added sequentially. The flask was stirred under Argon for 10 minutes. Cyclohex-2-en-1-one (2.00 g, 20.8 mmol, 1 equiv.) was added and the reaction flask was purged and back-filled with Argon three times. The reaction was heated to 100 °C and stirred overnight. Following consumption of starting material, the reaction was cooled to room temperature, filtered over celite, and concentrated. Purification via flash column chromatography (330 g silica gel cartridge, isocratic 12% EtOAc:hexanes elution) followed by concentration and lyophilization of product fractions yielded (*R*)-**S6** (3.17 g, 11.3 mmol, 54.3%) as a white solid. $^1H$ NMR ($CDCl_3$, 400 MHz) δ 7.3-7.6 (m, 5H), 7.19 (d, 2H, $J$ = 8.5 Hz), 7.00 (d, 2H, $J$ = 8.5 Hz), 5.09 (s, 2H), 3.01 (tt, 1H, $J$ = 3.9, 11.7 Hz), 2.6-2.7 (m, 1H), 2.4-2.6 (m, 3H), 2.0-2.3 (m, 2H), 1.8-2.0 (m, 2H); $^{13}C$ NMR ($CDCl_3$, 100 MHz) δ 211.2, 157.6, 137.1, 136.9, 128.7, 128.0, 127.6, 127.6, 115.0, 70.1, 49.3, 44.0, 41.2, 33.0, 25.6; MS (ESI) calc for $C_{19}H_{21}O_2$ [M+H]+: m/z 281.15 found 281.29.

(*R*)-**S7: (3*R*)-3-(4-hydroxyphenyl)cyclohexan-1-one.**

To a round bottom flask under inert Ar(g) atmosphere was added 10% Pd/C (0.317g, 0.298 mmol, 0.0264 equiv.) and ethyl acetate (50 mL). To this mixture was added (*R*)-**S6** (3.17g, 11.3mmol, 1 equiv.) and the flask was evacuated and purged with Ar(g) three times. The flask was once again evacuated and backfilled with Hydrogen at 1 atm. The solution was heated to 50 °C and stirred for 48 hours. At this point, the reaction was judged complete and the reaction mixture was placed under Ar(g) and filtered over celite, rinsing with ethyl acetate, and the filtrate concentrated and lyophilized to yield (*R*)-**S7** (1.97g, 10.4mmol, 91.6%) as a white powder that was used without further purification. $^{1}$H NMR ($CDCl_3$, 400 MHz) δ 7.10 (d, 2H, *J* = 8.5 Hz), 6.83 (d, 2H, *J* = 8.3 Hz), 2.97 (tdd, 1H, *J* = 3.9, 7.9, 15.5 Hz), 2.4-2.6 (m, 4H), 2.0-2.2 (m, 2H), 1.7-1.9 (m, 2H); $^{13}$C NMR ($CDCl_3$, 100 MHz) δ 212.2, 154.5, 128.6, 127.7, 115.5, 49.3, 44.0, 41.2, 33.0, 25.5; MS (ESI) calc for $C_{12}H_{15}O_2$ [M+H]+: m/z 191.10 found 191.17.

**(*R*)-S1: 4-[(1*R*)-3-oxocyclohexyl]phenyl acetate.**
To a solution of (3R)-3-(4-hydroxyphenyl)cyclohexan-1-one ((*R*)-**S7,** 1.90 g, 9.99 mmol, 1 equiv.) in dichloromethane (50mL) was added triethylamine (2.02 g, 20.0 mmol, 2 equiv.). The solution was cooled to 0C, and acetic anhydride (3.06 g, 30.0 mmol, 3 equiv.) was added dropwise. The solution was allowed to return to room temperature and was stirred for 45 minutes. Following completion the solution was diluted with deionized water and the organic layer was extracted over water, saturated $NaHCO_3$, and brine. The organic fraction was dried over MgSO4, concentrated, and lyophilized to yield (*R*)-**S1** (2.32 g, 9.99 mmol, 100%) as a white solid. Product was confirmed via UPLC/MS and used immediately in the next reaction. MS (ESI) calc for $C_{14}H_{17}O_3$ [M+H]+: m/z 233.11 found 233.23.

**(*R,R*)-S2: *trans*-(*R,R*)-4-(dispiro[adamantane-2,3'-[1,2,4]trioxolane-5',1''-cyclohexan]-3''-yl)phenyl acetate.**
To an oven-dried round bottom flask was added (*R*)-**S1** (1.00g, 4.31mmol, 1 equiv.), adamantan-2-one *O*-methyl oxime (2.00g, 11.2mmol, 2.59 equiv.) and carbon tetrachloride (50mL). The solution was cooled to 0°C and sparged with $O_2$ for 10min. The reaction was maintained at 0C while ozone was bubbled (2L/min, 40% power) through the solution. Following 3.5 hours, the reaction was deemed complete via UPLC/MS and TLC. The reaction mixture was concentrated to yield a crude mixture as a viscous oil. The crude material was purified via flash column chromatography (220 g silica gel cartridge, 5-15% EtOAc:Hex) and product fractions were concentrated and lyophilized to yield (*R,R*)-**S2** (1.146g, 2.876mmol, 66.8%) as a colorless solid. $^{1}$H NMR ($CDCl_3$, 400 MHz) δ 7.22 (d, 2H, *J* = 8.5 Hz), 7.03 (d, 2H, *J* = 8.8 Hz), 2.82 (tt, 1H, *J* = 3.4, 12.8 Hz), 2.31 (s, 3H), 2.1-2.2 (m, 1H), 1.9-2.0 (m, 7H), 1.7-1.9 (m, 12H), 1.5-1.6 (m, 1H), 1.2-1.5 (m, 1H); $^{13}$C NMR ($CDCl_3$, 100 MHz) δ 169.7, 148.9, 143.2, 127.8, 121.4, 111.4, 109.0, 42.2, 41.3, 36.8, 36.4, 34.8, 34.8, 34.2, 32.8, 26.9, 26.5, 23.5, 21.2; MS (ESI) calc for $C_{24}H_{30}O_5Na$ [M+Na]+: m/z 421.20 found 421.29.

**(*R,R*)-S3: *trans*-(*R,R*)-4-(dispiro[adamantane-2,3'-[1,2,4]trioxolane-5',1''-cyclohexan]-3''-yl)phenol.**
To an oven-dried round bottom flask was added (*R,R*)-**S2** (1.10 g, 2.760 mmol, 1 equiv.) in tetrahydrofuran (20mL) and methanol (20mL). Lithium hydroxide (0.264 g, 11.04 mmol, 4 equiv.) was added as a 1M aqueous solution and the reaction was stirred overnight at room temperature. The reaction was quenched with 1M $NH_4Cl$ and the aqueous layer was extracted 3 times with $CH_2Cl_2$. The combined organic fractions were dried over $MgSO_4$, concentrated, and lyophilized to yield (*R,R*)-**S3** (0.984 g, 2.76 mmol, 84.8%) as a white solid that was used in the next reaction without further purification. $^{1}$H NMR ($CDCl_3$, 400 MHz) δ 7.09 (br d, 2H, *J* = 8.3 Hz), 6.79 (br d, 2H, *J* = 8.5 Hz), 2.7-2.8 (m, 1H), 2.12 (br d, 4H, *J* = 12.4 Hz), 1.9-2.0 (m, 7H), 1.7-1.9 (m, 8H), 1.5-1.7 (m, 2H), 1.2-1.5 (m, 2H); $^{13}$C NMR ($CDCl_3$, 100 MHz) δ 153.9, 138.0, 127.9, 115.2, 111.4, 109.2, 100.0, 42.4, 41.0, 36.8, 36.4, 34.8, 34.8, 34.2, 33.0, 26.9, 26.5, 23.6; MS (ESI) calc for $C_{22}H_{29}O_4$ [M-H]-: m/z 355.19 found 355.26.

**(*R,R*)-S4: *trans*-(*R,R*)-4-(dispiro[adamantane-2,3'-[1,2,4]trioxolane-5',1''-cyclohexan]-3''-yl)phenyl (4-nitrophenyl) carbonate.**

To an oven-dried round bottom flask was added (*R,R*)-**S3** (0.834 g, 2.34 mmol, 1 equiv.) and dichloromethane (50 mL). The solution was cooled to 0°C, and DIEA (0.907 g, 7.02 mmol, 3 equiv.) and DMAP (0.057 g, 0.468 mmol, 0.2 equiv.) were added. The solution was stirred at 0 °C for 10 minutes, following which *p*-nitrophenyl chloroformate (1.41 g, 7.02 mmol, 3 equiv.) was added portionwise. Upon complete addition the reaction was allowed to return to room temperature and stirred overnight. The reaction was diluted with dichloromethane and 1M $Na_2CO_3$, and the organic layer was extracted repeatedly with 1M $Na_2CO_3$ until no yellow color was observed, indicating removal of *p*-nitrophenol. The organic layer was dried over $MgSO_4$ and concentrated to yield a crude colorless oil. This crude product was purified via flash column chromatography (80g silica gel cartridge, 0-25% EtOAC:Hex elution) and the product fractions were combined, concentrated, and lyophilized to afford (*R,R*)-**S4** (0.310 g, 2.34 mmol, 25%) as a white powder. $^1$H NMR (DMSO-d6, 400 MHz) δ 8.37 (d, 1H, *J* = 9.3 Hz), 7.71 (d, 1H, *J* = 9.0 Hz), 7.3-7.4 (m, 4H), 2.5-2.8 (m, 1H), 1.8-2.1 (m, 8H), 1.7-1.8 (m, 9H), 1.6-1.7 (m, 4H), 1.3-1.6 (m, 3H); $^{13}$C NMR (DMSO-d6, 100 MHz) δ 155.6, 151.3, 149.4, 145.9, 144.3, 128.5, 126.0, 123.2, 121.5, 111.1, 109.2, 41.7, 41.3, 38.9, 36.6, 36.3, 34.7, 33.9, 32.6, 26.7, 26.3, 23.6; MS (ESI) calc for $C_{29}H_{31}NO_8Na$ [M+Na]+: m/z 544.19 found 543.94.

(*R,R*)-**S5: *trans*-(*R,R*)-4-(dispiro[adamantane-2,3'-[1,2,4]trioxolane-5',1''-cyclohexan]-3''-yl)phenyl 4-((*tert*-butoxycarbonyl)amino)piperidine-1-carboxylate.**

To an oven-dried round bottom flask was added (*R,R*)-**S4** (0.300 g, 0.575 mmol, 1 equiv.) followed by tert-butyl piperidin-4-ylcarbamate (0.184 g, 0.920 mmol, 1.6 equiv.) and DMAP (0.014 g, 0.115 mmol, 0.2 equiv.). The reaction flask was evacuated and purged with argon. DMF (35mL) and DIEA (0.223 mg, 1.73 mmol, 3 equiv.) were added and the reaction mixture was stirred at room temperature overnight. The reaction was judged complete at this stage, and the mixture was diluted with ethyl acetate and the organic layer was extracted with 1M $Na_2CO_3$ until no yellow color was observed, indicating removal of p-nitrophenol. The organic layer was dried over $MgSO_4$ and concentrated to yield a crude colorless oil. The crude product was purified via flash column chromatography (40 g silica gel cartridge, 0-25% EtOAc:hexanes) to yield (*R,R*)-**S5** (0.226 g, 0.389 mmol, 67.5%) as a white powder. The identify of the product was confirmed via UPLC/MS analysis and used immediately in the next reaction. MS (ESI) calc for $C_{33}H_{46}N_2O_7Na$ [M+Na]+: m/z 605.32 found 605.52

(*R,R*)-**2 (RLA-5763): *trans*-(*R,R*)-4-(dispiro[adamantane-2,3'-[1,2,4]trioxolane-5',1''-cyclohexan]-3''-yl)phenyl 4-aminopiperidine-1-carboxylate.**

A solution of (*R,R*)-**S5** (290 mg, 0.498 mmol, 1.0 equiv.) in methanol (5 mL) and THF (5 mL) was cooled to 0 °C using an ice bath. After purging the flask with argon, acetyl chloride (389 mg, 4.98 mmol, 10 equiv.) was added dropwise. The solution was stirred at 0 °C for 15 minutes, then allowed to return to room temperature and stirred for 16 hours. The reaction was diluted with dichloromethane, treated with 1M $Na_2CO_3$, and the aqueous layer was extracted thrice with dichloromethane. The combined organic fractions were filtered, dried over $MgSO_4$, and concentrated to yield a crude product as a light-yellow oil. This material was purified in batches by HPLC (Agilent 1200 Series) using a 5 uM Cellulose-3 column (Phenomenex) with an isocratic mobile phase consisting of 40:60 20mM ammonium bicarbonate pH 8.5 in acetonitrile (with 0.1% diethylamine). Product peaks from each batch were combined, concentrated, and lyophilized to yield (*R,R*)-**2** (85.8 mg, 0.178 mmol, 36%) as a colorless solid. $^1$H NMR (DMSO-d6, 400 MHz) δ 7.27 (d, 2H, *J* = 8.5 Hz), 7.02 (d, 2H, *J* = 8.5 Hz), 3.9-4.2 (m, 2H), 3.1-3.3 (m, 1H), 3.10 (br s, 1H), 2.94 (br s, 1H), 2.5-2.8 (m, 2H), 1.8-2.0 (m, 8H), 1.6-1.8 (m, 12H), 1.4-1.5 (m, 3H); $^{13}$C NMR (DMSO-d6, 100 MHz) δ 153.5, 149.9, 142.8, 128.0, 122.2, 111.0, 109.2, 47.7, 41.8, 41.3, 36.5, 36.2, 34.7, 33.9, 32.7, 26.7, 26.3, 23.7; MS (ESI) calc for $C_{28}H_{39}N_2O_5$ [M+H]+: m/z 483.28 found 483.51.

(*S*)-**S6: (3*S*)-3-[4-(benzyloxy)phenyl]cyclohexan-1-one.**

To a round bottom flask under argon was added acetylacetonatobis(ethylene)-rhodium(**I**) (160 mg, 0.625 mmol, 0.03 equiv.) and dioxane (100mL). The flask was flushed with argon, and charged sequentially with S-BINAP (1.30 g, 2.08 mmol, 0.1equiv.), potassium hydroxide (1M solution, 10mL), and [4-(benzyloxy)phenyl]boronic acid (11.9 g, 52.0 mmol, 2.5 equiv.). The flask was stirred under argon for 10 minutes. Cyclohex-2-en-1-one (2.00 g, 20.8 mmol, 1 equiv.) was then added and the reaction flask purged and back-filled with argon three times. The reaction flask was then heated to 100 °C with stirring overnight. Following consumption of starting material, the reaction mixture was cooled to room temperature, filtered over celite, and concentrated. Purification via flash column chromatography (330 g silica gel cartridge, isocratic elution with 12% EtOAc:hexanes) followed by concentration and lyophilization of product fractions yielded (*S*)-**S6** (1.948 g, 6.95 mmol, 33.4%) as a white solid. $^1$H NMR ($CDCl_3$, 400 MHz) δ 7.4-7.6 (m, 5H), 7.19 (d, 2H, *J* = 8.5 Hz), 7.00 (d, 2H, *J* = 8.8 Hz), 5.09 (s, 2H), 3.00 (tt, 1H, *J* = 3.8, 11.7 Hz), 2.6-2.7 (m, 1H), 2.4-2.6 (m, 3H), 2.0-2.3 (m, 2H), 1.7-2.0 (m, 2H)**;** $^{13}$C NMR ($CDCl_3$, 100 MHz) δ 211.2, 157.6, 137.1, 136.9, 128.7, 128.0, 127.6, 127.5, 115.0, 70.1, 49.3, 44.0, 41.2, 33.0, 25.6; MS (ESI) calc for $C_{19}H_{21}O_2$ [M+H]+: m/z 281.15 found 281.24.

(*S*)-**S7: (3*S*)-3-(4-hydroxyphenyl)cyclohexan-1-one.**
To a round bottom flask under inert argon atmosphere was added 10% palladium on carbon (0.195 g, 0.183 mmol, 0.0264 equiv.) in ethyl acetate (50mL). To this mixture, (*S*)-**S6** (1.948 g, 6.948 mmol, 1 equiv.) was added and the flask was evacuated and purged with argon three times. The flask was once again evacuated and backfilled with hydrogen gas to 1atm. The solution was heated to 50 °C and stirred for 48 hours. Following completion of the reaction, the mixture was purged with argon and filtered over celite, rinsing with ethyl acetate, and the combined filtrate was then concentrated and lyophilized to yield (*S*)-**S7** (1.15 g, 6.04 mmol, 87.0%) as a white powder that was used in the next step without further purification. $^1$H NMR ($CDCl_3$, 400 MHz) δ 7.09 (d, 2H, *J* = 8.3 Hz), 6.83 (d, 2H, *J* = 8.8 Hz), 2.97 (dt, 1H, *J* = 3.9, 11.6 Hz), 2.4-2.6 (m, 4H), 2.0-2.2 (m, 2H), 1.7-1.9 (m, 2H)**;** $^{13}$C NMR ($CDCl_3$, 100 MHz) δ 212.3, 154.5, 136.4, 127.7, 115.5, 49.3, 44.0, 41.2, 33.0, 25.5; MS (ESI) calc for $C_{12}H_{15}O_2$ [M+H]+: m/z 191.10 found 191.22.

(*S*)-**S1: 4-[(1*S*)-3-oxocyclohexyl]phenyl acetate.**
To a solution of (*S*)-**S7** (1.10 g, 5.78 mmol, 1 equiv.) in dichloromethane (50 mL) was added triethylamine (1.17 g, 11.6 mmol, 2 equiv.). The solution was cooled to 0 °C, and acetic anhydride (1.77 g, 17.3 mmol, 3 equiv.) was added dropwise. The solution was allowed to return to room temperature and was stirred for 45 min. Following completion of the reaction, the solution was diluted with water and the organic layer was extracted with water, saturated $NaHCO_3$, and brine. The organic fraction was dried over $MgSO_4$, concentrated, and lyophilized to yield (*S*)-**S1** (2.32g, 9.99mmol, 100% crude yield) as a white solid. The identify of the material was confirmed via UPLC/MS and used immediately in the next reaction. MS (ESI) calc for $C_{14}H_{17}O_3$ [M+H]+: m/z 233.11 found 233.23.

(*S,S*)-**S2: *trans*-(*S,S*)-4-(dispiro[adamantane-2,3'-[1,2,4]trioxolane-5',1''-cyclohexan]-3''-yl)phenyl acetate.**
To an oven-dried round bottom flask was added (*S*)-**S1** (1.00 g, 4.31 mmol, 1 equiv.), adamantan-2-one *O*-methyl oxime (2.00 g, 11.2 mmol, 2.59 equiv.) and carbon tetrachloride (50 mL). The solution was cooled to 0 °C and sparged with $O_2$ for 10min. The reaction was maintained at 0 °C while ozone was bubbled (2L/min, 40% power) through the solution. After 3.5 hours, the reaction was deemed complete via UPLC/MS and TLC analysis. The reaction mixture was concentrated to yield the crude product as a viscous oil. This residue was purified via flash column chromatography (220 g silica gel cartridge, 5-15% EtOAc:hexanes) and product-bearing fractions were concentrated and lyophilized to yield **(***S,S*)-**S2** (0.714 g, 4.31 mmol, 41.6%) as a colorless solid that by $^1$H NMR analysis contained a small amount of starting material. $^1$H NMR ($CDCl_3$, 400 MHz) δ 7.23 (d, 2H, *J* = 7.5 Hz), 7.02 (d, 2H, *J* = 8.8 Hz), 2.40 (br s, 5H), 2.31 (s, 3H), 1.9-2.2 (m, 29H), 1.8-1.9 (m, 16H), 1.3-1.5 (m, 4H); $^{13}$C NMR ($CDCl_3$, 100 MHz) δ 169.7, 148.9, 127.7, 121.4, 111.4, 109.2, 109.0, 47.0, 42.2, 41.3, 39.3, 37.3, 36.8, 36.6, 36.4, 35.8, 35.0, 34.8, 34.8, 34.8, 34.2, 34.0, 33.8, 33.6, 33.2, 32.8, 32.3, 31.5, 31.0, 27.5, 27.2, 27.1, 27.0, 26.9, 26.5, 23.5, 21.2; MS (ESI) calc for $C_{24}H_{30}O_5Na$ [M+Na]+: m/z 421.20 found 421.19.

(*S,S*)-**S3: *trans*-(*S,S*)-4-(dispiro[adamantane-2,3'-[1,2,4]trioxolane-5',1''-cyclohexan]-3''-yl)phenol.**

To an oven-dried round bottom flask was added (*S,S*)-**S2** (0.700 g, 1.76 mmol, 1 equiv.) in tetrahydrofuran (20 mL) and methanol (20 mL). Lithium hydroxide (0.168 g, 7.03 mmol, 4 equiv.) was added as a 1M solution in water and the reaction was stirred overnight at room temperature. The reaction was quenched with 1M $NH_4Cl$ and the aqueous layer was extracted thrice with $CH_2Cl_2$. The combined organic fractions were dried over $MgSO_4$, concentrated, and lyophilized to yield (*S,S*)-**S3** (0.560 g, 1.57 mmol, 89.4%) as a white solid. $^1H$ NMR ($CDCl_3$, 400 MHz) δ 7.09 (br d, 2H, *J* = 8.3 Hz), 6.79 (br d, 2H, *J* = 8.0 Hz), 2.75 (br t, 1H, *J* = 12.7 Hz), 2.12 (br d, 2H, *J* = 12.7 Hz), 1.9-2.1 (m, 9H), 1.7-1.9 (m, 9H), 1.5-1.7 (m, 3H), 1.2-1.5 (m, 2H); $^{13}C$ NMR ($CDCl_3$, 100 MHz) δ 153.9, 138.1, 127.9, 115.2, 111.4, 109.2, 42.4, 41.0, 39.3, 36.8, 36.4, 34.8, 34.8, 34.2, 33.0, 26.9, 26.5, 23.6; MS (ESI) calc for $C_{22}H_{29}O_4$ [M-H]-: m/z 355.19 found 355.31.

(*S,S*)-**S4: *trans*-(*S,S*)-4-(dispiro[adamantane-2,3'-[1,2,4]trioxolane-5',1''-cyclohexan]-3''-yl)phenyl (4-nitrophenyl) carbonate.**

To an oven-dried round bottom flask was added (*S,S*)-**S3** (0.560 g, 1.57 mmol, 1 equiv.) and dichloromethane (50mL). The solution was cooled to 0C, and DIEA (0.609 g, 4.71 mmol, 3 equiv.) and DMAP (0.038 g, 0.314 mmol, 0.2 equiv.) were added. The solution was stirred at 0 °C for 10 minutes, following which *p*-(nitrophenyl)chloroformate (1.41g, 7.02mmol, 3 equiv.) was added portionwise. Upon complete addition the reaction was allowed to return to room temperature and stirred overnight. The reaction was diluted with dichloromethane and 1M $Na_2CO_3$, and the organic layer was extracted repeatedly with 1M $Na_2CO_3$ until no yellow color was observed, indicating removal of p-nitrophenol. The organic layer was dried over $MgSO_4$ and concentrated to yield a crude colorless oil. This crude material was purified via flash column chromatography (80 g silica gel cartridge, 0-25% gradient elution with EtOAC:hexanes) and the product-bearing fractions combined, concentrated, and lyophilized to yield (*S,S*)-**S4** (0.339 g, 0.65 mmol, 41.4%) as a white powder. $^1H$ NMR (DMSO-d6, 400 MHz) δ 8.37 (d, 2H, *J* = 9.0 Hz), 7.71 (d, 2H, *J* = 9.0 Hz), 7.3-7.4 (m, 4H), 2.5-2.8 (m, 1H), 1.9-2.0 (m, 7H), 1.6-1.8 (m, 13H), 1.3-1.6 (m, 3H); $^{13}C$ NMR (DMSO-d6, 100 MHz) δ 155.6, 151.3, 149.4, 145.9, 144.3, 128.5, 126.0, 123.2, 121.5, 111.1, 109.2, 41.7, 41.3, 38.9, 36.5, 36.3, 34.7, 33.9, 32.6, 26.7, 26.3, 23.6; MS (ESI) calc for $C_{29}H_{31}NO_8Na$ [M+Na]+: m/z 544.19 found 544.14.

(*S,S*)-**S5: *trans*-(*S,S*)-4-(dispiro[adamantane-2,3'-[1,2,4]trioxolane-5',1''-cyclohexan]-3''-yl)phenyl 4-((*tert*-butoxycarbonyl)amino)piperidine-1-carboxylate**

To an oven-dried round bottom flask was added (*S,S*)-**S4** (0.320 g, 0.614 mmol, 1 equiv.) followed by tert-butyl piperidin-4-ylcarbamate (0.197 g, 0.982 mmol, 1.6 equiv.) and DMAP (0.015 g, 0.123 mmol, 0.2 equiv.). The reaction flask was evacuated and purged with argon. DMF (35mL) and DIEA (0.238 mg, 1.84 mmol, 3 equiv.) were added and the reaction was stirred at room temperature overnight. Following completion, the reaction was diluted with ethyl acetate and the organic layer was extracted with 1M $Na_2CO_3$ until no yellow color was observed, indicating removal of p-nitrophenol. The organic layer was dried over $MgSO_4$ and concentrated to yield a crude colorless oil. The crude product was purified via flash column chromatography (40 g silica gel cartridge, gradient elution in 0-25% EtOAc:hexanes) to yield (*S,S*)-**S5** (0.315 g, 0.540 mmol, 88.0%) as a white powder. The identity of the product was confirmed via UPLC/MS analysis and the material used immediately in the next reaction. MS (ESI) calc for $C_{33}H_{46}N_2O_7Na$ [M+Na]+: m/z 605.32 found 605.32.

(*S,S*)-**2 (RLA-5764): *trans*-(*S,S*)-4-(dispiro[adamantane-2,3'-[1,2,4]trioxolane-5',1''-cyclohexan]-3''-yl)phenyl 4-aminopiperidine-1-carboxylate.**

A solution of (*S,S*)-**S5** (300 mg, 0.515 mmol, 1.0 equiv.) in methanol (5 mL) and THF (5 mL) was cooled to 0 °C in an ice bath. After purging the flask with argon, acetyl chloride (402.6 mg, 5.15 mmol, 10 equiv.) was added dropwise. The solution was stirred at 0 °C for 15 minutes, then allowed to return to room temperature and stirred for 16 hours. The reaction mixture was diluted with dichloromethane, quenched with 1M $Na_2CO_3$, and the aqueous layer was extracted thirce with dichloromethane. The combined organic fractions were filtered, dried over $MgSO_4$, and concentrated to yield a crude product as a light-yellow oil. The crude residue was purified in batches by HPLC (Agilent 1200 Series) using a 5 uM Cellulose-3 column (Phenomenex) with an isocratic mobile phase consisting of 40:60 20 mM ammonium bicarbonate pH 8.5 in acetonitrile (with 0.1% diethylamine). Product peaks from each batch were combined, concentrated, and lyophilized to yield (*S,S*)-**2** (150.3 mg, 0.311mmol, 61%) as a colorless solid. $^{1}H$ NMR (DMSO-d6, 400 MHz) δ 7.26 (d, 2H, *J* = 8.3 Hz), 7.01 (d, 2H, *J* = 8.3 Hz), 3.8-4.2 (m, 2H), 2.9-3.1 (m, 2H), 2.8-2.9 (m, 1H), 2.6-2.8 (m, 1H), 1.8-2.0 (m, 7H), 1.6-1.8 (m, 15H), 1.3-1.6 (m, 3H); $^{13}C$ NMR (DMSO-d6, 100 MHz) δ 153.5, 150.0, 142.6, 127.9, 122.2, 111.0, 109.2, 48.1, 41.8, 41.3, 36.5, 36.2, 34.7, 33.9, 32.7, 26.7, 26.3, 23.7; MS (ESI) calc for $C_{28}H_{39}N_2O_5$ [M+H]+: m/z 483.28 found 483.56.

(±)-**3 (RLA-3107)**: ***trans*-4-(2-(4-(dispiro[adamantane-2,3'-[1,2,4]trioxolane-5',1''-cyclohexan]-3''-yl)phenoxy)ethyl)morpholine**.
To a solution of (±)-**S3** (100 mg, 0.281 mmol, 1.0 equiv) in dry $CH_3CN$ (5 mL) was added tetrabutylammonium hydrogen sulfate (19.1 mg, 0.056 mmol, 0.2 equiv) and powdered NaOH (44.9 mg, 1.122 mmol, 4.0 equiv). This mixture was then allowed to stir at room temperature for 30 minutes, at which point 4-(2-chloroethyl)morpholine hydrochloride (104.4 mg, 0.561 mmol, 2.0 equiv) was added to the solution. The mixture was then placed in an oil bath preheated to 55 ˚C, and was allowed to stir at this temperature for 14.5 hours after which the reaction was judged complete. After cooling, the mixture was diluted with EtOAc (20 mL) and $H_2O$ (10 mL), which served to dissolve all of the inorganic solids present. Following separation of the layers, the aqueous layer was extracted with EtOAc (20 mL) which resulted in the formation of an emulsion, to which brine (10 mL) was added to aid in separation of the organic phase. The combined organic layers were then washed with brine (10 mL), dried over anhydrous $Na_2SO_4$, filtered, and concentrated under reduced pressure. The residue was then purified via flash column chromatography (25 g silica gel cartridge, 0–100% EtOAc/hexanes) to yield (±)-**3** (82 mg, 62%) as a clear colorless oil that was a single diastereomer by $^{1}H$ NMR analysis. $^{1}H$ NMR (400 MHz, $CDCl_3$) δ 7.11 (d, *J* = 8.6 Hz, 2H), 6.84 (d, *J* = 8.6 Hz, 2H), 4.09 (t, *J* = 5.7 Hz, 2H), 3.74 (t, *J* = 4.7 Hz, 4H), 2.79 (t, *J* = 5.7 Hz, 2H), 2.79–2.68 (m, 1H), 2.58 (t, *J* = 4.5 Hz, 4H), 2.15–2.06 (m, 1H), 2.04–1.87 (m, 7H), 1.87–1.54 (m, 13H), 1.40–1.24 (m, 1H); $^{13}C$ NMR (100 MHz, $CDCl_3$) δ 157.0, 138.1, 127.6, 114.5, 111.3, 109.0, 66.8, 65.7, 57.6, 54.0, 42.3, 41.0, 36.7, 36.4, 34.8, 34.7, 34.1, 32.9, 26.8, 26.4, 23.5; MS (ESI) calculated for $C_{28}H_{40}NO_5$ $[M + H]^+$ *m/z* 470.29, found 470.09.

(±)-**S9: 3-(3-hydroxyphenyl)cyclohexan-1-one.**
To an oven-dried round bottom flask containing a magnetic stir bar under an Ar(g) atmosphere was added palladium(II) acetylacetonate (0.158 g, 0.52 mmol, 0.05 equiv), 1,2-bis(diphenylphosphino)-benzene (0.232 g, 0.520 mmol, 0.05 equiv), copper(II)tetrafluoroborate hydrate (0.531 g, 2.081 mmol, 0.2 equiv), and 3-hydroxyphenylboronic acid (2.511 g, 18.21 mmol, 1.75 equiv). To the mixture of solid materials was added anhydrous dimethoxyethane (60 mL). To the stirring solution was then added 2-cyclohexen-1-one (1.0 g, 10.33 mmol, 1.0 equiv) via syringe at room temperature. The solution was allowed to stir at room temperature for 20 hours. Based on LCMS and TLC analysis, it was determined that the reaction was complete. The mixture was then concentrated under reduced pressure to yield a dark green oil. To this oil was added EtOAc (100 mL) followed by $H_2O$ (50 mL). Following separation of the layers, the organic layer was washed with additional $H_2O$ (50 mL). The organic layer was then filtered through a pad of Celite to remove all insoluble inorganic material. The pad was then rinsed with EtOAc (50 mL x 4). The aqueous layer was extracted with EtOAc, and the resulting organic solution was filtered through the same pad of Celite. The pad was then again rinsed with EtOAc (50 mL x 2). The clear yellow solution was concentrated under reduced pressure to yield a yellow oil. The residue was then purified by flash column chromatography (120 g silica gel cartridge, product eluted during 35–40% EtOAc/Hex) to yield (±)-**S9** (1.44 g, 73%) as a colorless oil. $^{1}H$ NMR (400 MHz, $CDCl_3$) δ 7.33 (s, 1H), 7.16 (t, *J* = 8.04 Hz, 1H), 6.72 - 6.77 (m, 3H), 2.88 - 2.97 (m, 1H), 2.43 - 2.61 (m, 3H), 2.32 - 2.41 (m, 1H), 2.08 - 2.15 (m, 1H), 2.02 (br d, *J* = 11.69 Hz, 1H), 1.67 - 1.84 (m, 2H); $^{13}C$ NMR (100 MHz, $CDCl_3$) δ 213.6, 156.3, 146.0, 129.8, 118.5, 113.8, 113.7, 48.7, 44.6, 41.1, 32.5, 25.4; MS (ESI) calculated for $C_{12}H_{13}O_2$ $[M - H]^-$ *m/z* 189.09, found 188.99.

(±)-**S10: 3-(3-oxocyclohexyl)phenyl acetate.**
To an oven-dried round bottom flask containing a magnetic stir bar under an Ar(g) atmosphere was added (±)-**S9** (0.525 g, 2.76 mmol, 1.0 equiv), dichloromethane (7 mL), acetic anhydride (0.29 mL, 3.04 mmol, 1.1 equiv), and 4-dimethylaminopyridine (0.337 g, 2.76 mmol, 1 equiv). The solution

was allowed to stir at room temperature for 2 hours. The reaction was then diluted with $H_2O$ (100 mL) and subsequently extracted with EtOAc (100 mL). The organic layer was dried over anhydrous $Na_2SO_4$, filtered, and concentrated under reduced pressure to yield a thick yellow oil. The residue was then purified by flash column chromatography (80 g silica gel cartridge, 0–25% EtOAc/hexanes, product eluted during 20% EtOAc/Hex) to yield (±)-**S10** (503 mg, 78%) as a white solid. $^1H$ NMR (400 MHz, $CDCl_3$) δ 7.15 - 7.24 (m, 1H), 6.97 - 7.02 (m, 1H), 6.87 - 6.92 (m, 2H), 2.85 - 2.94 (m, 1H), 2.40 - 2.51 (m, 2H), 2.23 - 2.35 (m, 2H), 2.17 - 2.20 (m, 3H), 1.90 - 2.02 (m, 2H), 1.57 - 1.75 (m, 2H); $^{13}C$ NMR (100 MHz, $CDCl_3$) 210.5, 169.0, 150.5, 145.7, 129.2, 123.7, 119.4, 119.4, 48.1, 43.8, 40.6, 32.0, 24.9, 20.6; MS (ESI) calculated for $C_{14}H_{16}NaO_3$ $[M + Na]^+$ *m/z* 255.10, found 255.04.

(±)-**S11:** ***trans*-3-(dispiro[adamantane-2,3'-[1,2,4]trioxolane-5',1''-cyclohexan]-3''-yl)phenyl acetate.**

A solution of adamantan-2-one *O*-methyl oxime (772 mg, 4.31 mmol, 2.0 equiv) and 3-(3-oxocyclohexyl)phenyl acetate ((±)-**S10,** 500 mg, 2.15 mmol, 1.0 equiv) in 50 mL carbon tetrachloride was cooled to 0 ˚C and subsequently sparged with $O_2$ for 10 minutes. The reaction was kept at 0 ˚C while ozone was then bubbled (2 L/min, 35% power). After stirring for 2 hrs, the reaction was deemed to be incomplete based on LCMS analysis and additional oxime (193 mg, 1.08 mmol, 0.5 equiv) was added in a single portion to the reaction. Ozone was bubbled through the reaction for another 60 mins, at which point the reaction was found to be complete. The reaction was then purged with $O_2$ for 10 minutes to remove any dissolved ozone, followed by sparging with argon gas for 10 minutes to remove any dissolved oxygen. The solution was then concentrated under reduced pressure to provide an extremely viscous oil. The residue was purified through by column chromatography (80 g silica gel cartridge, 0–10% EtOAc/Hexanes, product eluted during 8% EtOAc/Hex) to yield (±)-**S11** (724 mg, 84%) as a thick colorless oil, which solidified to a white solid under high vacuum. The material was determined to a 7.4:1 mixture in favor of the *trans* diastereomer. $^1H$ NMR (400 MHz, $CDCl_3$) δ 7.28 - 7.35 (m, 1H), 7.10 (d, *J* = 7.79 Hz, 1H), 6.93 - 6.96 (m, 2H), 2.98 (tt, *J* = 12.7, 3.1 Hz, 1H, minor diastereomer), 2.83 (tt, *J* = 12.8, 3.3 Hz, 1H), 2.30 - 2.32 (m, 3H), 2.14 - 2.20 (m, 1H), 2.07 - 2.12 (m, 1H), 1.90 - 2.05 (m, 7H), 1.61 - 1.89 (m, 12H), 1.39 (dq, J = 3.17, 12.42 Hz, 1H); $^{13}C$ NMR (100 MHz, $CDCl_3$) δ 169.5, 169.5 (minor diastereomer), 150.9, 147.6 (minor diastereomer), 147.4, 129.4, 124.4, 124.3 (minor diastereomer), 120.0 (minor diastereomer), 120.0, 119.5, 111.9 (minor diastereomer), 111.4, 109.0 (minor diastereomer), 108.9, 47.0, 42.0, 41.7, 41.5, 41.4, 39.3, 36.8, 36.5, 36.5, 36.4, 35.1, 35.0, 34.8, 34.8, 34.2, 34.1, 33.4, 32.6, 27.5, 27.0, 26.9, 26.5, 23.5, 21.2; MS (ESI) calculated for $C_{24}H_{30}O_5Na$ $[M + Na]^+$ *m/z* 421.20, found 421.14.

(±)-**S12:** ***trans*-3-(dispiro[adamantane-2,3'-[1,2,4]trioxolane-5',1''-cyclohexan]-3''-yl)phenol.**

To a solution of *trans*-3-(dispiro[adamantane-2,3'-[1,2,4]trioxolane-5',1''-cyclohexan]-3''-yl)phenyl acetate ((±)-**S11**, 724 mg, 1.0 equiv) in anhydrous THF (7 mL) and MeOH (14 mL) was added 15% aqueous KOH solution (3 mL, 4.5 equiv). The reaction mixture was then placed in an oil bath that had been preheated to 50 °C, and was allowed to stir at this temperature for 1.5 hours, and the solution then allowed to cool to room temperature. The mixture was then concentrated under reduced pressure, to this was then added $H_2O$ (30 mL). Citric acid was then added to this solution until pH ~ 4, and extracted with EtOAc (100 mL). The organic layer was then washed with brine, dried over anhydrous $Na_2SO_4$, filtered, and concentrated under reduced pressure, the residue was then purified by flash column chromatography (80 g silica gel cartridge, 0–50% EtOAc/Hexanes, product eluted during 30%) to yield (±)-**S12** (550 mg, 85%) as a thick colorless oil, which solidified to a white solid under high vacuum. This material was determined to be an 8.2:1 mixture in favor of the *trans* diastereomer. $^1H$ NMR (400 MHz, $CDCl_3$) δ δ 7.14 - 7.18 (m, 1H), 6.76 - 6.79 (m, 1H), 6.67 - 6.71 (m, 2H), 5.38 (br s, 1H), 2.98 (tt, *J* = 12.7, 3.1 Hz, 1H, minor diastereomer), 2.83 (tt, *J* = 12.8, 3.3 Hz, 1H), 2.10 - 2.16 (m, 1H), 1.89 - 2.04 (m, 7H), 1.65 - 1.86 (m, 12H), 1.58 - 1.65 (m, 1H), 1.30 - 1.41 (m, 1H); $^{13}C$ NMR (100 MHz, $CDCl_3$) δ 155.8 (minor diastereomer), 155.8, 147.9 (minor diastereomer), 147.7, 129.7, 119.3, 113.9, 113.3, 113.3 (minor diastereomer), 112.0 (minor diastereomer), 111.6, 109.3, 42.1, 41.8, 41.6, 41.6, 36.9, 36.5, 35.1, 35.0, 34.9, 34.9, 34.8, 34.3, 34.2, 33.4, 32.7, 27.0, 27.0, 26.6, 23.6; MS (ESI) calculated for $C_{22}H_{27}O_4$ $[M – H]^–$ *m/z* 355.19, found 355.13.

(±)-**4 (RLA-4741):** ***trans*-4-(2-(3-(dispiro[adamantane-2,3'-[1,2,4]trioxolane-5',1''-cyclohexan]-3''-yl)phenoxy)ethyl)morpholine. (RLA-4741).**

To a solution of (±)-**S12** (60.0 mg, 0.17 mmol, 1.0 equiv) in dry $CH_3CN$ (4 mL) was added tetrabutylammonium hydrogen sulfate (16.0 mg, 0.05 mmol, 0.3 equiv) and powdered NaOH (24.0 mg, 0.60 mmol, 3.6 equiv). This mixture was then allowed to stir at room temperature for 30 minutes, at which point 4-(2-chloroethyl)morpholine hydrochloride (63.0 mg, 0.34 mmol, 2.0 equiv) was added to the solution. The mixture was then placed in an oil bath that had been preheated to 55 ˚C, and was allowed to stir at this temperature for 14 hours. Following determination of reaction completion by TLC and LCMS analysis, the solution was then removed from the oil bath and allowed to cool to room temperature. Upon cooling, the mixture was diluted with EtOAc (20 mL) prior to the addition of $H_2O$ (10 mL), which served to dissolve all of the inorganic solids present. Following separation of the layers, the aqueous layer was extracted with EtOAc (20 mL). The organic layer was then washed with brine (10 mL), dried over anhydrous $Na_2SO_4$, filtered, and concentrated under reduced pressure. The residue was then purified through flash column chromatography (12 g silica gel cartridge, 0–50% EtOAc/hexanes, followed by 0–20% $MeOH/CH_2Cl_2$, and desired product eluted during 2% MeOH, to yield (±)-**4** (48.0 mg, 61%) as a light yellow oil. This material was found to be solely the *trans* diastereomer within limits of NMR detection. $^1H$ NMR (400 MHz, $CDCl_3$) δ 7.18 - 7.24 (m, 1H), 6.72 - 6.83 (m, 3H), 4.09 - 4.14 (m, 2H), 3.71 - 3.79 (m, 4H), 2.73 - 2.85 (m, 3H), 2.56 - 2.65 (m, 4H), 1.89 - 2.15 (m, 9H), 1.78 - 1.87 (m, 5H), 1.65 - 1.75 (m, 6H), 1.56 - 1.65 (m, 1H), 1.32 - 1.44 (m, 1H); $^{13}C$ NMR (100 MHz, $CDCl_3$) δ 158.8, 147.4, 129.5, 119.4, 113.4, 112.1, 111.4, 109.0, 73.2, 66.9, 65.6, 57.7, 54.1, 42.1, 42.0, 41.3, 36.8, 36.5, 35.8, 34.9, 34.8, 34.3, 32.7, 31.0, 26.9, 26.5, 25.9, 23.6; MS (ESI) calculated for $C_{28}H_{40}NO_5$ $[M + H]^+$ *m/z* 470.29, found 470.22.

(±)-**S14: 2-(3-oxocyclohexyl)phenyl acetate.**

To a suspension of 2-(3-oxocyclohexyl)phenol ((±)-**S13,** 2.7 g, 14.3 mmol, 1 equiv.) in 50 mL $CH_2Cl_2$ was added pyridine (5.8 mL, 71.5 mmol, 5 equiv.) and acetic anhydride (2.7 mL, 28.6 mmol, 1 equiv.) and the reaction mixture stirred at room temperature for 16 h. The mixture was then quenched by the addition of 1 M HCl aq. and extracted with $CH_2Cl_2$ three times. The resulting organic layers were combined, dried over $MgSO_4$ and concentrated. The residue was then purified by flash column chromatography (120 g silica gel cartridge, 0–100 % EtOAc/hexanes) to yield (±)-**S14** (3.2 g, 95 %) as a white solid. $^1H$ NMR (400 MHz, $CDCl_3$) δ 7.35 – 7.28 (m, 1H), 7.28 – 7.23 (m, 2H), 7.08 – 7.00 (m, 1H), 3.11 (tt, *J* = 11.7, 3.9 Hz, 1H), 2.59 – 2.35 (m, 4H), 2.32 (s, 3H), 2.16 (ddq, *J* = 12.9, 6.6, 3.0 Hz, 1H), 2.05 – 1.98 (m, 1H), 1.90 – 1.69 (m, 2H). $^{13}C$ NMR (100 MHz, $CDCl_3$) δ 210.5, 169.4, 148.0, 135.8, 127.5, 126.8, 126.4, 122.7, 47.8, 41.1, 38.0, 31.4, 25.5, 20.9. LRMS (ESI) calcd for $C_{14}H_{17}O_3Na$ $[M + Na]^+$ m/z 255.11, found 255.22.

(±)-**S16: *trans*-2-(dispiro[adamantane-2,3'-[1,2,4]trioxolane-5',1''-cyclohexan]-3''-yl)phenol.**

To a 100 mL flask containing 40 mL hexane was added (±)-**S14** (850 mg, 3.7 mmol, 1.0 equiv.) and adamantan-2-one O-methyl oxime (1.64 g, 9.2 mmol, 2.5 equiv.) The mixture was cooled to -78 °C and ozone was bubbled through the solution. The $O_2$ flow rate was 1 liter/min, ozone gauge set to 3.5 (this setting amounts to ~ 6 g/hour ozone production). The reaction flask was wrapped with aluminum foil and the reaction mixture stirred at -78 °C for 4 hours, at which point the reaction was complete based on UPLC-MS. The reaction mixture was then bubbled with $N_2$ for 10 mins and concentrated to afford crude (±)-**S15** which was used without further purification in the next step.

To the crude (±)-**S15** in THF (7 mL) and MeOH (14 mL) was added 15% aqueous KOH solution (3 mL), and the mixture heated at 50 °C for 1.5 hour, after which the solution was cooled to room temperature and concentrated under reduced pressure. To the resulting residue was then added $H_2O$ (30 mL) and 1M HCl aq. until the solution reached pH ~ 4, and the solution extracted with EtOAc (100 mL). The organic layer was then separated and washed with brine, dried over anhydrous $Na_2SO_4$, filtered, and concentrated under reduced pressure. The crude residue was then purified by flash column chromatography (80 g silica gel cartridge, 0–50 % EtOAc/hexanes, product eluted around 30%) to yield (±)-**S16** (0.9 g, 70 %) as a colorless oil. $^1H$ NMR (400 MHz, $CDCl_3$) δ 7.15 (dd, *J* = 7.7, 1.7 Hz, 1H), 7.08 (td, *J* = 7.7, 1.7 Hz, 1H), 6.90 (td, *J* = 7.5, 1.2 Hz, 1H), 6.77 (dd, *J* = 7.9, 1.2 Hz, 1H), 3.15 (tt, *J* = 12.9, 3.4 Hz, 1H), 2.15 – 1.62 (m, 21H), 1.44 (qd, *J* =

12.4, 3.4 Hz, 1H), $^{13}C$ NMR (100 MHz, $CDCl_3$) δ 153.2, 131.5, 127.2, 127.1, 120.9, 115.7, 111.5, 109.5, 40.5, 39.4, 36.9, 36.5, 36.5, 35.2, 35.0, 34.9, 34.9, 34.5, 31.2, 27.0, 26.6, 23.7. LRMS (ESI) calcd for $C_{22}H_{28}O_4Na$ [M + Na]$^+$ m/z 379.20, found 379.19.

**(±)-5 (RLA-5351): *trans*-4-(2-(2-(dispiro[adamantane-2,3'-[1,2,4]trioxolane-5',1''-cyclohexan]-3''-yl)phenoxy)ethyl)morpholine.**
To a solution of *trans*-2-(dispiro[adamantane-2,3'-[1,2,4]trioxolane-5',1''-cyclohexan]-3''-yl)phenol ((±)-**S16**, 50.0 mg, 0.14 mmol, 1.0 equiv.) in dry $CH_3CN$ (4 mL) was added tetrabutylammonium hydrogen sulfate (14.3 mg, 0.04 mmol, 0.3 equiv.) and powdered NaOH (20.2 mg, 0.51 mmol, 3.6 equiv.). This mixture was then allowed to stir at room temperature for 30 minutes, at which point 4-(2-chloroethyl)morpholine hydrochloride (42.0 mg, 0.28 mmol, 2.0 equiv.) was added to the solution. The mixture was then placed in an oil bath that had been preheated to 55˚C, and was allowed to stir at this temperature for 14 hours. At this point the reaction was judged complete by TLC and LCMS analysis, and so the reaction mixture was allowed to cool to room temperature. The mixture was then diluted with EtOAc (20 mL) prior to the addition of $H_2O$ (10 mL), which served to dissolve all of the inorganic solids present. The organic phase was separated and the aqueous layer extracted with EtOAc (20 mL). The combined organic phases were then washed with brine (10 mL), dried over anhydrous $Na_2SO_4$, filtered, and concentrated under reduced pressure. The residue was then purified through by column chromatography (12 g silica gel cartridge, 0–10 % MeOH/$CH_2Cl_2$) to yield (±)-**5** (40.2 mg, 61%). $^1H$ NMR (400 MHz, $CDCl_3$) δ 7.20 – 7.09 (m, 2H), 7.00 – 6.87 (m, 1H), 6.82 (d, *J* = 8.3 Hz, 1H), 4.15 (t, *J* = 5.4 Hz, 2H), 3.87 – 3.70 (m, 4H), 3.21 (t, *J* = 13.3 Hz, 1H), 3.01 – 2.82 (m, 2H), 2.70 (s, 4H), 2.61 (s, 2H), 2.20 – 1.17 (m, 22H); $^{13}C$ NMR (100 MHz, $CDCl_3$) δ 155.9, 133.9, 127.2, 126.8, 121.0, 111.5, 111.3, 109.4, 66.8, 66.1, 57.9, 54.2, 41.1, 40.8, 36.9, 36.5, 35.0, 34.9, 34.5, 31.3, 27.0, 26.6, 23.9. LRMS (ESI) calcd for $C_{28}H_{40}NO_5$ [M + H]$^+$ m/z 470.28, found 479.42.

**(±)-S17: trans-2-(dispiro[adamantane-2,3'-[1,2,4]trioxolane-5',1''-cyclohexan]-3''-yl)phenyl (4-nitrophenyl) carbonate.**
To a 20 mL vial charged with (±)-**S16** (230 mg, 0.65 mmol, 1 equiv.) dissolved in $CH_2Cl_2$ (5 mL) was added 4-nitrophenyl chloroformate (325 mg, 1.6 mmol, 2.5 equiv.), DIEA (0.34 mL, 1.9 mmol, 3 equiv.), and DMAP (8.0 mg, 0.07 mmol, 0.1 equiv.) at 0 °C. The reaction mixture was stirred at room temperature overnight and then treated with 1M NaOH aq. and diluted with $CH_2Cl_2$. The organic phase was washed with aq. $NaHCO_3$ thrice, once with brine, and then dried over $MgSO_4$ and concentrated. The crude residue was then purified by flash column chromatography (40 g silica gel cartridge, 0–80 % EtOAc/Hexanes) to yield the product (260 mg, 77 %) as a yellow oil. $^1H$ NMR (400 MHz, $CDCl_3$) δ 8.33 (d, *J* = 8.9 Hz, 2H), 7.57 (d, *J* = 8.8 Hz, 2H), 7.43 – 7.15 (m, 4H), 3.11 (tt, *J* = 12.5, 3.4 Hz, 1H), 2.26 – 2.13 (m, 1H), 2.06 – 1.65 (m, 20H), 1.58 – 1.42 (m, 1H). $^{13}C$ NMR (100 MHz, $CDCl_3$) δ 155.6, 151.5, 148.2, 145.8, 137.1, 127.6, 127.4, 125.5, 122.2, 121.7, 111.7, 108.9, 41.4, 36.8, 36.5, 36.5, 35.0, 34.9, 34.9, 34.9, 34.9, 34.3, 31.5, 26.9, 26.5, 23.7. LRMS (ESI) calcd for $C_{29}H_{32}NO_8$ [M + H]$^+$ m/z 522.21, found 522.91.

**(±)-7: *trans*-2-(dispiro[adamantane-2,3'-[1,2,4]trioxolane-5',1''-cyclohexan]-3''-yl)phenyl 4-aminopiperidine-1-carboxylate**
Step 1: A 20 mL vial was charged with (±)-**S17** (15 mg, 0.03 mmol, 1.0 equiv.), *tert*-butyl 4-aminopiperidine-1-carboxylate (8.7 mg, 0.4 mmol, 1.5 equiv.) and 5 mL of $CH_2Cl_2$. Triethylamine (12.0 µL, 0.09 mmol, 3.0 equiv.) was then added to the reaction mixture slowly and the mixture stirred at room temperature overnight. After diluting the reaction mixture with $CH_2Cl_2$, the organic phase was washed with 1M $NaHCO_3$ aq. 3 times, and brine. The organic layer was then dried over $MgSO_4$, filtered and concentrated, and the residue purified by flash column chromatography (12 g silica gel cartridge, 0–20 % MeOH/$CH_2Cl_2$) to yield the title compound in Boc protected form (6.4 mg, 38%). $^1H$ NMR (400 MHz, $CDCl_3$) δ 7.25 – 7.15 (m, 3H), 7.09 (d, *J* = 7.9 Hz, 1H), 5.12 (d, *J* = 7.7 Hz, 1H), 4.06 (s, 2H), 3.73 (s, 1H), 3.04 – 2.80 (m, 3H), 2.16 (d, *J* = 12.8 Hz, 1H), 2.03 – 1.47 (m, 32H), 1.29 – 1.22 (m, 2H); $^{13}C$ NMR (100 MHz, $CDCl_3$) δ 154.9, 154.0, 148.5, 137.6, 127.1, 126.8, 126.1, 123.0, 111.6, 109.2, 79.8, 48.8, 41.5, 36.9, 36.6, 35.2, 35.1, 35.0, 34.8, 34.3, 32.3,

31.0, 28.6, 27.0, 26.6, 23.8. LRMS (ESI) calcd for $C_{33}H_{46}N_2O_7Na$ [M + Na]$^+$ m/z 605.33, found 605.38.
Step 2: To a solution of *trans*-2-(dispiro[adamantane-2,3'-[1,2,4]trioxolane-5',1''-cyclohexan]-3''-yl)phenyl 4-((*tert*-butoxycarbonyl)amino)piperidine-1-carboxylate (62.4 mg, 107 μmol, 1.0 equiv.) in methanol (1.0 mL) cooled to 0 ˚C in an ice bath, was added acetyl chloride (228 uL, 3.2 mmol, 30 equiv.) dropwise via microsyringe. The solution was allowed to stir for 10 minutes at 0 ˚C, after which the solution was removed from the bath and allowed to warm to room temperature overnight. The reaction mixture was then quenched with $NH_3$ in MeOH (7M). The resulting solution was concentrated and purified by column chromatography (12 g silica gel cartridge, 0–20% of 10% $NH_4OH$ in MeOH/$CH_2Cl_2$) to yield (±)-**7** (11.1 mg, 21.5 %). $^1$H NMR (400 MHz, MeOD) δ 7.40 – 7.30 (m, 1H), 7.29 – 7.18 (m, 2H), 7.09 – 7.00 (m, 1H), 4.46 (s, 1H), 4.24 (s, 1H), 3.31 – 3.17 (m, 2H), 3.14 – 2.87 (m, 2H), 2.30 – 1.40 (m, 28H), $^{13}$C NMR (100 MHz, $CD_3OD$) δ 155.3, 149.8, 138.7, 128.1, 127.4, 123.9, 112.5, 110.2, 43.4 (m), 37.9, 37.8, 37.7, 36.4, 35.9, 35.8, 35.7, 35.2, 32.0 (rotamer, m), 28.3, 28.0, 24.8. LRMS (ESI) calcd for $C_{28}H_{39}N_2O_5$ [M + H]$^+$ m/z 483.27, found 483.27.

(±)-**8: *trans*-2-(dispiro[adamantane-2,3'-[1,2,4]trioxolane-5',1''-cyclohexan]-3''-yl)phenyl (2-amino-2-methylpropyl)carbamate**
To a solution of (±)-**17** (35 mg, 0.07 mmol, 1.0 equiv.) in $CH_2Cl_2$ was added $Et_3N$ (28 μL, 0.20 mmol, 3.0 equiv.) followed by 1,1-dimethyl-1,2-ethanediamine (11 μL, 0.1 mmol, 1.5 equiv.). The bright yellow mixture was allowed to stir at room temperature for 16 h and then quenched by the addition of $NH_3$ in MeOH (7M). The resulting solution was concentrated and purified by column chromatography (12 g silica gel cartridge, 0–20% of 10% $NH_4OH$ in MeOH/$CH_2Cl_2$) to yield the product (±)-**8** (10.8 mg, 34 %). $^1$H NMR (400 MHz, $CDCl_3$) δ 7.26 – 7.13 (m, 3H), 7.13 – 7.03 (m, 1H), 5.69 (t, $J$ = 6.3 Hz, 1H), 3.23 (dd, $J$ = 13.5, 6.8 Hz, 1H), 3.12 (dd, $J$ = 13.5, 5.8 Hz, 1H), 3.02 (tt, $J$ = 12.6, 3.2 Hz, 1H), 2.17 (dq, $J$ = 13.0, 3.7, 2.8 Hz, 1H), 1.98 – 1.65 (m, 20H), 1.47 – 1.38 (m, 1H), 1.19 (s, 6H). $^{13}$C NMR (100 MHz, $CDCl_3$) δ 155.5, 148.6, 137.6, 127.1, 126.8, 126.0, 123.0, 111.6, 109.2, 52.8, 50.6, 41.4, 36.9, 36.6, 36.5, 35.3, 35.0, 35.0, 34.9, 34.4, 31.1, 28.4, 28.3, 28.2, 27.0, 26.6, 23.8. LRMS (ESI) calcd for $C_{27}H_{39}N_2O_5$ [M + H]+ m/z 471.28, found 471.20.

(±)-**S18: *trans*-3-((dispiro[adamantane-2,3'-[1,2,4]trioxolane-5',1''-cyclohexan]-3''-yl)phenyl (4-nitrophenyl) carbonate**
To an oven-dried round bottom flask containing a magnetic stir bar under an Ar(g) atmosphere was added (±)-**S12** (0.300 mg, 0.84 mmol, 1.0 equiv), dichloromethane (10 mL), N,N-diisopropylethylamine (0.48 mL, 2.74 mmol, 3.25 equiv), and 4-dimethylaminopyridine (0.123 g, 1.01 mmol, 1.2 equiv). The mixture was cooled to 0 °C while 4-nitrophenyl chloroformate (0.551 g, 2.74 mmol, 3.25 equiv) was added as a solid in one portion. The solution was allowed to stir at room temperature for 3 hours. The reaction was then diluted with DI $H_2O$ (100 mL) and subsequently extracted with EtOAc (100 mL). The organic layer was washed repeatedly by potassium carbonate solution until the aqueous layer was colorless and no longer yellow (indicating successful removal of *p*-nitrophenol from the organic layer). The organic layer was dried over anhydrous $Na_2SO_4$, filtered, and concentrated under reduced pressure to yield a thick yellow oil. The residue was then purified through flash column chromatography (80 g silica gel cartridge, 0–25% EtOAc/Hexanes, product eluted during 8% EtOAc/Hex) to yield intermediate (±)-**S18** (347 mg, 79%) as a colorless solid. $^1$H NMR (400 MHz, $CDCl_3$) δ 8.31 - 8.36 (m, 2H), 7.49 - 7.53 (m, 2H), 7.35 - 7.41 (m, 1H), 7.12 - 7.19 (m, 3H), 3.01 (tt, $J$ = 3.32, 12.63 Hz, 1H), 2.87 (tt, $J$ = 3.32, 12.63 Hz, 1H), 2.15 - 2.21 (m, 1H), 1.77 - 2.05 (m, 13H), 1.65 - 1.75 (m, 5H), 1.59 - 1.65 (m, 1H), 1.39 - 1.46 (m, 1H); $^{13}$C NMR (100 MHz, CDCl3) δ 155.4, 151.2, 150.9, 148.2, 148.0, 145.7, 129.8, 125.5, 125.4, 121.9, 119.2, 118.6, 112.0 (minor diastereomer), 111.6, 108.9, 42.0, 41.7, 41.5, 36.9, 36.5, 35.0, 34.9, 34.2, 33.5, 32.7, 27.0, 26.6, 23.5. MS (ESI) calculated for $C_{29}H_{31}NNaO_8$ [M + Na]$^+$ *m/z* 544.19, found 544.02.

(±)-**9: *trans*-3-(dispiro[adamantane-2,3'-[1,2,4]trioxolane-5',1''-cyclohexan]-3''-yl)phenyl 4-aminopiperidine-1-carboxylate**
Step 1: To a solution of (±)-**S18** (87.4 mg, 0.15 mmol, 1.0 equiv) in dichloromethane (1.5 mL), was added 4-(N-Boc-amino)piperidine (48 mg, 0.24 mmol, 1.5 equiv), and triethylamine (34 μL, 0.24 mmol, 1.5 equiv) at room temperature and was set to stir for 2 days. The reaction was deemed

complete by LCMS and TLC, diluted with EtOAc (10 mL) and subsequently extracted with 1M NaOH (15 mL). The organic layer was washed with brine (12 mL), dried over anhydrous $MgSO_4$, filtered, and concentrated under reduced pressure to yield a clear oil. The residue was then purified through silica gel flash column chromatography (0-30% EtOAc/Hex), with the product eluting around 20% EtOAc/Hex to yield *trans*-3-(dispiro[adamantane-2,3'-[1,2,4]trioxolane-5',1''-cyclohexan]-3''-yl)phenyl 4-((tert-butoxycarbonyl)amino)piperidine-1-carboxylate (40 mg, 45%) as a colorless oil. $^1H$ NMR (400 MHz, $CDCl_3$) δ 7.33 – 7.23 (m, 1H), 7.06 (d, *J* = 7.7 Hz, 1H), 6.97 – 6.91 (m, 2H), 4.53 (s, 1H), 4.22 (s, 1H), 4.14 (q, *J* = 7.1 Hz, 4H), 3.07 (d, *J* = 49.6 Hz, 2H), 2.81 (tt, *J* = 12.8, 3.5 Hz, 1H), 2.05 – 1.57 (m, 24H), 1.48 (s, 9H). MS (ESI) calculated for $C_{33}H_{46}N_2NaO_7$ $[M + Na]^+$ *m/z* 605.73, found 605.33.
Step 2: To a solution of *trans*-3-(dispiro[adamantane-2,3'-[1,2,4]trioxolane-5',1''-cyclohexan]-3''-yl)phenyl 4-((tert-butoxycarbonyl)amino)piperidine-1-carboxylate (29.1 mg, 0.05 mmol, 1.0 equiv) in methanol (1.5 mL), was added acetyl chloride (110 uL, 1.5 mmol, 30 equiv) in methanol (0.7 mL). The reaction was then allowed to stir at 0° C, warmed up to room temperature and stirred for 3 days, at which point the reaction was judged complete by LCMS. The reaction mixture was concentrated to dryness and triturated with diethyl ether to yield the desired product (21.7 mg, 89.4%) as a hydrochloride salt, a white powder. $^1H$ NMR (400 MHz, DMSO-$d_6$) δ 7.33 – 7.26 (m, 1H), 7.11 (d, *J* = 7.6 Hz, 1H), 7.07 – 6.88 (m, 2H), 4.29 – 3.87 (m, 2H), 3.27-3.03 (m,1H), 3.03 – 2.90 (m, 1H), 2.69 (m, *J* = 15.8, 12.2, 7.4, 4.6 Hz, 1H), 2.17-1.15 (m, 24H). MS (ESI) calculated for $C_{28}H_{38}N_2NaO_5$ $[M + Na]^+$ *m/z* 505.61, found 505.15.

**(±)-10: *trans*-3-((dispiro[adamantane-2,3'-[1,2,4]trioxolane-5',1''-cyclohexan]-3''-yl)phenyl (2-amino-2-methylpropyl)carbamate**

To a solution of (±)-**S18** (87.4 mg, 0.15 mmol, 1.0 equiv) in dichloromethane (1.5 mL), was added 2-methylpropane-1,2-diamine (20 mg, 0.24 mmol, 1.5 equiv), and triethylamine (34 μL, 0.24 mmol, 1.5 equiv) at room temperature and was set to stir for 2 days. The reaction was deemed complete by LCMS and TLC, diluted with EtOAc (10 mL) and subsequently extracted with 1M NaOH (15 mL). The organic layer was washed with brine (12 mL), dried over anhydrous $MgSO_4$, filtered, and concentrated under reduced pressure to yield a clear oil. The residue was then purified through silica gel flash column chromatography (0-30% EtOAc/Hex), the product eluting around 20% EtOAc/Hex to yield (±)-**10** (19.1 mg, 23%) as a white solid. $^1H$ NMR (400 MHz, DMSO-$d_6$) δ 7.29 (t, *J* = 7.9 Hz, 1H), 7.13 – 7.01 (m, 2H), 6.95 (dd, *J* = 8.1, 2.3 Hz, 1H), 3.08 (t, *J* = 5.1 Hz, 2H), 2.78 – 2.63 (m, 1H), 1.94 – 1.63 (m, 22H), 1.12 (t, *J* = 2.2 Hz, 6H). MS (ESI) calculated for $C_{27}H_{39}N_2O_5$ $[M + H]^+$ *m/z* 471.61, found 471.30.

**(±)-11: *trans*-3-(dispiro[adamantane-2,3'-[1,2,4]trioxolane-5',1''-cyclohexan]-3''-yl)phenyl 3-aminopyrrolidine-1-carboxylate**

Step1: To a solution (±)-**S18** in dichloromethane (1.5 mL), was added 3-(Boc-amino)pyrrolidine (44.7 mg, 0.24 mmol, 1.5 equiv), and triethylamine (34 μL, 0.24 mmol, 1.5 equiv) at room temperature and was set to stir for 2 days. The reaction was deemed complete by LCMS and TLC, diluted with EtOAc (10 mL) and subsequently extracted with 1M NaOH (15 mL). The organic layer was washed with brine (12 mL), dried over anhydrous $MgSO_4$, filtered, and concentrated under reduced pressure to yield a clear oil. The residue was then purified through silica gel flash column chromatography (0-30% EtOAc/Hex), the product eluting around 20% EtOAc/Hex to yield Boc-protected **11** (55 mg, 65%) as a colorless oil. $^1H$ NMR (400 MHz, $CDCl_3$) δ 7.29 (d, *J* = 2.7 Hz, 1H), 7.06 (d, *J* = 7.7 Hz, 1H), 7.04 – 6.92 (m, 2H), 4.71 (s, 1H), 4.30 (s, 1H), 3.93 – 3.70 (m, 1H), 3.67 – 3.55 (m, 2H), 3.40 (ddd, *J* = 27.0, 11.5, 4.5 Hz, 1H), 2.82 (tt, *J* = 12.9, 3.5 Hz, 1H), 2.04 – 1.54 (m, 24H), 1.49 (s, 9H). $^{13}C$ NMR (100 MHz, $CDCl_3$) δ 153.27, 151.43, 147.14, 129.17, 123.74, 120.07, 119.58, 111.37, 108.98, 54.62, 44.74, 41.94, 41.65, 41.23, 39.28, 36.81, 36.66, 36.44, 35.77, 34.83, 34.19, 32.64, 30.96, 29.70, 26.89, 26.50, 25.85, 24.70, 23.50. MS (ESI) calculated for $C_{32}H_{44}N_2NaO_7$ $[M + Na]^+$ *m/z* 591.70, found 591.48.
Step 2: To a solution of the intermediate above (28.4 mg, 0.05 mmol, 1.0 equiv) in methanol (1.5 mL), was added acetyl chloride (110 uL, 1.5 mmol, 30 equiv) in methanol (0.7 mL). The reaction was then allowed to stir at 0° C, warmed up to room temperature and stirred for 3 days, at which point, the reaction was judged complete by LCMS. The reaction mixture was concentrated to dryness and triturated with diethyl ether to yield (±)-**11** (16.8 mg, 72%) as a hydrochloride salt, a

white powder. $^{1}H$ NMR (400 MHz, $CDCl_3$) δ 7.28 (s, 1H), 7.05 (dd, *J* = 7.6, 1.5 Hz, 1H), 7.00 – 6.96 (m, 2H), 4.72 (s, 1H), 4.30 (s, 1H), 3.86-3.74 (m, *J* = 11.6, 6.2 Hz, 1H), 3.67 (td, *J* = 6.8, 6.2, 1.8 Hz, 1H), 3.58 (td, *J* = 7.1, 2.6 Hz, 1H), 3.43-3.36 (m, *J* = 11.5, 4.4 Hz, 1H), 2.81 (tt, *J* = 12.9, 3.5 Hz, 1H), 2.05 –1.27 (m, 24H). $^{13}C$ NMR (100 MHz, $CDCl_3$) δ 155.24, 153.11, 151.32, 147.18, 129.20, 123.84, 119.99, 111.82, 108.99, 77.36, 52.24, 44.44, 41.93, 41.64, 36.80, 36.41, 34.82, 34.19, 34.07, 32.64, 32.09, 31.13, 30.94, 28.38, 26.88, 26.49, 23.49. MS (ESI) calculated for $C_{27}H_{36}N_2NaO_5$ $[M + Na]^+$ *m/z* 491.58, found 491.20.

**(±)-12: *trans*-3-(dispiro[adamantane-2,3'-[1,2,4]trioxolane-5',1''-cyclohexan]-3''-yl)phenyl (2-aminoethyl)carbamate**

To a solution (±)-**S18** (60 mg, 0.12 mmol, 1.0 equiv) in dichloromethane (2.0 mL) was added $Et_3N$ (24 μL, 0.17 mmol, 1.5 equiv), followed by ethylenediamine (12 μL, 0.17 mmol, 1.5 equiv) at room temperature. The bright yellow mixture was then allowed to stir at room temperature for 5 h, at which point, the reaction was judged complete by TLC and LCMS. The reaction was then diluted with DI $H_2O$ (100 mL) and subsequently extracted with EtOAc (100 mL). The organic layer was washed repeatedly by potassium carbonate solution until the aqueous layer was colorless and no longer yellow (meaning that all of the *p*-nitrophenol had been successfully removed from the organic layer). The combined aqueous layers were then back extracted with EtOAc (30 mL). The combined organic layers were then washed with brine (20 mL), dried over anhydrous $Na_2SO_4$, filtered, and concentrated under reduced pressure. The residue was then purified through flash column chromatography (12 g silica gel cartridge, 0–100% EtOAc/Hexanes, followed by 0–20% MeOH (containing 0.7 N $NH_3$)/$CH_2Cl_2$, and desired product eluted during 20% MeOH (containing 0.7 N $NH_3$)/ $CH_2Cl_2$], to yield (±)-**12** (38.0 mg, 75%) as a colorless oil. 1H NMR (400 MHz, $CDCl_3$) δ 7.22 - 7.27 (m, 1H), 7.03 (br d, *J* = 7.55 Hz, 1H), 6.91 - 7.01 (m, 2H), 3.51 (br s, 2H), 3.13 (br s, 2H), 2.06 - 2.13 (m, 1H), 1.88 - 2.01 (m, 6H), 1.83 - 1.88 (m, 1H), 1.74 - 1.83 (m, 5H), 1.58 - 1.73 (m, 8H), 1.27 - 1.46 (m, 2H), 1.15 - 1.27 (m, 1H); MS (ESI) calculated for $C_{25}H_{35}N_2O_5$ $[M + H]^+$ *m/z* 443.25, found 443.28.

**(±)-13: *trans*-3-(dispiro[adamantane-2,3'-[1,2,4]trioxolane-5',1''-cyclohexan]-3''-yl)phenyl 3-aminoazetidine-1-carboxylate**

Step 1: To a solution of (±)-**S18** (87.4 mg, 0.15 mmol, 1.0 equiv) in dichloromethane (1.5 mL), was added *tert*-butyl azetidin-3-ylcarbamate (41.2 mg, 0.24 mmol, 1.5 equiv), and triethylamine (34 μL, 0.24 mmol, 1.5 equiv) at room temperature and was set to stir for 2 days. The reaction was deemed complete by LCMS and TLC, diluted with EtOAc (10 mL) and subsequently extracted with 1M NaOH (15 mL). The organic layer was washed with brine (12 mL), dried over anhydrous $MgSO_4$, filtered, and concentrated under reduced pressure to yield a clear oil. The residue was then purified through silica gel flash column chromatography (0-30% EtOAc/Hex), the product eluting around 20% EtOAc/Hex to yield Boc-protected **13** (30.7mg, 37%) as a colorless oil. $^{1}H$ NMR (400 MHz, Chloroform-*d*) δ 7.28 (dt, *J* = 7.5, 4.6 Hz, 1H), 7.05 (dt, *J* = 7.7, 1.4 Hz, 1H), 6.97 (ddd, *J* = 5.3, 3.4, 1.4 Hz, 2H), 4.44 (t, *J* = 45.6 Hz, 2H), 3.97 (s, 1H), 3.75 – 3.60 (m, 1H), 3.09 – 2.92 (m, 1H), 2.81 (tt, *J* = 12.9, 3.5 Hz, 1H), 2.61 (d, *J* = 6.1 Hz, 1H), 2.26 – 1.54 (m, 22H), 1.52 – 1.37 (m, 9H). $^{13}C$ NMR (100 MHz, $CDCl_3$) δ 154.88, 154.25, 151.15, 151.04, 147.24, 129.24, 123.93, 119.85, 119.37, 111.83, 111.39, 108.95, 62.86, 44.45, 41.92, 41.62, 41.36, 40.82, 36.79, 36.43, 34.82, 34.18, 32.62, 29.71, 28.34, 26.88, 26.49, 23.48. MS (ESI) calculated for $C_{31}H_{42}N_2NaO_7$ $[M + Na]^+$ *m/z* 577.67, found 577.32.

Step 2: To a solution of Boc-protected **13** (27.7 mg, 0.05 mmol, 1.0 equiv) in methanol (1.5 mL), was added acetyl chloride (110 uL, 1.5 mmol, 30 equiv) in methanol (0.7 mL). The reaction was then allowed to stir at $0^{o}$ C, warmed up to room temperature and stirred for 3 days, at which point, the reaction was judged complete by LCMS. The reaction mixture was concentrated to dryness and triturated with diethyl ether to yield (±)-**13** (12.7 mg, 56%) as a hydrochloride salt, a white powder. $^{1}H$ NMR (400 MHz, DMSO-$d_6$) δ 7.30 (qd, *J* = 7.6, 3.4 Hz, 1H), 7.13 (m, *J* = 13.5, 9.3, 4.0 Hz, 1H), 7.02 – 6.90 (m, 2H), 4.41-3.98 (m, 4H), 3.10 (d, *J* = 2.3 Hz, 1H), 2.68 (qd, *J* = 11.9, 5.7 Hz, 1H), 2.17 – 1.11 (m, 22H). MS (ESI) calculated for $C_{26}H_{34}N_2NaO_5$ $[M + Na]^+$ *m/z* 477.56, found 477.25.

(±)-**14: *trans*-3-(dispiro[adamantane-2,3'-[1,2,4]trioxolane-5',1''-cyclohexan]-3''-yl)phenyl ((1-aminocyclobutyl)methyl)carbamate**

Step 1: To a solution of (±)-**S18** (101.8 mg, 0.20 mmol) in dichloromethane (2 mL), was added *tert*-butyl(1-(aminomethyl)cyclobutyl)carbamate (55 mg, 0.27 mmol), and triethylamine (38.3 μL, 0.27 mmol). After stirring at room temperature for 72 h, the reaction mixture was then diluted with ethyl acetate and washed with thrice with 1M aqueous sodium hydroxide solution. The organic layer was then washed with brine, dried over magnesium sulfate, filtered, and concentrated under reduced pressure. The residue was then purified by flash column chromatography (0-30% ethyl acetate/hexanes) to yield 35.8 mg (32%) of Boc-protected **14** as a clear oil. $^1$H NMR ($CDCl_3$, 400 MHz) δ 7.21 – 7.33 (m, 1H), 6.91-7.10 (m, 3H), 5.82-6.21 (m, 2H), 3.54-3.69 (m, 2H), 2.73-2.78 (m, 1H), 1.25-2.20 (m, 37H).

Step 2: To a cooled (0° C) solution of Boc-protected **14** (35.8 mg, 0.06 mmol) in methanol (1.5 mL), was added acetyl chloride (132 uL, 1.8 mmol). The reaction was then allowed to stir at room temperature for 72 h. The reaction mixture was concentrated azeotropically with toluene under reduced pressure and the residue was washed with diethyl ether to obtain (±)-**14** 22.5 mg (70%) as a white powdery hydrochloride salt. $^1$H NMR ($CD_3OD$, 400 MHz) Shift 7.33 (t, 1H, J=7.9 Hz), 7.13 (d, 1H, J=7.8 Hz), 6.99-7.06 (m, 2H), 3.58 (s, 2H), 2.76-2.82 (m, 1H), 2.26-2.30 (m, 3H), 1.44-2.06 (m, 25H). MS (ESI) calculated for $C_{28}H_{39}N_2O_5$ $[M + H]^+$ *m/z* 483.28, found 483.37.

(±)-**15: *trans*-3-(dispiro[adamantane-2,3'-[1,2,4]trioxolane-5',1''-cyclohexan]-3''-yl)phenyl morpholine-1-carboxylate**

To a solution of (±)-**S18** (60 mg, 0.12 mmol, 1.0 equiv) in dichloromethane (2.0 mL) was added $Et_3N$ (24 μL, 0.17 mmol, 1.5 equiv), followed by morpholine hydrochloride (23 mg, 0.19 mmol, 1.5 equiv) at room temperature. The bright yellow mixture was then allowed to stir at room temperature for 5 h, at which point, the reaction was judged complete by TLC and LCMS. The reaction was then diluted with $H_2O$ (100 mL) and subsequently extracted with EtOAc (100 mL). The organic layer was washed repeatedly by potassium carbonate solution until the aqueous layer was colorless and no longer yellow (indicating successful removal of *p*-nitrophenol from the organic layer). The combined aqueous layers were then back extracted with EtOAc (30 mL). The combined organic layers were then washed with brine (20 mL), dried over anhydrous $Na_2SO_4$, filtered, and concentrated under reduced pressure. The residue was then purified through flash column chromatography (12 g silica gel cartridge, 0–100% EtOAc/Hexanes, followed by 0–20% MeOH (containing 0.7 N $NH_3$)/ $CH_2Cl_2$, and desired product (±)-**15** eluted near 10% MeOH (containing 0.7 N $NH_3$)/ $CH_2Cl_2$, to yield the desired product (31.6 mg, 56.1%) as a colorless oil $^1$H NMR (400 MHz, $CDCl_3$) δ 7.28 - 7.33 (m, 1H), 7.06 (d, J = 7.79 Hz, 1H), 6.94 – 6.98 (m, 2H), 3.75 - 3.8 (t, 4H), 3.55 - 3.73 (d, 4H), 2.80 (tt, J = 3.17, 12.66 Hz, 1H), 2.13 (br d, J = 13.39 Hz, 1H), 2.01 - 2.10 (m, 1H), 1.87 - 2.01 (m, 6H), 1.77 - 1.85 (m, 4H), 1.63 - 1.73 (m, 7H), 1.43 - 1.63 (m, 1H), 1.31 - 1.42 (m, 1H), 1.19 - 1.31 (m, 1H); $^{13}$C NMR (100 MHz, $CDCl_3$) δ 153.79, 151.34, 147.30, 129.28, 124.01, 120.02, 119.51, 111.40, 108.94, 66.61, 44.88, 41.94, 41.66, 41.36, 36.80, 36.43, 36.41, 34.81, 34.78, 34.19, 32.65, 26.88, 26.49, 23.48; MS (ESI) calculated for $C_{27}H_{35}NO_6$ $[M + Na]^+$ *m/z* 492.24, found 492.1.

(±)-**16: *trans*-3-(dispiro[adamantane-2,3'-[1,2,4]trioxolane-5',1''-cyclohexan]-3''-yl)phenyl ((1-aminocyclopentyl)methyl)carbamate**

Step 1: To a solution of (±)-**S18** (80.4 mg, 0.15 mmol, 1.0 equiv) in dichloromethane (1.5 mL), was added *tert*-butyl (1-(aminomethyl)cyclopentyl)carbamate (52.1 mg, 0.24 mmol, 1.5 equiv), and triethylamine (34 μL, 0.24 mmol, 1.5 equiv) at room temperature and was set to stir for 2 days. The reaction was deemed complete by LCMS and TLC, diluted with EtOAc (10 mL) and subsequently extracted with 1M NaOH (15 mL). The organic layer was washed with brine (12 mL), dried over anhydrous $MgSO_4$, filtered, and concentrated under reduced pressure to yield a clear oil. The residue was then purified through silica gel flash column chromatography (0-30% EtOAc/Hex), the product eluting around 20% EtOAc/Hex to yield Boc-protected **16** (30.6 mg, 33%). $^1$H NMR (400 MHz, $CDCl_3$) δ 7.28 (s, 1H), 7.12 (t, 1H), 6.91 (d, 1H), 6.81 (d, 1H), 6.45 (s, 1H), 5.25 (s, 1H), 3.43 (s, 1H), 3.07 (s, 1H), 2.65 (s, 1H), 2.07 – 1.08 (m, 39H).

Step 2: To a solution of Boc-protected **16** (30.6 mg, 0.05 mmol, 1.0 equiv) in methanol (1.5 mL), was added acetyl chloride (110 uL, 1.5 mmol, 30 equiv) in methanol (0.7 mL). The reaction was then allowed to stir at 0° C, warmed up to room temperature and stirred for 3 days, at which point, the reaction was judged complete by LCMS. The reaction mixture was concentrated to dryness and triturated with diethyl ether to yield (±)-**16** (24.5 mg, 89.4%) as a hydrochloride salt, a white powder. $^1$H NMR (400 MHz, DMSO) δ 8.16 – 8.00 (m, 3H), 7.29 (t, 1H), 7.16 – 6.96 (m, 3H), 3.32 – 3.27 (m, 2H), 2.77 – 2.65 (m, 1H), 1.76 (s, 31H). MS (ESI) calculated for $C_{29}H_{40}N_2O_5$ [M + H]$^+$ *m/z* 496.65, found 497.38.

(±)-**17: *trans*-4-(dispiro[adamantane-2,3'-[1,2,4]trioxolane-5',1''-cyclohexan]-3''-yl)phenyl (2-amino-2-methylpropyl)carbamate**

To a solution of (±)-**S4** (52.0 mg, 0.10 mmol, 1.0 equiv) in dichloromethane (2.0 mL) was added $Et_3N$ (21 μL, 0.15 mmol, 1.5 equiv), followed by 2-methylpropane-1,2-diamine (16 μL, 0.15 mmol, 1.5 equiv) at room temperature. The bright yellow mixture was then allowed to stir at room temperature for 5 h, at which point, the reaction was judged complete by TLC and LCMS. The reaction was then diluted with $H_2O$ (100 mL) and subsequently extracted with EtOAc (100 mL). The organic layer was washed repeatedly with potassium carbonate solution until the aqueous layer was colorless and no longer yellow (indicating successful removal of *p*-nitrophenol from the organic layer). The combined aqueous layers were then back extracted with EtOAc (30 mL). The combined organic layers were then washed with brine (20 mL), dried over anhydrous $Na_2SO_4$, filtered, and concentrated under reduced pressure. The residue was then purified through flash column chromatography (12 g silica gel cartridge, 0–100% EtOAc/Hexanes, followed by 0–20% MeOH (containing 0.7 N $NH_3$)/$CH_2Cl_2$), to yield (±)-**17** (43.0 mg, 76%) as a white solid. $^1$H NMR (400 MHz, $CDCl_3$) δ 7.14 - 7.19 (m, 2H), 7.09 - 7.14 (m, 2H), 3.34 (br s, 2H), 2.78 (br t, *J* = 12.66 Hz, 1H), 2.07 - 2.13 (m, 1H), 1.93 - 2.04 (m, 7H), 1.65 - 1.86 (m, 14H), 1.59 - 1.64 (m, 1H), 1.37 - 1.46 (m, 1H), 1.39 (s, 4H), 1.27 - 1.33 (m, 2H); $^{13}$C NMR (100 MHz, $CDCl_3$) δ 156.2, 149.4, 143.0, 127.9, 122.1, 115.4, 111.5, 109.1, 56.1, 42.5, 42.3, 41.4, 41.1, 36.5, 34.9, 34.3, 34.3, 33.2, 34.9, 27.0, 26.6, 23.7, 23.6. MS (ESI) calculated for $C_{29}H_{41}N_2O_5$ [M + H]$^+$ *m/z* 471.29, found 471.27.

(±)-**18: *trans*-4-(dispiro[adamantane-2,3'-[1,2,4]trioxolane-5',1''-cyclohexan]-3''-yl)phenyl (4-trans-aminocyclohexyl)carbamate**

To a solution of (±)-**S4** (58 mg, 0.11 mmol, 1.0 equiv) in dichloromethane (2.0 mL) was added $Et_3N$ (24 μL, 0.17 mmol, 1.5 equiv), followed by (1r,4r)-cyclohexane-1,4-diamine (19 mg, 0.17 mmol, 1.5 equiv) at room temperature. The bright yellow mixture was then allowed to stir at room temperature for 5 h, at which point, the reaction was judged complete by TLC and LCMS. The reaction was then diluted with $H_2O$ (100 mL) and subsequently extracted with EtOAc (100 mL). The organic layer was washed repeatedly by potassium carbonate solution until the aqueous layer was colorless and no longer yellow (meaning that all of the *p*-nitrophenol had been successfully removed from the organic layer). The combined aqueous layers were then back extracted with EtOAc (30 mL). The combined organic layers were then washed with brine (20 mL), dried over anhydrous $Na_2SO_4$, filtered, and concentrated under reduced pressure. The residue was then purified through flash column chromatography (12 g silica gel cartridge, 0–100% EtOAc/Hexanes, followed by 0–20% MeOH (containing 0.7 N $NH_3$)/$CH_2Cl_2$) to afford the product (±)-**18** (43.0 mg, 76%) as a white solid. $^1$H NMR (400 MHz, $CDCl_3$) δ 7.13 (br d, *J* = 8.28 Hz, 2H), 6.98 (br d, *J* = 8.28 Hz, 2H), 3.44 (br s, 1H), 2.96 (br s, 2H), 2.74 (br t, *J* = 12.66 Hz, 1H), 2.55 - 2.66 (m, 1H), 1.99 - 2.09 (m, 3H), 1.83 - 1.98 (m, 9H), 1.61 - 1.82 (m, 13H), 1.12 - 1.37 (m, 6H); $^{13}$C NMR (100 MHz, $CDCl_3$) δ 149.3, 142.6, 127.6, 127.5, 121.5, 115.3, 111.4, 111.3, 109.0, 49.7, 49.5, 42.1, 41.3, 36.7, 36.4, 34.8, 34.7, 34.6, 34.1, 32.8, 31.7, 26.8, 26.5, 23.5. MS (ESI) calculated for $C_{29}H_{41}N_2O_5$ [M + H]$^+$ *m/z* 497.30, found 497.28.

(±)-**19: *trans*-4-(dispiro[adamantane-2,3'-[1,2,4]trioxolane-5',1''-cyclohexan]-3''-yl)phenyl (4-aminocyclohexyl)carbamate**

To a solution of (±)-**S4** (52.0 mg, 0.10 mmol, 1.0 equiv) in dichloromethane (2.0 mL) was added $Et_3N$ (21 μL, 0.15 mmol, 1.5 equiv), followed by ethylenediamine (12 μL, 0.18 mmol, 1.5 equiv) at room temperature. The bright yellow mixture was then allowed to stir at room temperature for 5 h, at which point, the reaction was judged complete by TLC and LCMS. The reaction was then diluted with $H_2O$ (100 mL) and subsequently extracted with EtOAc (100 mL). The organic layer was washed repeatedly by potassium carbonate solution until the aqueous layer was colorless and no longer yellow (indicating removal of *p*-nitrophenol from the organic layer). The combined aqueous layers were then back extracted with EtOAc (30 mL). The combined organic layers were then washed with brine (20 mL), dried over anhydrous $Na_2SO_4$, filtered, and concentrated under reduced pressure. The residue was then purified by flash column chromatography (12 g silica gel cartridge, 0–100% EtOAc/Hexanes, followed by 0–20% MeOH (containing 0.7 N $NH_3$)/$CH_2Cl_2$) to yield the desired product (±)-**S19** (43.0 mg, 76%) as a white foam. 1H NMR (400 MHz, $CDCl_3$) δ 7.03 - 7.06 (m, 2H), 6.77 - 6.80 (m, 1H), 3.37 (br s, 1H), 2.93 - 3.01 (m, 1H), 2.66 - 2.83 (m, 1H), 2.08 - 2.15 (m, 1H), 1.90 - 2.03 (m, 8H), 1.76 - 1.87 (m, 6H), 1.56 - 1.75 (m, 8H), 1.30 - 1.39 (m, 1H); 13C NMR (100 MHz, $CDCl_3$) δ 154.9, 137.3, 127.8, 121.6, 115.5, 111.4, 109.3, 42.5, 41.1, 41.0, 36.9, 36.5, 34.9, 34.8, 34.3, 34.2, 33.1, 33.1, 27.0, 26.6, 23.6. MS (ESI) calculated for $C_{29}H_{41}N_2O_5$ $[M + H]^+$ *m/z* 443.25, found 443.20.

(±)-**20: *trans*-4-(dispiro[adamantane-2,3'-[1,2,4]trioxolane-5',1''-cyclohexan]-3''-yl)phenyl 3-aminoazetidine-1-carboxylate**

Step 1: To a cooled (0° C) solution of (±)-**S4** (50 mg, 0.096 mmol) in *N,N'*-dimethylformamide (1 mL) were added 3-*N*-Boc-amino azetidine (25 mg, 0.14 mmol) and triethylamine (0.02 mL, 0.14 mmol). After stirring at 35° C for 72 h, the reaction mixture was diluted with ethyl acetate and washed multiple times with 1N aqueous sodium hydroxide solution. The organic layer was washed with brine, dried over $MgSO_4$, concentrated under reduced pressure, and purified by flash column chromatography (5-10% MeOH/dichloromethane) to afford 11 mg (21% yield) of Boc-protected **20** as yellow solid. $^1$H NMR ($CDCl_3$, 400 MHz) δ 7.19 (d, 2H, *J*=8.5 Hz), 7.04 (d, 2H, *J*=8.5 Hz), 4.99 (br s, 1H), 4.55 (br s, 1H), 4.44 (br s, 2H), 3.97 (br dd, 2H, *J*=1.5, 3.4 Hz), 2.77-2.84 (m, 1H), 1.97-2.15 (m, 8H), 1.65-1.86 (m, 13H), 1.49 (s, 9H), 1.30-1.36 (m, 1H).

Step 2: To a cooled (0° C) solution of Boc-protected **20** (9.4 mg, 0.017 mmol) in methanol (0.5 mL), was added acetyl chloride (12 uL, 0.17 mmol). The reaction was then allowed to stir at room temperature for 48 h. The reaction mixture was concentrated azeotropically with toluene *in vacuo* and the residue was washed with diethyl ether to obtain (±)-**20** (4.3 mg, 56%) as a white powdery hydrochloride salt. $^1$H NMR (DMSO-d6, 400 MHz) δ 8.55 (br s, 3H), 7.27-7.29 (m, 2H), 7.03-7.05 (m, 2H), 4.3-4.5 (m, 1H), 4.39 (br s, 1H), 4.18-4.20 (m, 2H), 4.03-4.06 (m, 2H), 2.68-2.71 (m, 1H), 1.65-1.93 (m, 18H), 1.20-1.60 (m, 4H). MS (ESI) calculated for $C_{26}H_{34}N_2O_5$ $[M + H]^+$ *m/z* 455.25, found 455.25.

(±)-**21: *trans*-4-(dispiro[adamantane-2,3'-[1,2,4]trioxolane-5',1''-cyclohexan]-3''-yl)phenyl ((*trans*)-3-aminocyclobutyl)carbamate**

Step 1: To a cooled (0° C) solution of (±)-**S4** (20 mg, 0.038 mmol) in N,N'-dimethylformamide (0.5 mL), were added *tert*-butyl 3-aminocyclobutyl)carbamate (11 mg, 0.058 mmol) and triethylamine (8 μL, 0.058 mmol). After stirring at room temperature overnight, the reaction mixture was then diluted with ethyl acetate and washed with thrice with 1M aqueous sodium hydroxide solution. The organic layer was then washed with brine, dried over magnesium sulfate, filtered, and concentrated under reduced pressure. The residue was then purified by flash column chromatography (0-100% ethyl acetate/hexanes) to obtain 12 mg (55%) of Boc-protected **21** as a colorless oil. $^1$H NMR ($CDCl_3$, 400 MHz) δ 7.20 (d, 2H, *J* = 8.5 Hz), 7.04-7.08 (m, 2H), 5.27 (br d, 1H, *J* = 6.6 Hz), 4.79 (br s, 1H), 4.24-4.32 (m, 2H), 2.80-2.84 (m, 1H), 2.36-2.40 (m, 4H), 2.13 (br d, 1H, *J* = 13.4 Hz), 1.63-2.02 (m, 20H), 1.47 (s, 9H), 1.35-1.39 (m, 1H).

Step 2: To a solution of Boc-protected **21** (49.5 mg, 0.09 mmol) in methanol (3.0 mL), was added acetyl chloride (186 uL, 2.61 mmol). The reaction was then allowed to stir at room temperature for 24 h. The reaction mixture was concentrated azeotropically with toluene under reduced pressure and the residue was washed with diethyl ether to obtain the desired product (±)-**21** (32 mg, 79%) as a white powdery salt. $^1$H NMR ($CDCl_3$, 400 MHz) δ 7.10 (br d, 2H, *J* = 8.0 Hz), 6.96-6.98 (m, 2H), 6.61 (br s, 1H), 4.48 (br s, 1H), 4.01 (br s, 1H), 3.75 (br s, 1H), 2.56-2.70 (m, 5H), 1.27-2.07 (m, 21H). MS (ESI) calculated for $C_{27}H_{37}N_2O_5$ $[M + H]^+$ *m/z* 468.59, found 469.20.

(±)-**22: *trans*-4-(dispiro[adamantane-2,3'-[1,2,4]trioxolane-5',1''-cyclohexan]-3''-yl)phenyl morpholine-4-carboxylate**

To a solution of (±)-**S4** (49 mg, 0.09 mmol, 1.0 equiv) in dichloromethane (2.0 mL), was added triethylamine (39 μL, 0.28 mmol, 3 equiv), followed by morpholine hydrochloride (17 mg, 0.14 mmol, 1.5 equiv). The bright yellow mixture was then allowed to stir at room temperature for 3 d, at which point, the reaction was judged complete by TLC and LCMS. The reaction was then diluted with dichloromethane (15 mL) and subsequently extracted with 1M NaOH (20mL). The organic layer was then washed with brine (20 mL), dried over anhydrous $MgSO_4$, filtered, and concentrated under reduced pressure. The residue was then purified through flash column chromatography (0–60% EtOAc/Hex), to afford (±)-**22** (40 mg, 90%) as a white powder. $^1$H NMR (400 MHz, $CDCl_3$) δ 7.21 (d, *J* = 8.6 Hz, 2H), 7.05 (d, *J* = 8.6 Hz, 2H), 3.80 – 3.73 (m, 4H), 3.63 (d, *J* = 37.4 Hz, 4H), 2.88 – 2.75 (m, 1H), 2.18 – 1.55 (m, 18H), 1.44 – 1.27 (m, 4H). MS (ESI) calculated for $C_{27}H_{35}NO_6$ $[M + Na]^+$ *m/z* 492.58 found 492.20.

(±)-**23: *trans*-4-(dispiro[adamantane-2,3'-[1,2,4]trioxolane-5',1''-cyclohexan]-3''-yl)phenyl thiomorpholine-4-carboxylate 1-oxide**

To a solution of (±)-**S4** (75 mg, 0.14 mmol, 1.0 equiv) in DMF (0.5 mL) cooled to 0 ˚C in an ice bath, was added triethylamine (60 μL, 0.43 mmol, 3.0 equiv), thiomorpholine 1-oxide hydrochloride (34 mg, 0.22 mmol, 1.5 equiv) in DMF (2.0 mL) dropwise via micro syringe. The bright yellow mixture was then allowed to stir for 48 h under argon and allowed to warm up to room temperature, at which point, the reaction was deemed complete by TLC and LCMS. The reaction was then diluted with EtOAc (10 mL) and subsequently extracted with 1M NaOH (15 ml). The organic layer was washed three times, at which point the aqueous layer was colorless and no longer yellow. The organic layer was washed with brine (15 mL), dried over anhydrous $MgSO_4$ and concentrated under reduced pressure. The residue was then purified by silica gel flash column chromatography (0–15% MeOH/$CH_2Cl_2$), the product eluting around 7% MeOH/$CH_2Cl_2$ and yielding (±)-**23** (45 mg, 62%) as a yellow solid. $^1$H NMR (400 MHz, $CDCl_3$) δ 7.23 (d, *J* = 8.5 Hz, 2H), 7.04 (d, 2H), 4.23 (t, 4H), 2.94 (t, 1H), 2.88 – 2.77 (m, 4H), 2.06 – 1.66 (m, 20H), 1.42 – 1.35 (m, 2H). MS (ESI) calculated for $C_{27}H_{35}NO_6S$ $[M + Na]^+$ *m/z* 524.64, found 524.06.

**In vitro antiplasmodial growth inhibition and ring-stage survival assays**

***P. falciparum* culture**
Erythrocytic cultures of *P. falciparum* strain Dd2 (BEI Resources) and CamWT_C580Y (BEI Resources) were maintained using standard methods at 2% hematocrit in RPMI 1640 medium (Invitrogen) supplemented with 0.5% AlbuMAX II (Gibco Life Technologies), 0.1mM hypoxanthine, 30 μg/mL gentamicin, 24mM $NaHCO_3$, and 2 5mM HEPES pH 7.4 (herein referred to as RPMIc) at 37°C in an atmosphere of 5% $O_2$, 5% $CO_2$, and 90% $N_2$. General parasitemia for culture maintenance was monitored via hand count of giemsa-stained iRBC thin smears by light microscopy. Media was changed and fresh RBC was added following a maximum period of 48 hours. Any necessary synchronization steps were completed by incubating ring-heavy iRBC pellets in 5% sorbitol at 37 °C for 15min.

**SYBR Green $IC_{50}$ Assay**
Synchronized ring stage *P. falciparum* parasites maintained in RPMIc (see above) at 37°C under an atmosphere of 5% $O_2$, 5% $CO_2$, and 90% $N_2$ were incubated with test compound at 15000, 4500, 1350, 405, 121.5, and 35.45 nM for 72 hours. Tests were conducted in 96 well plates, each well with 200 μL synchronized ring stage parasite culture at 1% parasitemia and 2% hematocrit. Following this incubation period, parasites and RBCs were lysed by repeated freeze/thaw cycles between -20 °C and room temperature. Following initial lysis, 100 μL SYBR Green buffer was added (20 mM Tris, 5 mM EDTA, 0.02% saponin, 0.08% Triton-X, with 2 μL 10,000 X SYBR Green Nucleic Acid Gel Stain (ThermoFisher) per 10mL buffer) and wells were agitated to mix. Plate fluorescence was then recorded using a Tecan Infinite 200 Pro, Excitation = 254 Emission = 520. Plate data was graphed and $IC_{50}$ values quantified using GraphPad Prism 9.

**YOYO-1 FACS Assay**
Synchronized ring stage *P. falciparum* parasites maintained in RPMIc (see above) at 37°C under an atmosphere of 5% $O_2$, 5% $CO_2$, and 90% $N_2$ were incubated with test compound at 17500, 5250, 1575, 1250, 937.5, 703.1, 472.5, 351.6, 175.8, 141.8, 87.9, and 42.5 nM for 72 hours. Tests were conducted in 96 well plates, each well with 200 μL synchronized ring stage parasite culture at 1% parasitemia and 2% hematocrit. Following this period, RBCs were pelleted via centrifugation, media was aspirated, and RBCs were fixed by the addition of 200 μL 2% formaldehyde in PBS and incubation at 37 °C overnight. PBS-fixed plates were stored at 4 °C until tested by FACS. For testing, a separate plate was populated with YOYO-1 FACS buffer (100 mM $NH_4Cl$, 0.1% Triton-X, PBS 7.4 with freshly added 25 nM YOYO-1) at 190 μL per well. 10 μL of the fixed formaldehyde solution from test plates was transferred from test plates to the prepared FACS plates and allowed to incubate at 4 °C for a minimum of 24 hours. Following this period, wells were analyzed via FACS on an Attune-NxT (ThermoFisher) using the instrument's BL1 fluorescent channel (530/30 excitation/emission) and parasitemia was quantified as a ratio of gated iRBC "hits" versus total RBC count using ranges determined during instrument calibration as well as from internal controls. Values for each well were graphed in GraphPad Prism 9, wherein $IC_{50}$ values were quantified.

**Ring Stage Survival Assay (Laboratory Strains)**
The ring stage survival assay was performed according to protocols established by Witkowski et al *(4)*. Erythrocytic cultures of *P. falciparum* strain W2 (BEI Resources) and CamWT_C580Y (BEI Resources) were maintained using standard methods at 2% hematocrit in RPMI 1640 medium (Invitrogen) supplemented with 0.5% AlbuMAX II (Gibco Life Technologies), 0.1mM hypoxanthine, 30 μg/mL gentamicin, 24mM $NaHCO_3$, and 25 mM HEPES pH 7.4 at 37 °C in an atmosphere of 5% $O_2$, 5% $CO_2$, and 90% $N_2$. Parasite cultures were synchronized by three 5% sorbitol treatments. Parasites at the mature schizont stage were selected using segmentation in 75% percoll to yield a highly synchronous culture which was returned to standard incubation conditions for 3 hours. Cultures were then sorbitol synchronized again to yield exclusively 0-3 hr post-invasion rings (100% by stage). 0-3hr post-invasion rings at 1% parasitemia were exposed to experimental compounds at 700 nM or an equal volume of DMSO for 6 hours. Following exposure, RBCs were pelleted via

centrifugation and washed with RPMIc three total times, with the final resuspension in a fresh 6-well plate to minimize the potential of drug carryover. Cultures were incubated at 37°C for 66 hours. Giemsa-stained thin smears for each group were assessed microscopically to determine postexposure parasitemia. Four experimental replicates were completed for all drug and control groups.

**Growth Inhibition and ex vivo RSA assays for clinical isolates.** Susceptibility of freshly collected isolates of *P. falciparum* to dihydroartemisinin (**DHA**), artefenomel (**1**) and RLA-4735 ((±)-**2**), were determined using the standard microplate growth inhibition assay and a modification of the ring-stage survival assay (RSA) reported by Witkowski *(4)*, with more rigorous washing steps and transfer to unused plates as further detailed below.

Sample collection and processing: Whole blood parasite samples were collected from consenting uncomplicated malaria patients at study sites in northern (Patongo, Kalongo) and eastern (Tororo, Busiu) Uganda, from June 2023 – March 2024. Samples were transported under refrigeration to culture laboratories in Kalongo and Tororo respectively, where they were refrigerated overnight and processed on the following day, averagely < 24 h from time of sample collection. Samples were washed 3X using RPMI wash media to deplete leucocytes and plasma constituents and reconstituted to 50% hematocrit using RPMI complete culture media. Parasitemia was determined from Giemsa-stained thin films by microscopy.

Ex vivo parasite culture assays: For growth inhibition assays, parasite isolates were incubated in serially diluted solutions of DHA, **1** and (±)-**2**, at assay starting parasitemia of 0.2% and 2% hematocrit, in RPMI complete culture medium. Incubation was done at 37 °C in a humid tri-gas (92% $N_2$, 5% $CO_2$, 3% $O_2$) atmosphere for 72 h. Parasite growth was determined at 72 h by incubating parasites in lysis buffer containing 0.02% v/v SYBR Green I dye for 1 h, and detecting fluorescence using a BMG FLUOstar Omega plate reader at ex 485 nm /em 530 nm. $IC_{50}$ values were derived by plotting fluorescence intensity against log drug concentrations and fit to a non-linear curve using a four-parameter Hill equation in GraphPad Prism (version 9.3).

Ex vivo parasite RSA: Parasites were incubated in 700 nM of DHA, **1** and (±)-**2**, and 0.1% DMSO as the vehicle control, at a starting parasitemia of 1% (samples with >1% parasitemia were diluted using complete culture media and donor erythrocytes to 1%) and 2% hematocrit, in 2 mL volumes in a sterile 24-well plate. Cultures were maintained at 37 °C in a humidified tri-gas (92% $N_2$, 5% $CO_2$, 3% $O_2$) atmosphere. At 6 h post-drug exposure, cultures were resuspended and transferred to a 15 mL conical centrifuge tube and washed 3 times by centrifugation (10 min @ 2000 rpm) with 10 mL of RPMI media at 37 °C. Cultures were resuspended into unused wells in 2 mL volumes in complete media and cultured for an additional 66 h. Giemsa-stained thin smears were prepared from drug- and DMSO-treated cultures 66 h post-wash (72 h after the start of the assay). Parasitemia was assessed in the drug-treated cultures by counting parasite-infected erythrocytes from 100 fields containing at least 100 total erythrocytes under 100X light microscopy (Model CX21FS1, Olympus Corp., Tokyo, Japan) by two independent readers. Parasite survival rates were expressed as the parasitemia in the drug-pulsed cultures relative to that in the DMSO controls, at the end of the 72 h assay. All isolates with 72 h control parasitemia of ≥ 0.2% (≥ 20% of initial assay starting parasitemia) was considered valid. RSA survival rates were averaged between the two readers and compared using the Friedman test with Dunn's post-hoc correction, using GraphPad Prism 9.3. Differences were considered significant at $p < 0.05$. RSA percentages less than 10% are generally considered indicative of sensitivity to DHA.

***P. berghei* mouse malaria model**

Female Swiss Webster mice (~20 g body weight) were infected intraperitoneally with $10^6$ *P. berghei*-infected erythrocytes collected from a previously infected mouse. Beginning 1 h after inoculation the mice were treated by oral gavage for 1 or 2 days as indicated with 100 μL of solution of test compound formulated in 10% DMSO, 40% (20% 2-hydroxypropyl-beta-cyclodextrin solution in water), and 50% PEG400. There were five mice in each test arm. Infections were monitored by daily microscopic evaluation of Giemsa-stained blood smears starting on day seven. Parasitemia was determined by counting the number of infected erythrocytes per 1000 erythrocytes. Body weight was measured over the course of the treatment. Mice were euthanized when parasitemia exceeded 50% or when weight loss of more than 15% occurred. Parasitemia, animal survival, and morbidity were closely monitored for 30 days postinfection, when experiments were terminated.

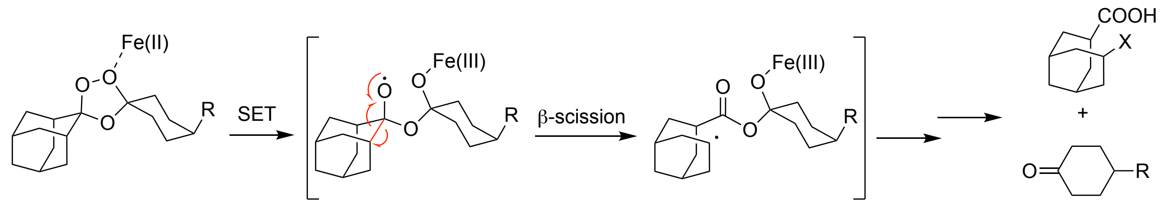


**Fig. S1.** Canonical mechanism of 1,2,4-trioxolane activation via inner-sphere coordination and Fenton-type single-electron transfer reaction with ferrous iron, followed by β-scission to generate carbon-centered radical species that react with parasite macromolecules (X), ultimately resulting in parasite death.

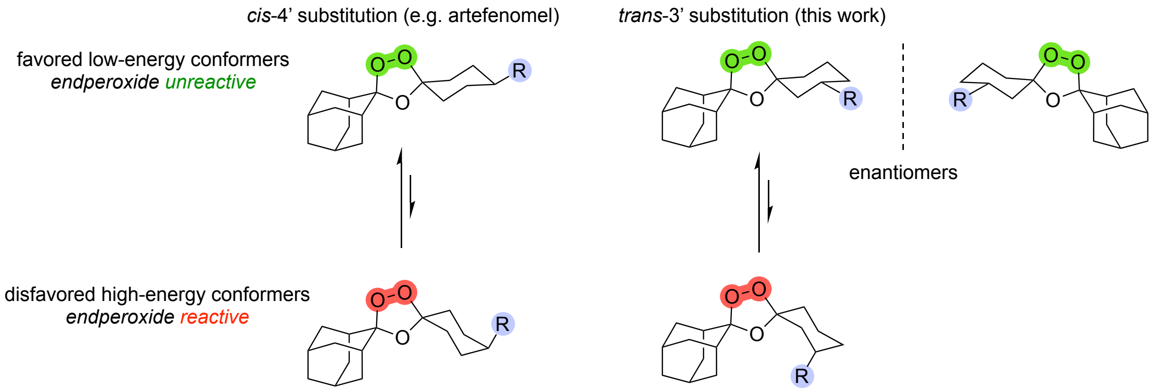


**Fig. S2.** Conformational dynamics underlying 1,2,4-trioxolane activation and antiplasmodial action. Classical trioxolane antimalarials like arterolane and artefenomel (**1**) bear cis-4’ substitution that stabilizes an iron-unreactive conformer (top, in green). Similarly, *trans*-3’ substitution (as in compounds **2**-**23** studied herein) stabilizes the same unreactive ground state conformation. Substitution at the 3’ position introduces molecular asymmetry and stereoisomerism, as in the enantiomeric forms (*R,R*)-**2** and (*S,S*)-**2**. Chair–chair interconversion exposes the endoperoxide bond (bottom in red) for reaction with ferrous iron sources in the parasite, ultimately leading to alkylation of parasite proteins and cell death.

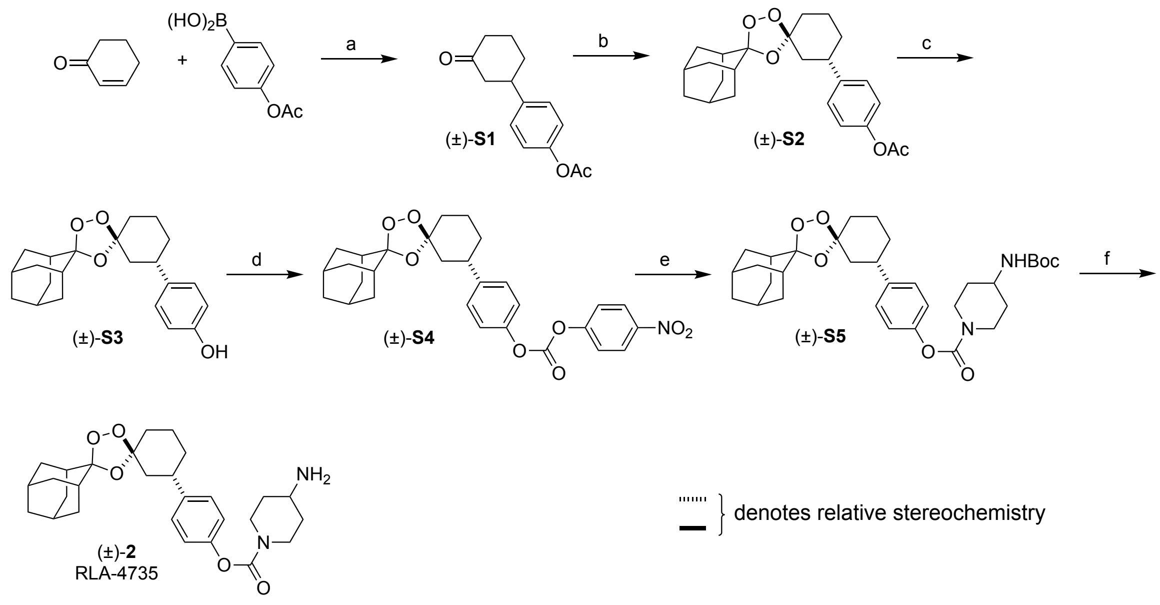


**Fig. S3.** Synthesis of racemic (±)-**2** (RLA-4735) comprising both (*R,R*) and (*S,S*) forms. Conditions: (a) 5 mol% Pd(acac)$_2$, 20 mol% Cu(BF$_4$)$_2$•H2O), 5 mol% dppben, DME, rt, 91%. (b) adamantan-2-one *O*-methyl oxime, CCl$_4$, O$_3$, 0°C, 86% (~7:1 *trans:cis* dr). (c) 15% aq. KOH, THF/MeOH (1:2), 50 °C, 2.5h, 93%. (d) bis-(4-nitrophenyl)carbonate, DIEA, CH$_2$Cl$_2$, rt. (e) *tert*-butylpiperidin-4-ylcarbamate, 2h, rt, 29% over two steps). (f) AcCl, MeOH, rt, 65%.

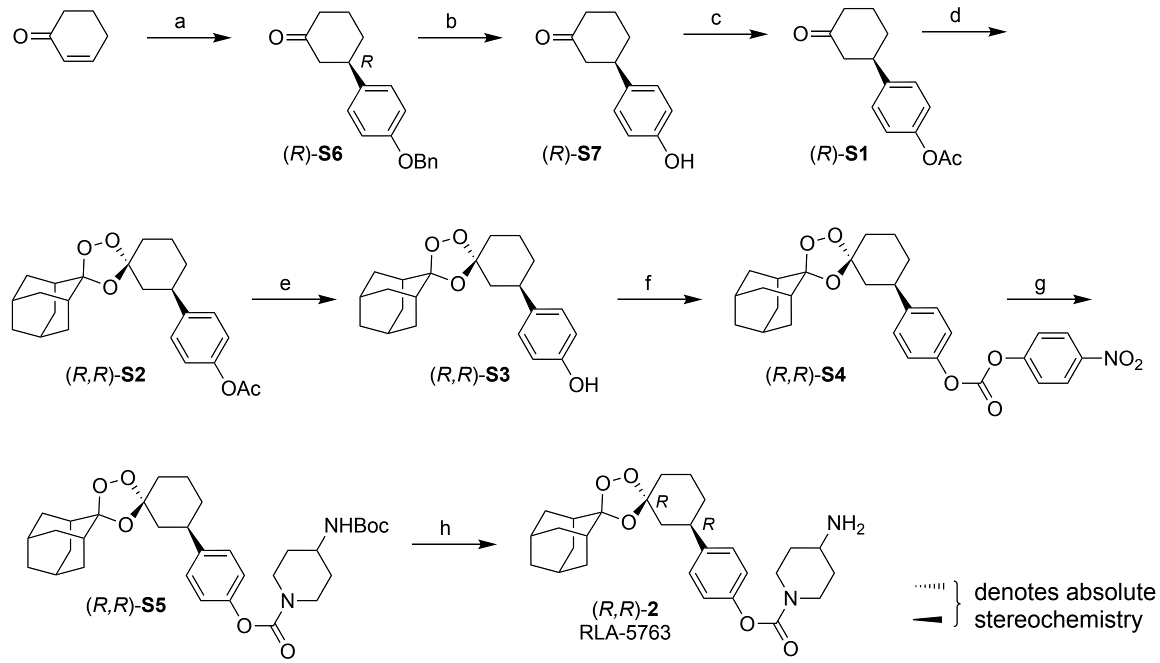


**Fig. S4.** Synthesis of (*R,R*)-**2** (RLA-5763). Conditions: Conditions: (a) 4-(benzyloxy)phenylboronic acid, 4 mol% acetylacetonatobis(ethylene)rhodium(I), 10 mol% *R-BINAP, aq KOH, dioxane, 100 °C, 16 h, 54%. (b) $H_2$, Pd/C 10%, EtOAc, 50°C, 92%. (c) $Ac_2O$, $Et_3N$, $CH_2Cl_2$, 0°C to rt, 95%. (d) 2.5 eq. adamantan-2-one-*O*-methyl oxime, $CCl_4$, $O_3$, 0°C, 67%. (e) LiOH, THF/MeOH, rt, 16 h, 85%. (f) 4-nitrophenyl chloroformate, DIEA, 0.2 eq DMAP, $CH_2Cl_2$, 0°C to rt, 26%. (g) *tert*-butylpiperidin-4-ylcarbamate, DIEA, 0.2 eq DMAP, DMF 2h, rt, 68%. (h) AcCl, MeOH, 0°C to rt, 36%.

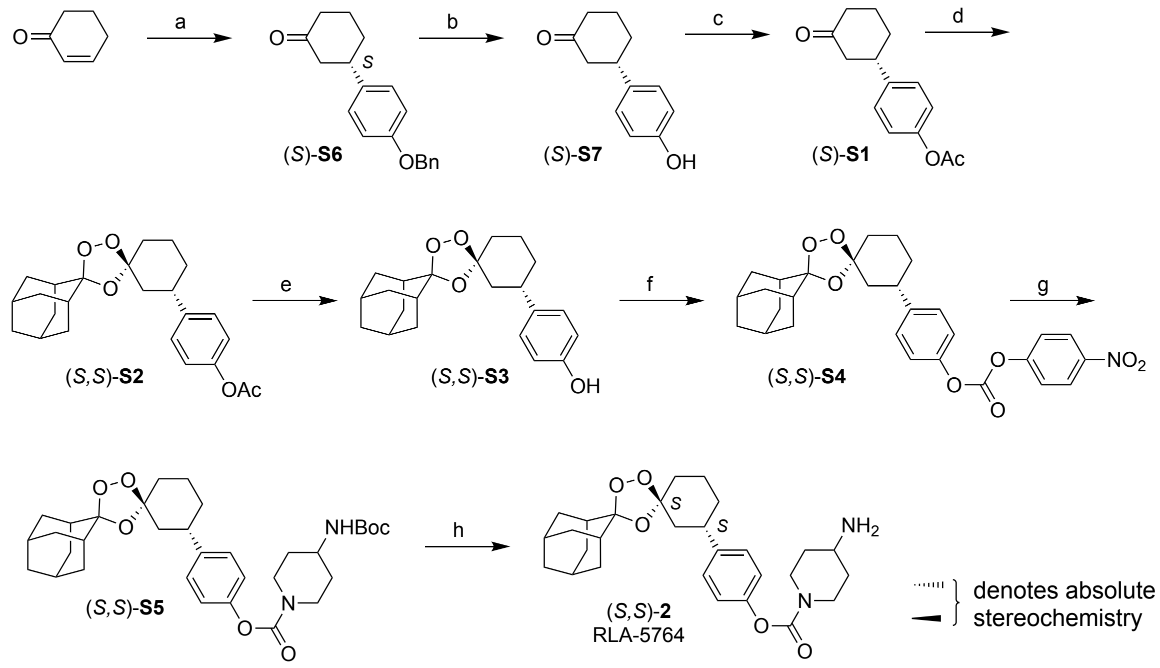


**Fig. S5.** Synthesis of (*S,S*)-**2** (RLA-5764). Conditions: Conditions: (a) 4-benzyloxyphenylboronic acid, 4 mol% acetylacetonatobis(ethylene)rhodium(**I**), 10 mol% *S-BINAP, aq KOH, dioxane, 100 $^{o}$C, 16 h, 33%. (b) $H_2$, Pd/C 10%, EtOAc, 50$^{o}$C, 87%. (c) $Ac_2O$, $Et_3N$, $CH_2Cl_2$, 0$^{o}$C to rt, 95%. (d) 2.5 eq. adamantan-2-one-*O*-methyl oxime, $CCl_4$, $O_3$, 0$^{o}$C, 42%. (e) LiOH, THF/MeOH, rt, 16 h, 89%. (f) 4-nitrophenyl chloroformate, DIEA, 0.2 eq DMAP, $CH_2Cl_2$, 0$^{o}$C to rt, 41%. (g) *tert*-butylpiperidin-4-ylcarbamate, DIEA, 0.2 eq DMAP, DMF 2h, rt, 88%. (h) AcCl, MeOH, 0$^{o}$C to rt, 61%.

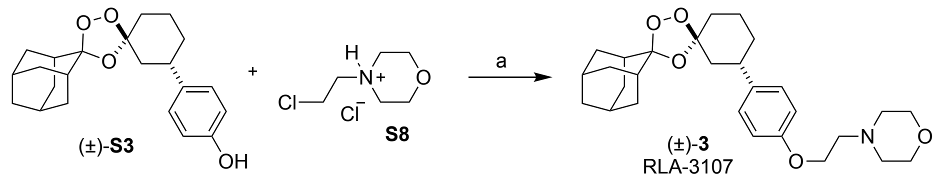


**Fig. S6.** Synthesis of (±)-**3** (RLA-3107). Conditions (a) NaOH, $(n\text{-Bu})_4N(HSO_4)$ 0.2 eq, $CH_3CN$, 55 °C, 15 h, 62%.

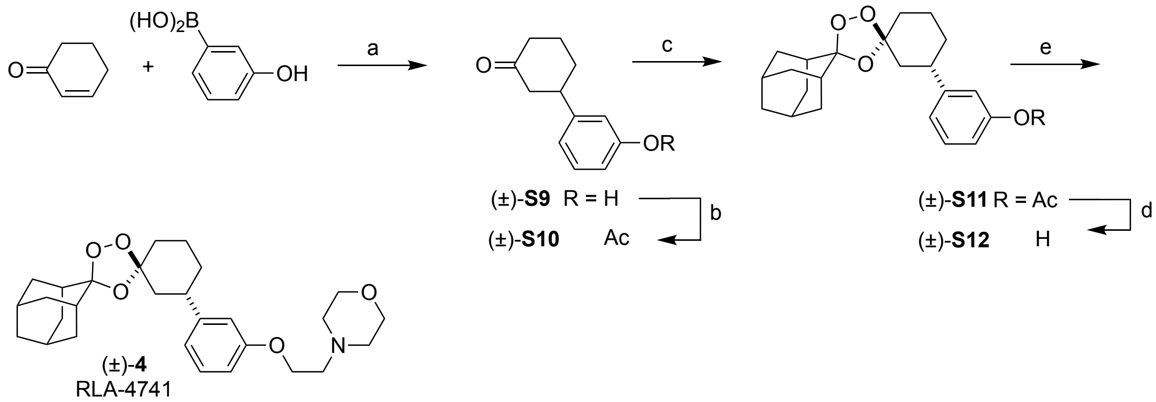


**Fig. S7.** Synthesis of (±)-**4** (RLA-4741). Conditions: (a) 5 mol% Pd(acac)$_2$, 5 mol% dppbz, 20 mol% Cu(BF$_4$)$_2$•H$_2$O, DME, rt, 20 h, 73%. (b) Ac$_2$O, DMAP, CH$_2$Cl$_2$, rt, 78%. (c) 2.5 eq. adamantan-2-one-*O*-methyl oxime, CCl$_4$, O$_3$, 0ºC, 84%, ~7:1 *trans:cis* dr. (d) KOH, THF/MeOH, 50 ºC, 1.5 h, 85%. (e) 2.0 eq **S8**, 3.6 eq NaOH, 0.3 eq (n-Bu)$_4$N(HSO$_4$) 0.2 eq, CH$_3$CN, 55 ºC, 15 h, 61%, pure *trans* diastereomer.

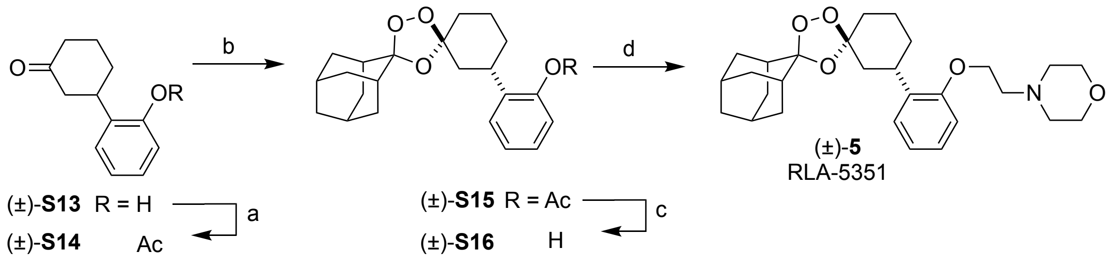


**Fig. S8.** Synthesis of (±)-**5** (RLA-5351). Conditions: (a) $Ac_2O$, 5 eq pyridine, $CH_2Cl_2$, rt, 95%. (b) 2.5 eq. adamantan-2-one-*O*-methyl oxime, $CCl_4$, $O_3$, 0ºC. (c) KOH, THF/MeOH, 50 ºC, 1.5 h, 70% over two steps. (d) 2.0 eq **S8**, 3.6 eq NaOH, 0.3 eq $(n\text{-}Bu)_4N(HSO_4)$ 0.2 eq, $CH_3CN$, 55 ºC, 15 h, 61%.

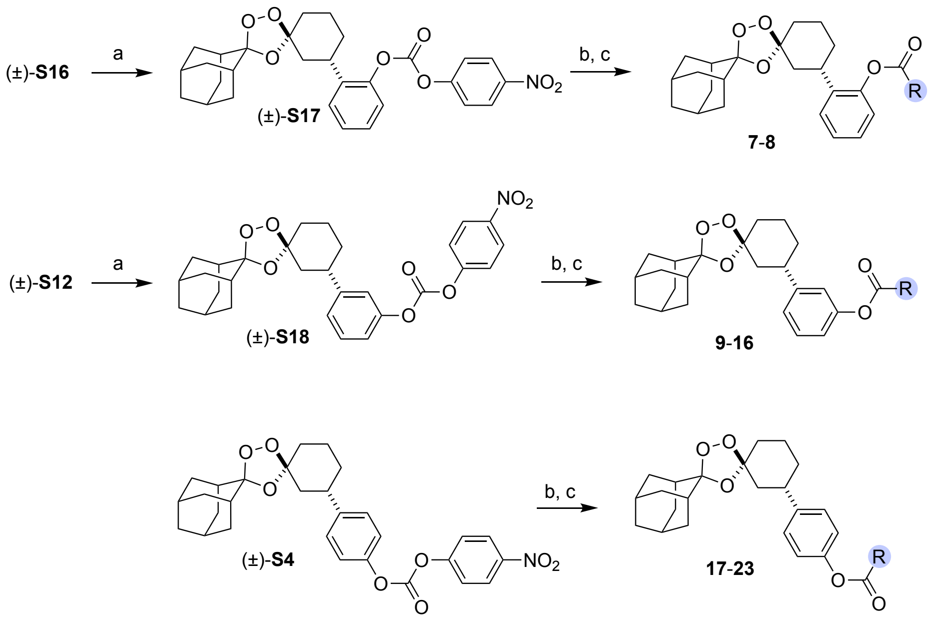


**Fig. S9.** General synthetic schemes for the synthesis of aryl carbamate analogs **7-23** starting from key phenolic intermediates (±)-**S3**, (±)-**S12**, and (±)-**S16**. Conditions (a) 2-3 eq. *p*-($NO_2$)PhOC(=O)Cl, 0.2 eq. DMAP, 3 eq. DIEA, $CH_2Cl_2$, 0 ºC to rt, 77%. (b) $R^1(R^2)NH$, $Et_3N$, $CH_2Cl_2$. (c) excess AcCl, MeOH (only when final Boc deprotection of side chain is required). Complete structures of **7-23** are provided in Figure S7 above.

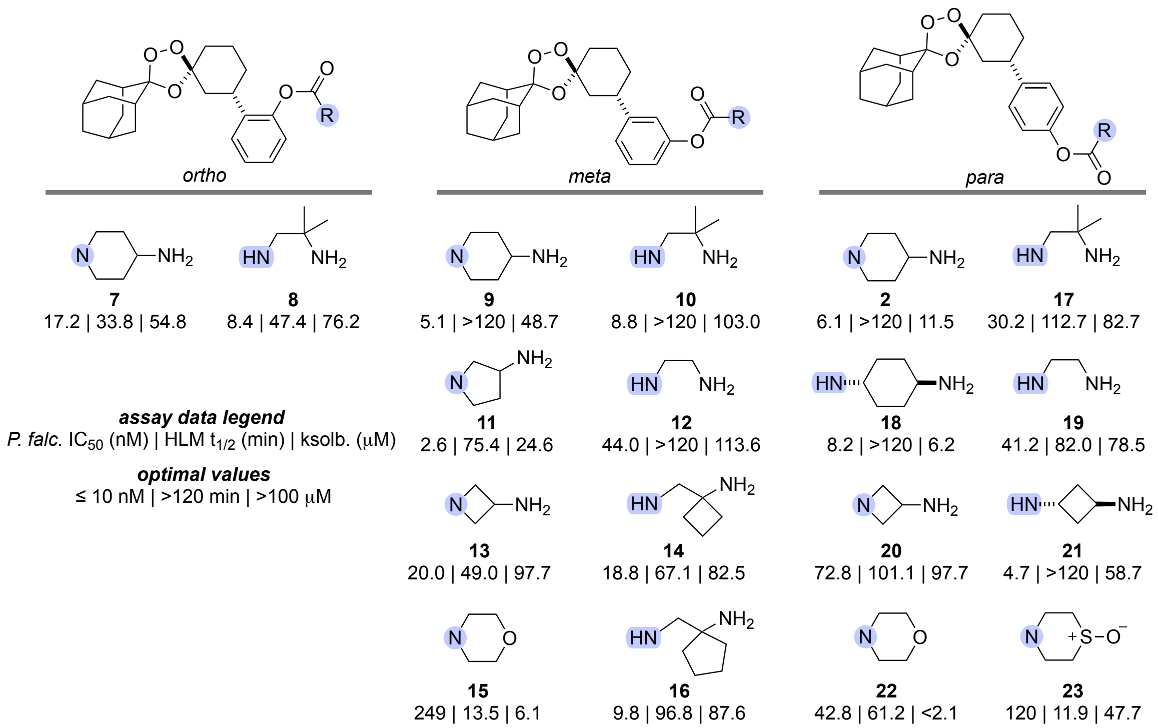


**Fig. S10.** Summary of Structure-Activity Relationships (SAR) for ortho, meta, and para-substituted analogs with aryl carbamate substitution. All compounds are pure trans diastereomers in racemic form. *P. falciparum* $IC_{50}$ values are for W2 strain, chloroquine-resistant parasites. Human liver microsome (HLM) stability $t_{1/2}$ values were generated at Quintara Biosciences (South San Francisco, CA). Kinetic solubility (ksolb) values were generated at Analiza, Inc., Cleveland, OH.

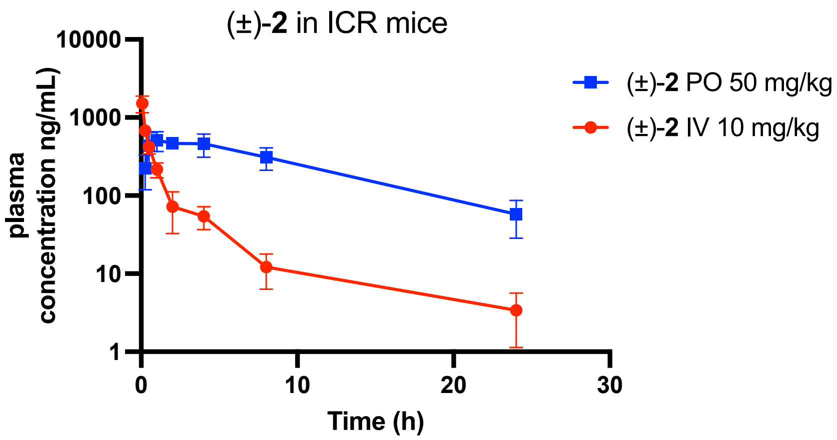


**Figure S11.** Pharmacokinetic profile of (±)-**2** following IV or PO administration in ICR mice at the indicated doses.

A.

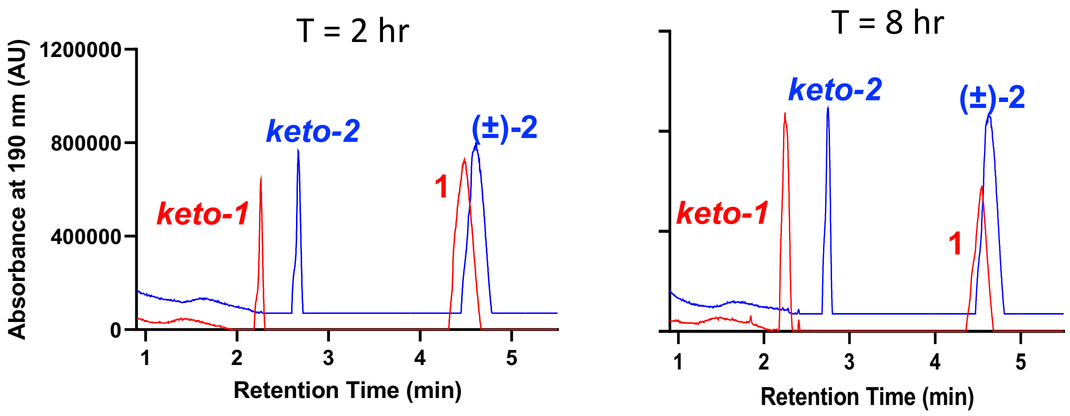


B.

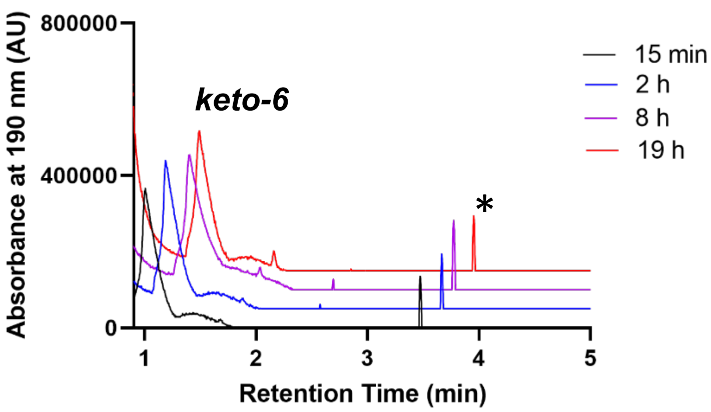


C.

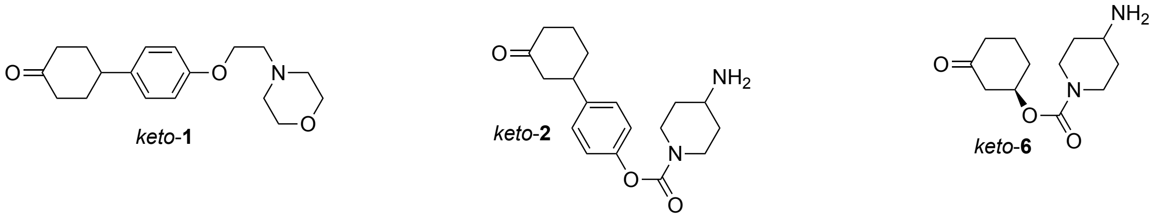


**Fig. S12.** Representative chromatographic traces from the incubations of **1**, (±)-**2**, and (*R,R*)-**6** with ferrous iron solutions leading to fragmentation of the trioxolane ring to form the expected ketone products, confirmed by LC/MS analysis (see Fig. S2 for mechanistic details). Aryl substituted analogs **1** and (±)-**2** (**A**) react much more slowly than comparator (*R,R*)-**6**, which is fully consumed by 15 min (**B**). Structures of respective ketone products are shown at bottom (**C**). Peak labelled with asterisk in **B** represents a minor, uncharacterized reaction product. Reaction conditions: 50 mM ferrous ammonium sulfate [$Fe(NH_4)_2(SO_4)_2{\bullet}6H_2O$], citrate buffer, 37°C.

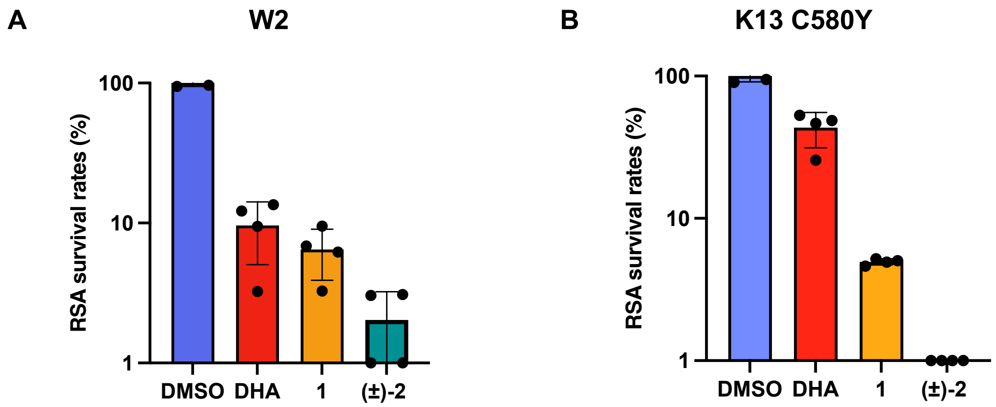


**Fig. S13.** In vitro RSA survival rates for laboratory strains. (A) in vitro RSA values determined for test compounds against susceptible W2 strain *P. falciparum* and (B) DHA partial resistant *P. falciparum* K13 C580Y mutant parasites.

**Table S1.** Pharmacokinetic parameters from PK study of **1**, (±)-**2**, and (*R,R*)-**6** male CD-1 mice (n = 6 per group) following a single dose of 50 mg/kg by oral gavage.

| compound | $T_{max}$ (hr) | $C_{max}$ ± SEM (ng/mL) | $AUC_{last}$ ± SEM (hr•ng/mL) |
|---|---|---|---|
| **1** | 2 | 5660 ± 732 | 52600 ± 4810 |
| (±)-**2** | 2 | 1190 ± 58 | 16400 ± 1640 |
| (*R,R*)-**6** | 2 | 631 ± 205 | 3430 ± 772 |

**Table S2.** In vitro ADME Properties of (*S,S*)-**2** and relevant controls.[a]

| metabolic stability | | | | | | |
|---|---|---|---|---|---|---|
| | microsome $t_{1/2}$ by species (min) | | | hepatocyte $t_{1/2}$ by species (min) | | |
| | rat | dog | human | rat | dog | human |
| (*S,S*)-**2** | >240 | >240 | >240 | 276.3 | >480 | >480 |
| *verapamil* | *5.1* | *9.1* | *8.0* | | | |
| *midazolam* | | | | *20.38* | *36.94* | *92.63* |

| plasma stability by species | | |
|---|---|---|
| | mouse plasma $t_{1/2}$ (min) | human plasma $t_{1/2}$ (min) |
| (*S,S*)-**2** | 251.4 ± 27.7 | >480 |
| *propantheline* | 31.6 ± 1.1 | 14.9 ± 1.6 |

| binding and distribution | | | | | | | |
|---|---|---|---|---|---|---|---|
| | plasma protein binding by species (%) | | | hu heps. binding[b] (%) | hu mics. binding[c] (%) | Bl/Pl ratio | $K_{RBC/PL}$ |
| | Rat | Dog | Human | | | | |
| (*S,S*)-**2** | >99.98 | >99.98 | >99.98 | 99.91 | 99.94 | 1.61 | 3.55 |
| *propranolol* | *90.65* | *86.74* | *82.75* | | | | |
| *warfarin* | *99.38* | *94.71* | *99.20* | | | | |
| *chlorpromazine* | | | | *99.01* | *92.21* | | |
| *chloroquine* | | | | | | *2.04* | *2.51* |

| solubility and permeability | | | | | | |
|---|---|---|---|---|---|---|
| | thermodynamic solubility (μg/mL) | | Caco-2 $P_{app}$ A-B in pH 7.4 hu plasma ($x10^{-6}$ cm/s) | Caco-2 $P_{app}$ B-A in pH 7.4 hu plasma ($x10^{-6}$ cm/s) | Caco-2[b] $P_{app}$ A-B in pH 7.4 hu plasma ($x10^{-6}$ cm/s) | Caco-2[b] $P_{app}$ A-B in pH 7.4 hu plasma ($x10^{-6}$ cm/s) |
| | FaSSIF (pH 6.5) | FeSSIF (pH 5.0) | | | | |
| (*S,S*)-**2** | 2.3 | 4525.5 | 0.6 | 0.2 | 1074 | 284 |
| *digoxin* | | | *1.4* | *4.2* | | |
| *propranolol* | | | *10.5* | *4.2* | *65* | *26* |

[a]Data reported in this table was generated by Quintara Biosciences, South San Francisco, CA.
[b]Corrected permeability values are adjusted to account for the unbound fraction of drug from human plasma protein binding assay using the approach reported by Charman and co-workers ***(5)***.

**Table S3**. Pharmacokinetic parameters for (±)-**2** following IV (10 mg/kg) and PO (50 mg/kg) doses in male ICR mice (n = 3 per group).

| IV | | | | PO | | | |
|---|---|---|---|---|---|---|---|
| Parameter | units | mean | SD | Parameter | units | mean | SD |
| $t_{1/2}$ | hr | 3.42 | 1.71 | $t_{1/2}$ | hr | 7.12 | 2.65 |
| $T_{max}$ | hr | 0.083 | NA | $T_{max}$ | hr | 2.17 | 1.76 |
| $C_{max}$ | ng/mL | 1520 | 365 | $C_{max}$ | ng/mL | 565 | 133 |
| $C_0$ | ng/mL | 2270 | 650 | $AUC_{0\text{-}t}$ | ng/mL*hr | 6250 | 1490 |
| $AUC_{0\text{-}t}$ | ng/mL*hr | 1130 | 194 | $AUC_{0\text{-}inf}$ obs | ng/mL*hr | 6920 | 1380 |
| $AUC_{0\text{-}inf}$ obs | ng/mL*hr | 1150 | 185 | $MRT_{0\text{-}inf}$ obs | hr | 9.830 | 3 |
| $MRT_{0\text{-}inf}$ obs | hr | 2.53 | 0.99 | F | % | 120 | 23.9 |
| CL, obs | mL/min/kg | 147 | 22.7 | | | | |
| CL, obs | L/hr/kg | 8.8 | 1.36 | | | | |
| Vss, obs | L/kg | 22.7 | 11.7 | | | | |

Data was processed by Phoenix WinNonlin 8.3, and samples below limit of quantitation excluded in the calculation of PK parameters and mean concentration. CL, obs values are provided in both mL/min/kg and L/hr/kg. Liver blood flow (mouse) = 7.2 L/hr/kg

**Table S4**. Pharmacokinetic parameters for (±)-**2** following IV (10 mg/kg) and PO (50 mg/kg) doses in male SD rats (n = 3 per group).

| IV | | | | PO | | | |
|---|---|---|---|---|---|---|---|
| Parameter | units | mean | SD | Parameter | units | mean | SD |
| $t_{1/2}$ | hr | 10.2 | 2.41 | $t_{1/2}$ | hr | BLOQ | NA |
| $T_{max}$ | hr | 0.083 | NA | $T_{max}$ | hr | 8.000 | NA |
| $C_{max}$ | ng/mL | 1660 | 69.2 | $C_{max}$ | ng/mL | 220 | 41.8 |
| $C_0$ | ng/mL | 2210 | 50.9 | $AUC_{0\text{-}t}$ | ng/mL*hr | 4010 | 640 |
| $AUC_{0\text{-}t}$ | ng/mL*hr | 2500 | 101 | $AUC_{0\text{-}inf}$ obs | ng/mL*hr | BLOQ | NA |
| $AUC_{0\text{-}inf}$ obs | ng/mL*hr | 2910 | 148 | $MRT_{0\text{-}inf}$ obs | hr | BLOQ | NA |
| $MRT_{0\text{-}inf}$ obs | hr | 10.1 | 2.50 | F | % | 27.6 | 4.4 |
| CL, obs | L/hr/kg | 3.44 | 0.18 | | | | |
| Vss, obs | L/kg | 34.4 | 7.32 | | | | |

Data was processed by Phoenix WinNonlin 8.3, and samples below limit of quantitation excluded in the calculation of PK parameters and mean concentration. Liver blood flow (rat) = 4.1 L/hr/kg

**Table S5**. Pharmacokinetic parameters for (*S,S*)-**2** following IV (3 mg/kg) and PO (10 mg/kg) doses in male SD rats (n = 3 per group).

| IV | | | | PO | | | |
|---|---|---|---|---|---|---|---|
| Parameter | units | mean | SD | Parameter | units | mean | SD |
| CL | L/hr/kg | 5.36 | 0.543 | $T_{max}$ | hr | 6.67 | 2.31 |
| Vss | L/kg | 17.6 | 2.65 | $C_{max}$ | ng/mL | 82.2 | 12.8 |
| $T_{1/2}$ | hr | 2.94 | 0.449 | $T_{1/2}$ | hr | 8.93† | – |
| $AUC_{last}$ | ng/mL*hr | 498 | 43.7 | $AUC_{last}$ | ng/mL*hr | 1255 | 129 |
| $AUC_{INF}$ | ng/mL*hr | 564 | 54.4 | $AUC_{INF}$ | ng/mL*hr | 1456† | – |
| $MRT_{INF}$ | hr | 3.30 | 0.592 | F | % | 75.6 | 7.75 |
| CL, renal | L/hr/kg | 0.816 | 0.055 | | | | |
| f, urine | % | 0.878 | 0.506 | | | | |
| $CL_{blood}$ | L/hr/kg | 3.33 | 0.34 | | | | |
| $CL_{blood}$ | mL/min/kg | 55.3 | 5.6 | | | | |
| $Vss_{blood}$ | L/kg | 10.9 | 1.65 | | | | |

†Lower limit estimate from one of three rats in study. Timepoints beyond 24 hr would be required to calculate true half-life and $AUC_{INF}$. Liver blood flow (rat) = 4.1 L/hr/kg; kidney blood flow (rat) = 2.2 L/hr/kg. $CL_{blood}$ and $Vss_{blood}$ were calculated by first converting plasma concentrations to blood concentrations using the blood:plasma ratio (1.61). The $CL_{blood}$ value is provided in both L/hr/kg and mL/min/kg.

**Table S6**. Sample characteristics and drug susceptibility of Ugandan *P. falciparum* isolates tested in 2023 to DHA, **1**, and (±)-**2**.

| **Participant and sample characteristics** | |
|---|---|
| sample source | Northern and eastern Uganda |
| study period | June 2023 - March 2024 |
| number of samples collected, n | 166 |
| study participant age, years, (median IQR) | 8.0 (4.0 - 13.0) |
| sample parasitemia, % (median IQR) | 1.2 (0.6 - 2.5) |
| sample parasitemia, %, range | 0.2-23.7 |
| $IC_{50}$ assay starting parasitemia, % | 0.2 |
| RSA starting parasitemia, % | 1.0 |

| **Results and parameters** | | | |
|---|---|---|---|
| drug/compound | **DHA** | **1** | (±)-**2** |
| Number of $IC_{50}$ assays done, n | 166 | 166 | 166 |
| Number of successful $IC_{50}$ assays, n | 153 | 153 | 151 |
| $IC_{50}$ values, nM, (median IQR) | **3.8** (2.4-5.3) | **3.0** (1.6-4.5) | **3.6** (2.2-5.2) |
| Number of RSAs performed, n | 78 | 78 | 78 |
| RSAs with 2 reads, n | 60 | 60 | 60 |
| RSAs with valid reads, n | 42 | 42 | 42 |
| RSA survival rate, % (median IQR) | **5.3** (2.2-11.3) | **0.0** (0.0-0.0) | **0.0** (0.0-0.0) |

### Supporting Materials References


1. B. R. Blank, J. Gut, P. J. Rosenthal, A. R. Renslo, Artefenomel Regioisomer RLA-3107 Is a Promising Lead for the Discovery of Next-Generation Endoperoxide Antimalarials., *ACS Med. Chem. Lett.* **14**, 493–498 (2023).

2. B. R. Blank, R. L. Gonciarz, P. Talukder, J. Gut, J. Legac, P. J. Rosenthal, A. R. Renslo, Antimalarial Trioxolanes with Superior Drug-Like Properties and In Vivo Efficacy., *ACS Infect. Dis.* **6**, 1827–1835 (2020).

3. C. M. Woodley, G. L. Nixon, N. Basilico, S. Parapini, W. D. Hong, S. A. Ward, G. A. Biagini, S. C. Leung, D. Taramelli, K. Onuma, T. Hasebe, P. M. O'Neill, Enantioselective synthesis and profiling of potent, nonlinear analogues of antimalarial tetraoxanes E209 and N205., *ACS Med. Chem. Lett.* **12**, 1077–1085 (2021).

4. B. Witkowski, D. Menard, C. Amaratunga, R. M. Fairhurst, Ring-stage Survival Assays (RSA) to evaluate the in-vitro and ex-vivo susceptibility of Plasmodium falciparum to artemisinins. , *Natl. Inst. Health Proced. RSAv1* , 1–16 (2013).

5. K. Katneni, T. Pham, J. Saunders, G. Chen, R. Patil, K. L. White, N. Abla, F. C. K. Chiu, D. M. Shackleford, S. A. Charman, Using Human Plasma as an Assay Medium in Caco-2 Studies Improves Mass Balance for Lipophilic Compounds., *Pharm. Res.* **35**, 210 (2018).
